# A spatiotemporal cancer cell trajectory underlies glioblastoma heterogeneity

**DOI:** 10.1101/2025.05.13.653495

**Authors:** Grant de Jong, Fani Memi, Tannia Gracia, Olga Lazareva, Oliver Gould, Alexander Aivazidis, Manas Dave, Qianqian Zhang, Melanie Jensen, Ahmet Sureyya Rifaioglu, Joao D. Barros-Silva, Sabine Eckert, Di Zhou, Yvette Wood, Elizabeth Tuck, Sezgin Er, Henry Marshall, Kenny Roberts, Andrew L. Trinh, Shreya Rai, Tyler Shaw, Agnes Oszlanczi, Hayden Powell, Robert Petryszak, Zoi Katsirea, Irfan Mamun, Ilaria Mulas, Annelies Quaegebeur, Mayen Briggs, Stanislaw Makarchuk, Jessica Cox, Jimmy Tsz Hang Lee, Laura Rueda, Manu Saraswat, Harry Bulstrode, Adam Young, Minal Patel, Tarryn Porter, Elena Prigmore, Moritz Mall, Julio Saez-Rodriguez, James Briscoe, David H. Rowitch, Richard Mair, Sam Behjati, Oliver Stegle, Omer Ali Bayraktar

**Author notes:** **Correspondence:** (S.B.), (O.S.) and (O.A.B.). equal contribution (Grant de Jong, Fani Memi, Tannia Gracia, Olga Lazareva).

## Abstract

Cancer cells display highly heterogeneous and plastic states in glioblastoma, an incurable brain tumour. However, how these malignant states arise and whether they follow defined cellular trajectories across tumours is poorly understood. Here, we generated a deep single cell and spatial multi-omic atlas of human glioblastoma that pairs transcriptomic, epigenomic and genomic profiling of 12 tumours across multiple regions. We identify that glioblastoma heterogeneity is driven by spatially-patterned transitions of cancer cells from developmental-like states towards those defined by a glial injury response and hypoxia. This cellular trajectory regionalises tumours into distinct tissue niches and manifests in a molecularly conserved manner across tumours as well as genetically distinct tumour subclones. Moreover, using a new deep learning framework to map cancer cell states jointly with clones *in situ*, we show that tumour subclones are finely spatially intermixed through glioblastoma tissue niches. Finally, we show that this cancer cell trajectory is intimately linked to myeloid heterogeneity and unfolds across regionalised myeloid signalling environments. Our findings define a stereotyped trajectory of cancer cells in glioblastoma and unify glioblastoma tumour heterogeneity into a tractable cellular and tissue framework.

## Main

The extensive tumour heterogeneity of glioblastoma (GB), an incurable adult brain cancer, presents a major obstacle to treatment. Nearly two decades of molecular profiling has found that malignant cells exist across a spectrum of cellular states within each tumour, resembling developmental neural progenitors as well as mesenchymal-like and injury response states^1–5^. These states are suggested to vary in abundance across tumours and stratify GBs into subtypes^1,2^. Furthermore, patient derived xenografts have shown that malignant cells can transition across cellular states, indicating their plasticity^2,6^. However, despite these extensive profiling and modelling efforts, we still do not understand how cancer cells diversify into distinct states and give rise to GB heterogeneity. In particular, how malignant cells undergo state transitions and whether they follow stereotyped or variable cellular trajectories across GB tumours is unknown. A cancer cell trajectory that is conserved and tractable across GBs would present a highly attractive therapeutic target.

Single cell, bulk and spatial transcriptomics have revealed stereotypical aspects of GBs. Multiple malignant cell states can be detected within each tumour^2,5,7^ and their single cell transcriptional profiles can be mapped onto an axis between dev-like and injury response-associated gene expression programs^4,8^. Furthermore, these states show a recurrent spatial organisation across tumours^5,9^. Yet, these observations have not been linked together to resolve malignant cellular trajectories in GB, in part due to the lack of multi-modal profiling efforts that can integrate insights from various cell and tissue atlassing technologies.

To chart the cellular trajectories of malignant cells in GB, it is necessary to account for their genetic heterogeneity. Whole genome and exome sequencing has identified multiple phylogenetically distinct subclonal lineages within individual GBs^5,7^. Whereas single cell and spatial transcriptomics suggest diverse malignant cell states can arise within each subclone^2,9^, these efforts inferred subclones from transcriptomic data and do not incorporate “ground truth” somatic cancer mutations identified at the genomic DNA level. Hence, the subclonal origins of diverse malignant cell states and whether GB intratumoural genetic heterogeneity can alter these states is not well understood.

Beyond malignant cells, GB heterogeneity extends to the tumour microenvironment (TME). Myeloid cells are the most abundant TME cell type in GB^2,10^, underlying a highly immunosuppressive environment linked to poor clinical outcomes^10–13^. While macrophage-derived signals were previously shown to alter GB malignant cell states^14^, recent studies have identified highly diverse myeloid subtypes in tumours^15–17^. How the heterogeneity of malignant and myeloid cells relate to one another in GBs and how distinct myeloid subtypes influence the pathobiology of diverse malignant cells states is not well understood.

Here, we mapped the cellular, clonal and tissue architecture of GB heterogeneity by deep multi-region profiling that integrates single cell multi-omics, spatial transcriptomics and spatial whole genome sequencing. We identified a malignant cellular trajectory that is spatially patterned in tumours, that is conserved across tumours and spatially-intermixed subclones within each tumour, and that is accommodated by a regionalised myeloid signalling environment. Hence, we find that a stereotyped trajectory of cancer cells gives rise to GB heterogeneity and we provide a cell and tissue framework that unifies GB tumour heterogeneity.

## Results

### Deep spatial multi-omic profiling of GB

To chart the cellular and tissue organisation of GB, we generated a single cell and spatial multi-omic atlas of 12 isocitrate dehydrogenase wildtype (IDH-wt) primary GB tumours (**Fig. 1A, Table S1**). We combined three major approaches to comprehensively profile GB heterogeneity, extending previous surveys^2,5,8,9,18^.

**Figure 1:**
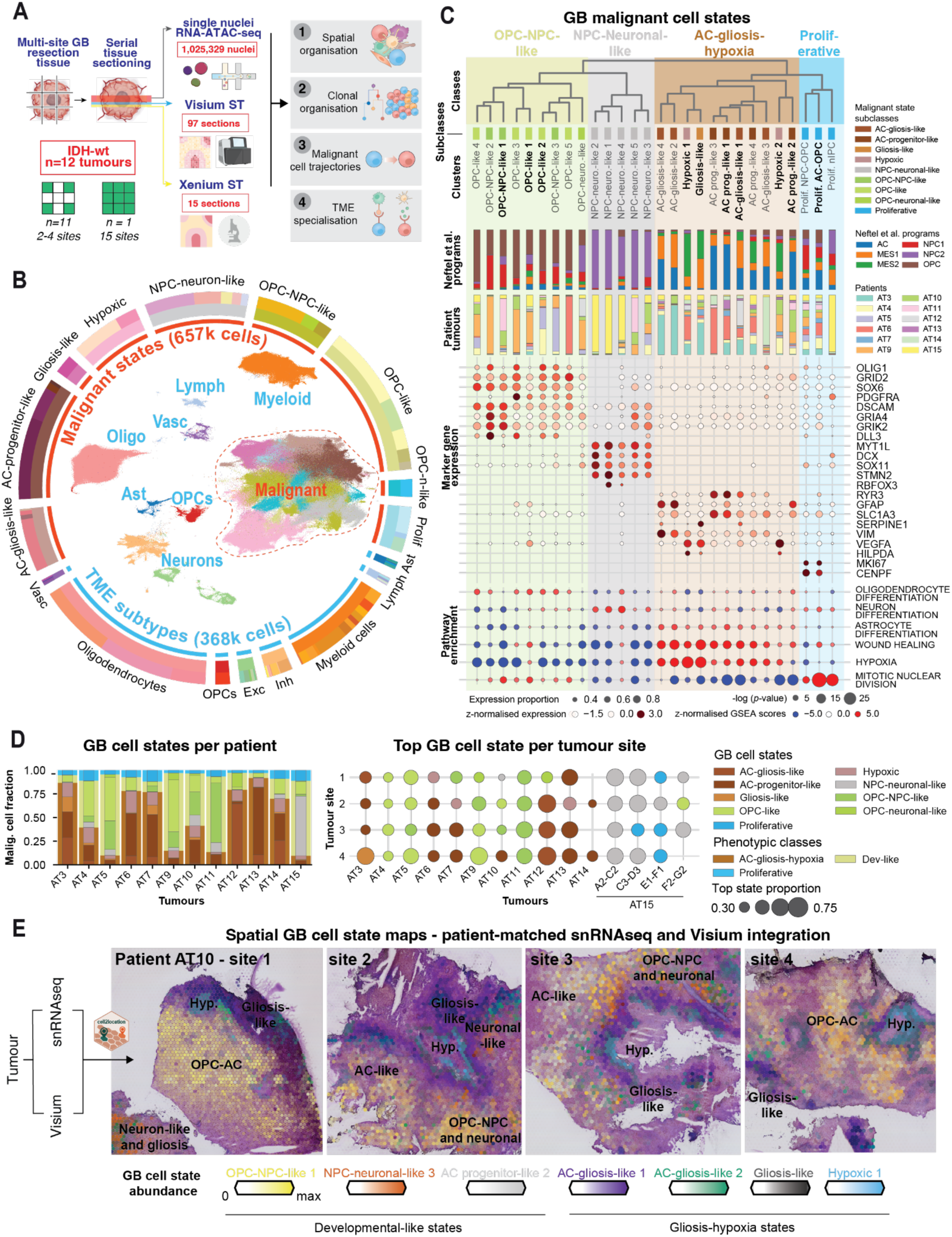
Spatially resolved single cell multiomic atlas of GB tumour heterogeneity. **A)** Schematic illustration of experimental design and primary aims of this study. **B)** Overview of the cellular composition of our GBM-space atlas. UMAP visualisation of the snRNA-seq profiles of malignant and TME cells outlining the major constituent populations, and a ring plot illustrating the relative distribution of cells at three levels of granularity: malignant (657,275 cells) versus TME (368,054 cells) status, coarse-level populations (text labels around the outer ring), and fine grained cell states. **C)** Dendrogram depicting transcriptomic clustering of malignant cell states (top) and their biological characteristics, including their hierarchical annotation, distribution of top Neftel et al. expression signatures, distribution of cell states across patient tumours, marker gene expression, and significantly enriched MSigDB pathways. Malignant clusters present across more than 50% of the tumours are shown in bold. **D)** Intra- and inter-tumour GB cell state heterogeneity across our dataset. The barplot (left) shows cell state distribution summarised across each tumour. The dotplot (right) depicts the top cell state for a given sampling site within a given tumour (colour) and its relative abundance (size). **E)** Visium ST sections from each sampling site of tumour AT10 coloured according to GB cell state abundances derived from cell2location mapping. The top seven most frequently observed states at the tumour-level were visualised, with colour intensity proportional to inferred cell state abundance. Broader trends, particularly areas of cell state co-localisation are highlighted in bold text.

First, to capture intra-tumour heterogeneity, we profiled each tumour across multiple sites. Our sampling ranged from 4 sites (∼1.25 cm^3^ each) for the majority of cases to 15 sites that effectively covered one whole tumour (**Extended Data Fig. 1A, 1B**), profiling 57 tumour sites in total.

Second, to account for inter- and intra-tumour heterogeneity during multi-modal data integration, we generated paired single nuclei and spatial omic data from each tumour and sampling site. We profiled consecutive tissue sections from each site using single nuclei joint transcriptome- and chromatin accessibility-sequencing (“multiome” snRNA+ATAC-seq) and Visium spatial transcriptomics (ST) (10X Genomics) (**Fig. 1A**). Additionally, we assayed select sites with Xenium ST and immunohistochemistry as orthogonal spatial approaches, as well as whole genome sequencing of tumour microbiopsies to interrogate GB clonal architecture (**Extended Data Fig. 1B**).

Third, to enable comprehensive analysis at the level of each tumour and tumour subclone, we profiled large numbers of nuclei and multiple spatial tissue sections per site. We generated over 1 million single nuclei multiome profiles and over 0.3 million spatially resolved transcriptomes across 97 Visium sections (**Extended Data Fig. 1, Table S2**). The resulting “GBM-space” dataset, accessible on a user-friendly interactive webportal^19^ (www.gbmspace.org/), enables comprehensive multi-modal characterisation of GB architecture and presents a unique holistic tumour atlas for cancer research.

### Malignant cells vary from developmental-like to gliosis and hypoxia states

To investigate GB heterogeneity in our atlas, we first annotated malignant and TME cell states in our single nuclei multiome data. We performed standard data processing workflows and filtered nuclei based on quality of both RNA and ATAC profiles, resulting in 1,025,329 nuclei with over 85,000 nuclei per tumour on average (**Methods**, **Fig. 1A and Extended Data Fig. 2**). We then focused on the snRNA-seq data to annotate transcriptomic cell states, and examined the ATAC data to map GB gene regulatory networks in a companion study^20^.

We integrated the snRNA-seq data using scVI^21^ to account for batch variations such as 10X reaction, tumour and tumour site (**Methods**) (**Extended Data Fig. 2A**). We then distinguished malignant clusters based on inferred copy number (CN) gain of chromosome 7 and loss of chromosome 10, as well as enrichment of known GB expression programs compared to TME cells^2^ (**Fig. 1B, Extended Data Fig. 3B,C**). Finally, subclustering yielded 28 malignant clusters, including 10 highly common clusters present across 50% of the tumours, as well as 71 TME clusters (**Fig. 1C, Extended Data Fig. 4A,B**).

We annotated malignant clusters by 1) quantifying their expression of GB cell state gene programmes described by Neftel et al^2^ (**Extended Data Fig. 5A**), 2) relating them to the developing human brain cell types via label transfer from a scRNA-seq atlas^22^ (**Extended Data Fig. 5B**) and 3) gene set (MSigDB) enrichment analysis (**Extended Data Fig. 5C, Table S5**). As a result, we hierarchically grouped malignant clusters into 4 major classes and 9 subclasses of GB cell states that were each observed across multiple tumours (**Fig. 1C,D, and Extended Data Fig. 4A, Table S4**).

Two major classes of GB cell states resembled distinct developmental progenitors. The first developmental-like (dev-like) state, oligodendrocyte precursor- and neural progenitor-like (OPC-NPC-like) cells, showed universally high expression of the “OPC-like” Neftel et al programme and varying degrees of the “NPC1-like” programme (**Fig. 1C**). They were most similar to developing human brain OPCs, expressing OPC markers (*PDGFRA*, *OLIG1, SOX6)* and gene sets (**Fig. 1C and Extended Data Fig. 5B**). OPC-like clusters resembled early progenitors (i.e. OPC-like 2) differentiating towards myelinating oligodendrocytes (i.e. OPC-like 4), whereas OPC-NPC- and OPC-neuronal-like clusters additionally expressed neuroblast and neuron-related genes (*GRIK2, DLL3, GRIA4*) (**Fig. 1C and Extended Data Fig. 5B**).

The second dev-like state, neuronal progenitor- and neuronal-like (NPC-neuronal-like) cells, highly expressed the “NPC2-like” Neftel et al programme and were most similar to developmental neuroblasts and neurons (**Fig. 1C, and Extended Data Fig. 5A,B**). They expressed neuronal development genes (*MYT1L, STMN2*) and pathways, resembling early neuroblast (e.g. *SOX11^+^ DCX^+^*NPC-neuronal-like 2) differentiation towards neurons (i.e. NPC-neuronal-like 1 with elevated synaptic gene expression) (**Fig. 1C and Extended Data Fig. 5B**). These were the least abundant states, consistent with the low frequency of neuronal-like “Neural” GB tumour subtypes^1^.

The third major GB cell state exhibited a transcriptomic spectrum from astrocyte progenitor (AC-prog)-like to gliosis-like (i.e glial injury response) and hypoxic states. These states broadly expressed core astrocyte markers (*SLC1A3, GFAP)* (**Fig. 1C**). The AC-prog-like cells were enriched for the Neftel “AC-like” programme and were most similar to developmental glioblasts (**Fig. 1C and Extended Data Fig. 5A**). In AC-gliosis- and gliosis-like cells, this developmental signature was diminished and replaced by the expression of genes related to wound healing and injury response (*VIM*, *SERPINE1)* and angiogenesis (*VEGFA)* (**Fig. 1C)**. The remaining cells were characterised by hypoxia (*HILPDA)* (**Fig. 1C)**.

The gliosis-like and hypoxic cells were enriched for the Neftel “MES1” and “MES2” programmes, respectively, that were initially described as mesenchymal-like states^1,2^. Recent studies have interpreted these GB states as similar to reactive astrocytes responding to injury^4^ and glial wound healing signatures^4,8^. Here, we adopted the gliosis and hypoxia nomenclature that better reflected malignant cellular trajectories in our dataset (see sections below).

The fourth and final major malignant state represented proliferative cells. While dominated by a proliferation gene expression signature (**Fig. 1C)**, these cells also expressed dev-like or gliosis-hypoxia programmes (**Fig. 1C)**, as previously observed^2^, suggesting that both dev-like and gliosis-hypoxia states proliferate in GBs.

Our snRNA-seq atlas also captured all major cell lineages in the GB TME (**Extended Data Fig. 4B and Extended Data Fig. 6)**. OPCs, mature oligodendrocytes and myeloid cells were the most abundant, with the latter including diverse subtypes (described in detail in the final section). We finely annotated neurons, including 9 cortical layer-specific excitatory and 8 inhibitory subtypes. Astrocytes, while rare, included several homeostatic and reactive populations. Lymphocytes were similarly rare, yet spanned dendritic cells, NK cells, B cells, plasma cells and multiple T cell subtypes including CD8+, CD4+, proliferative and regulatory T cells. Finally, we also captured vasculature spanning pericytes, vascular leptomeningeal cells (VLMCs), and multiple endothelial subtypes, the latter including chr7&10 CN+ cells that may represent tumour derived vasculature^23^.

Taken together, our single nuclei transcriptomic atlas presents a deep, fine-grained cellular census of malignant and TME states in GB.

### Individual GB tumours span dev-like to gliosis and hypoxia states

Next, we examined the inter- and intra-tumour heterogeneity of malignant cell states in snRNA-seq. Most tumours showed a dominant GB cell state across the major classes of OPC-NPC-like (e.g. AT5 and 9), NPC-neuronal-like (e.g. AT15) or AC-gliosis-hypoxia (e.g. AT3 and 6) (**Fig. 1D, left**). Yet, within these groupings, tumours showed highly variable subclass composition (e.g. AC-prog-like states enriched in AT6 versus the full spectrum of AC-gliosis-hypoxia in AT3) (**Fig. 1D**).

Importantly, despite their stratification by GB cell states, each tumour contained substantial fractions of both dev-like and AC-gliosis-hypoxia cells. Two tumours (AT4 and 10) showed near equal proportions of both classes, while in others, the minority states represented 5 to 21% of all malignant cells (**Fig. 1D, Table S4**). While other studies have also noted the stratification of GBs by their cell state composition^2,4,24^, our deep profiling is the first to identify the presence of both dev-like and AC-gliosis-hypoxia malignant states within every tumour.

At the intra-tumour level, individual sampling sites also contained a mixture of GB cell states that broadly reflected the overall cellular composition of the parent tumour (**Extended Data Fig. 7B,C**). Yet, we could often identify at least one site per tumour with a unique cell state composition. In a striking example, tumour AT10 displayed distinct dominant GB cell states at each site (**Fig. 1D, right**). Consistent with other studies^5,7,18^, these observations highlight regional GB heterogeneity and show that single tumour biopsies can be significantly confounding to stratify GB tumour phenotypes.

### Spatial segregation of dev-like and gliosis-hypoxia states

Next, we examined the spatial organisation of GB cell states. While previous studies have applied ST to GB tumours, they could not distinguish fine cellular states as they relied on either the coarse-resolution Visium assay (i.e. 55 µm diameter Visium spots sampling multiple cells per location)^9^ or reference-free deconvolution of Visium into gene expression programs that, for example, could not delineate between neuronal-like cancer cells and non- malignant neurons^18^.

Here, we leveraged computational integration of our multi-modal atlas and deconvolved fine GB cell states defined by snRNA-seq in Visium ST data (n=97 sections, 338,481 spot transcriptomes) (**Fig. 1E**). While integrating paired snRNA-seq and ST from each tumour accounted for GB heterogeneity, it also enabled us to link malignant transcriptional trajectories, subclonal organisation and TME signalling that we subsequently identified in snRNA-seq to tumour spatial organisation.

We mapped malignant and TME clusters from each patient’s snRNA-seq profile to their matched Visium data using cell2location^25^ (**Fig. 1E**). This revealed striking spatial segregation of GB cell states across all tumours. In particular, malignant cells associated with gliosis and hypoxia segregated from tumour areas enriched for the dev-like states including OPC-, NPC- neuronal- and AC-prog-like cells (**Fig. 1E, Extended Data Fig. 8A**). Broadly, dev-like GB states tended to spatially intermix, whereas tumour regions dominated by gliosis-hypoxia states had more homogeneous cell composition (**Extended Data Fig. 9**), as previously shown^18^. Our spatial mapping suggested the cellular composition of GB tumours are more similar to each other than measured with snRNA-seq, with some tumours showing elevated hypoxic cell states in ST that may be lost during tissue dissociation prior to snRNA-seq (**Extended Data Fig. 7D**). Finally, our ST analysis also captured regional heterogeneity of GB cell states across different tumour sampling sites. For example, tumour AT10 site 1 was enriched for OPC-NPC-like states whereas site 3 was dominated by AC-gliosis-hypoxia (**Fig. 1E and Extended Data Fig. 7C**).

### Stereotyped spatial transitions of malignant cell states

To assess the conservation of GB spatial architecture, we examined whether malignant and TME cell states form tissue niches in a stereotyped manner across tumours (**Fig. 2A**). To do this, we first identified spatially co-localised cell states (i.e. niches) per tumour via non-negative matrix factorisation of cell2location results^25^ (**Methods**). This resulted in 192 factors summarising individual tissue niches across our cohort (**Table S7**). We then clustered these factors by their cell state similarity and identified 14 major tissue niches that recurrently appear across tumours (**Fig. 2B**).

**Figure 2:**
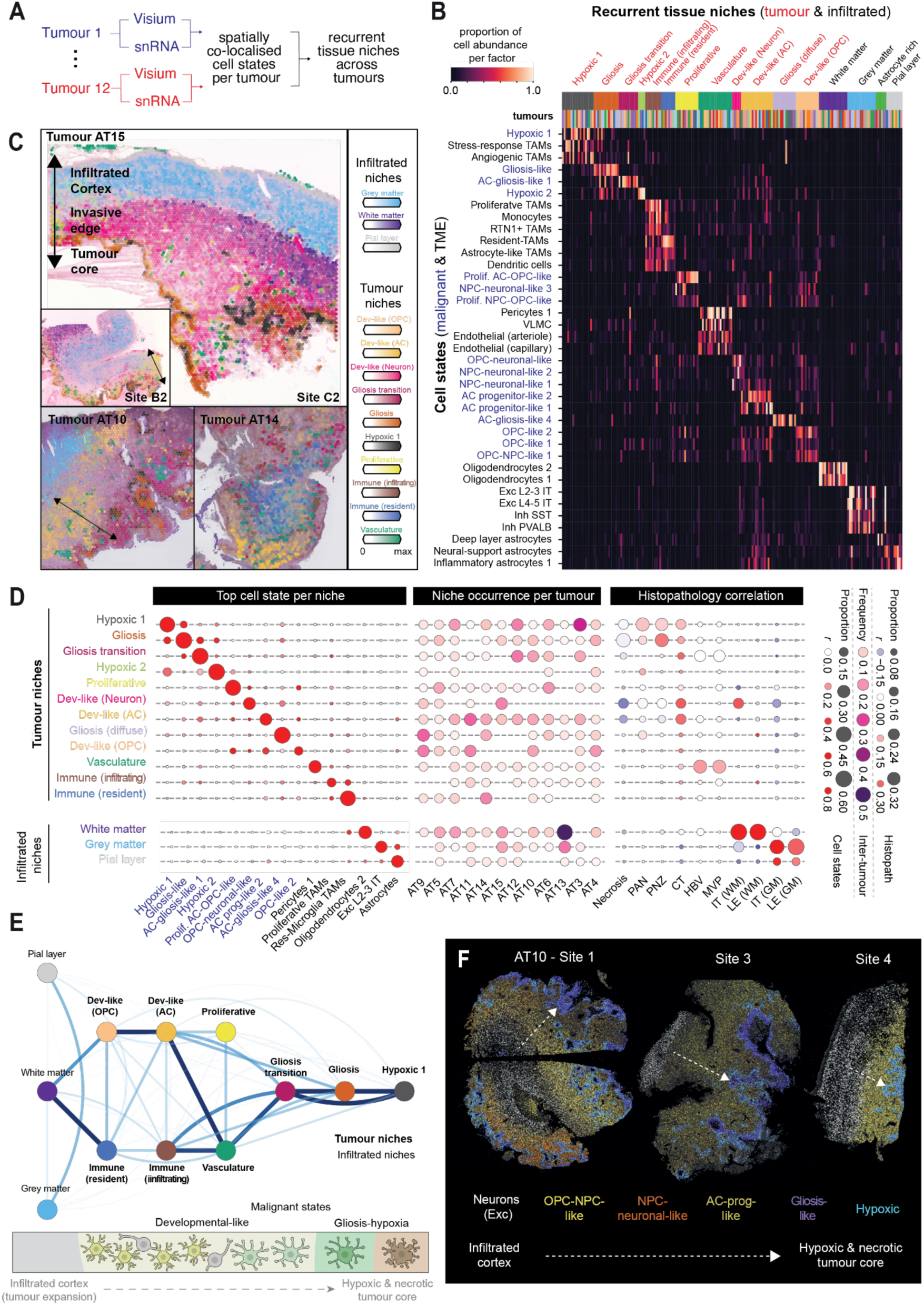
Recurrent tissue niches across GBs indicate stereotyped spatial transitions of malignant cell states. **A)** Workflow for identification of GB spatial tissue niches that recur across tumours. **B)** Distinct cellular compositions of recurrent GB tissue niches. The heatmap depicts the proportion of cell state abundance associated with a given tumour-specific NMF factor. Factors (columns) are ordered by their corresponding spatial niche clusters (top colour bar). The most frequently observed cell states for each niche are shown per row, where black and blue font colours represent TME and malignant cells, respectively. **C)** Spatial niche abundances for several Visium ST sections showing key niche distribution patterns in an intra-tumoural context (AT15) and across different tumours. **D)** Spatial niche summaries. Left: Most frequently observed cell state (column) found within each niche (row). Dotplot shows the proportion of the top cell state per spatial niche averaged across tumours (size) and the average Pearson’s correlation coefficient (r) across tumours. Centre: Relative frequency of each niche across tumours (size and colour). Right: Correlation of spatial niches with histopathological features annotated on Visium H&E images based on IvyGAP annotations (columns). Dotplot shows the proportion of spots associated with a spatial niche which overlap with a given histological feature (size) and the point biserial correlation coefficient (r) between feature annotations and spatial niche abundance. **E)** Network diagram illustrating spatial proximity between recurrent spatial niches. Edge width and colour represent the mean pairwise Euclidean distance between niche spots. **F)** Xenium sections depicting spatial maps of major cell states in tumour AT10.

We annotated the recurrent GB niches based on their cellular composition and anatomical location (**Fig. 2B, 2C**). “Tumour” niches represented malignant, immune and vasculature cell states located in the tumour core, in contrast to “infiltrated” niches of non-malignant neurons and glia in the surrounding infiltrated regions (**Fig. 2B,C**). Most tumour niches were defined by their distinct malignant cell state composition (**Fig. 2D**). *Dev-like* niches were enriched for either OPC-, NPC-neuronal- or AC-prog-like GB states, often containing multiple spatially co-localised subclusters (e.g. OPC-like 1 and 2) (**Fig. 2B, 2D**). In contrast, *Gliosis transition*, *Gliosis* and *Hypoxia* niches were dominated by a single respective malignant cluster (**Fig. 2B, 2D, and Extended Data Fig. 10A**).

The majority of tumour niches were observed across multiple donors (**Fig. 2D and Extended Data Fig. 10C**). Core malignant niches such as *Dev-like (OPC)*, *Dev-like (AC)*, *Proliferative*, *Gliosis transition*, *Gliosis*, and *Hypoxic 1* were detected in over 50% of tumours (**Fig. 2D**), while others such as *Dev-like (Neuron)* and *Hypoxic 2* were rare. Crucially, 8 out of 12 tumours (e.g. AT3, AT10) showed nearly the full complement of core malignant niches (**Fig. 2D**), suggesting these features are fundamental to GB organisation.

Previous ST studies have noted stereotyped anatomical organisation of GB cell states, locating hypoxic mesenchymal-like states at the necrotic core and dev-like states at the infiltrating edge of tumours^9,18^. To correlate our tissue niches with GB anatomy, we annotated histologically distinct anatomical features on the Visium H&E images according to the Ivy Glioblastoma Atlas (IvyGAP) criteria^26^ (n=39 Visium sections) (**Methods, Extended Data Fig. 10B**). *Dev-like* niches occurred in the cellular tumour (CT), the major tumour compartment bordering the infiltrating tumour (IT) and leading edge (LE) (**Fig. 2C, 2D**). In contrast, niches associated with gliosis and hypoxia were localised further from the infiltrating regions and formed distinct zones around necrotic areas. *Gliosis transition* (i.e. enriched for AC-gliosis states) and *gliosis* niches were found in the perinecrotic zone (PNZ) at the CT border (**Fig. 2D, mid panel**), forming spatially distinct layers in some tumours (**Fig 2C).** *Hypoxia* niches were located even deeper in tumours, extending into areas marked by pseudopalisading cells and necrosis (PAN) and necrosis (**Fig. 2C,D, and Extended Data Fig. 10B)**.

Our observations thus show that GB is organised into layered tissue niches, consistent with recent findings^18^. By incorporating snRNA-seq definitions of malignant cell identities into spatial mapping, our approach further suggested that malignant cell states show stepwise spatial transitions across niches, most evident in tissue areas with gliosis and hypoxia. To quantify their precise spatial arrangement, we calculated the pairwise spatial proximities of GB tissue niches across all tumours (n=97 Visium sections) and summarised them in a network graph (**Methods**).

We observed that GB tissue niches are ordered along a major anatomical axis between the infiltrating tumour edge and the necrotic tumour core (**Fig. 2E**). Amongst core malignant niches, *Dev-like (OPC)* was found closest to infiltrated white matter. *Dev-like (AC)* spatially overlapped with the *Dev-like (OPC)*, yet extended deeper into the tumour towards successive layers of *Gliosis transition*, *Gliosis* and *Hypoxia* (**Fig. 2E**). *Dev-like (Neuron)* was similarly positioned near the infiltrated regions when present (**Fig. 2C**). In the tumour that we grid-profiled in whole (AT15), these gradual transitions from dev-like niches to gliosis and hypoxia were observed across each sampling site (**Extended Data Fig. 11A,B**).

We validated these observations with the orthogonal imaging-based Xenium ST technology. We applied a 315-plex probe panel, including 62 markers of GB and TME cell states curated from our snRNA-seq data (**Methods, Table S8**), to 17 sections from 8 tumours. We annotated Xenium data at single cell level by integrating them with patient-matched snRNA-seq data via Tangram^27^. This validated the zonation of malignant cell states, where dev-like states bordered infiltrated regions marked by neurons, contrasting with the gliosis and hypoxia states situated further around necrotic and perinecrotic regions (**Fig. 2F and Extended Data Fig. 11C**). Importantly, profiling centimeter-scale tumour sections, we observed this pattern at each sampling site (**Fig. 2F**), indicating stereotyped spatial organisation throughout each tumour.

The GB tissue niches also reflected TME organisation. *Vasculature* captured pericytes, VLMCs and endothelial cells (**Fig. 2B**) and anatomically correlated with microvascular proliferation (MVP) and hyperplastic blood vessels (HBV) (**Fig. 2D**). *Dev-like (AC)*, *Gliosis transition* and *Proliferative* malignant niches were found in close proximity to *Vasculature* (**Fig. 2E**). Additionally, two *Immune* niches captured distinct myeloid cell compartments (see last section). Amongst infiltrated regions, *White matter* was found in close proximity to tumour leading edge (**Fig. 2B-E**), consistent with GB invasion observed through cortical white matter^28–30^.

Taken together, we reveal stereotyped stepwise spatial transitions of malignant cells from dev-like to gliosis and hypoxia states in GB.

### A putative malignant cell trajectory towards gliosis and hypoxia

We hypothesised that the stereotyped spatial organisation of malignant cell states relates to their cellular trajectories. Specifically, we hypothesised that as GB tumours expand at their leading edge into normal brain tissue, the dev-like malignant cells progressively transition towards gliosis and hypoxia states in a recurrent manner. To test this, we first examined putative malignant trajectories related to gliosis and hypoxia.

In the snRNA-seq data, we observed a gene expression trajectory from AC-prog-like malignant states to gliosis then hypoxia (**Fig. 3A**). The AC-prog-like states showed the highest levels of developmental and homeostatic astrocyte gene expression (*EGFR, SLC1A3, AQP4, ALDH1L1*). These were downregulated in AC-gliosis-like cells and replaced by a transcriptomic program resembling astrogliosis, including genes from the integrin-N-cadherin pathway (*ITGAV, ITGB1, CDH2*) associated with glial scar formation^31^. This program expanded in gliosis-like cells to genes associated with JAK-STAT signalling (*JAK2, STAT3, SMARC4*), wound healing (*ANXA2, ANXA5, YAP1, F1, AKAP12*), cytokine production (*IFI16, IL6R, IL1R1*), coagulation (*SERPINE1*) as well as angiogenesis and hypoxia (*VEGFA*)^32–34^. This trajectory terminated in hypoxia (*HILPDA, BNIP3L, VEGFA*) and stress response gene expression (*JUN, FOS, HSPA1B*) (**Fig. 3A and Extended Data Fig. 12A**).

**Figure 3:**
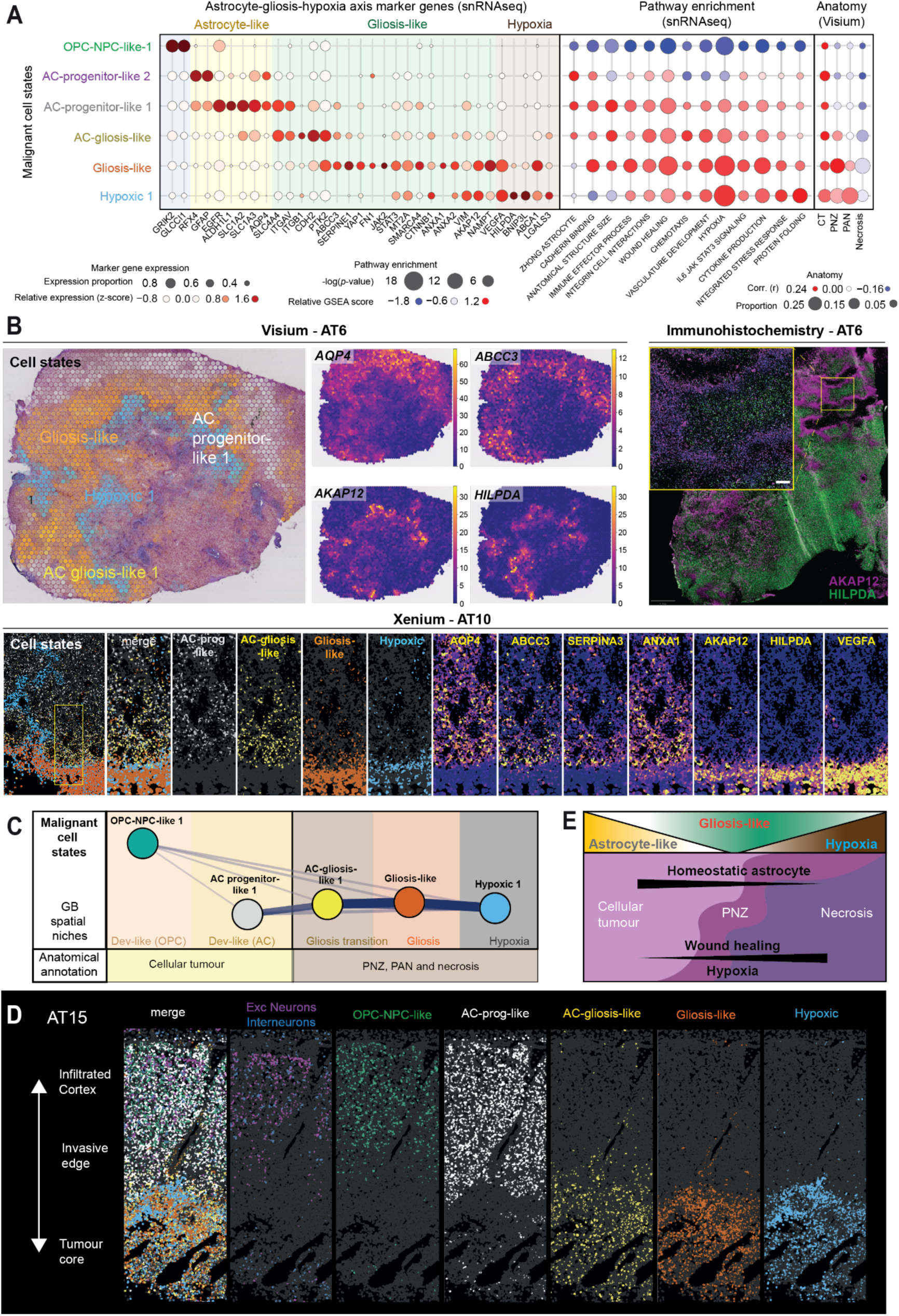
Spatial zonation of a putative malignant trajectory from astrocyte progenitor-like states to gliosis and hypoxia. **A)** Transcriptional continuum across AC-gliosis-hypoxia GB cell states. Left: Marker gene expression across states in snRNA-seq, showing relative z-score normalised gene expression (colour), and proportion of cells per state expressing the gene (size). Centre: Pathway enrichment across states in snRNA-seq, showing the z-score normalised enrichment score (colour) and the −log(p-value) of each comparison (size). Right: Spatial correlation of cell states with anatomical features in Visium ST data, showing the Pearson’s correlation coefficient (r) (colour) and relative abundance of a cell state within feature-annotated Visium spots (size). **B)** Spatial zonation of major AC-gliosis-hypoxia GB states and marker gene expression in ST data and immunohistochemistry validation. Top left: Cell state abundance (left panel, colour) and gene expression (right panel, colour) in Visium ST data. Top right: Antibody staining showing distinct zonation of AKAP12 (gliosis-hypoxia) and HILPDA (hypoxia). Bottom: Cell states (left panels, colour) and marker genes (right panels, colour) in Xenium ST data. **C)** Network diagram summarising the spatial proximity of relevant cell states in Visium data, with coloured banners indicating the primary spatial niche of each cell state, and boxes indicating the associated histological features. **D)** Xenium section demonstrating zonation of AC-gliosis-hypoxia cell states ranging from infiltrated grey matter to the necrotic tumour core. **E)** Schematic summarising the AC-gliosis transcriptional gradient in a histological context, including relevant functional axes.

While gliosis and hypoxia-associated malignant states have been described as mesenchymal-like^2^ and wound healing and injury response genes overlap with epithelial-to-mesenchymal transition (EMT) programs^35,36^, we observed negligible and non-specific expression of key EMT regulators (e.g. *SNAI1/2, TWIST1/2, ZEB1/2*) in these malignant states (**Extended Data Fig. 12A**). Hence, our nomenclature places these states in the more specific biological context of glial injury response and hypoxia, putatively stemming from developmental astrocyte-like malignant cells.

The molecular trajectory was correspondingly patterned in space, manifesting as distinct concentric segregation of AC-gliosis-hypoxia states in Visium and Xenium ST data (**Fig. 3B,C**). AC-prog-like cell states were the most spatially distinct, located in the non-necrotic CT in the *Dev-like (AC)* niche (**Fig. 3B-D**, **Fig. 2D**). AC-gliosis- and gliosis-like states formed *Gliosis transition* and *Gliosis niches*, respectively, that spatially intermixed to varying degrees. In some tumours, there was a clear gradient where AC-gliosis-like was located at the margin of CT and transitioned into gliosis-like within the PNZ (e.g. AT10), whereas in others they broadly overlapped (e.g. AT6) (**Fig. 3B**). Hypoxic cell states were located either within or adjacent to the necrotic core of patient tumours, including the PNZ and PAN (**Fig. 3B and Fig. 2D**). Consistently, the marker genes of these states showed spatially zonated expression in ST data including *AQP4* (AC-progenitor), *ABCC3* (AC-gliosis), *AKAP12* (Gliosis) and *HILPDA* (Hypoxia), which we also validated using orthogonal immunohistochemistry (**Fig. 3B and Extended Data Fig. 12B**).

Taken together, we identify putative cellular transitions of malignant cells from developmental astrocyte-like to gliosis then hypoxia dominated states, which unfolds across an anatomical axis from tumour expansion towards hypoxia and necrosis (**Fig. 3E**).

### Shared clonal origins of dev-like and gliosis-hypoxia malignant cells

To characterise GB cell state transitions, we first set out to validate whether dev-like and gliosis-hypoxic states could emerge within the same tumour subclones (i.e. genetically related malignant sublineages). Previous studies have applied copy number aberration (CNA) inference to single cell or spatial RNA-seq data to resolve GB subclones^2,3,9^. Here, we initially leveraged multiome data to validate inferred CNAs across multiple modalities and performed joint clone calling on snRNA-seq and snATAC-seq profiles of malignant cells (**Fig. 4A**). We inferred clones independently on each modality, then greedily matched them on the basis of shared cells and CNA profile similarity (**Methods**).

**Figure 4:**
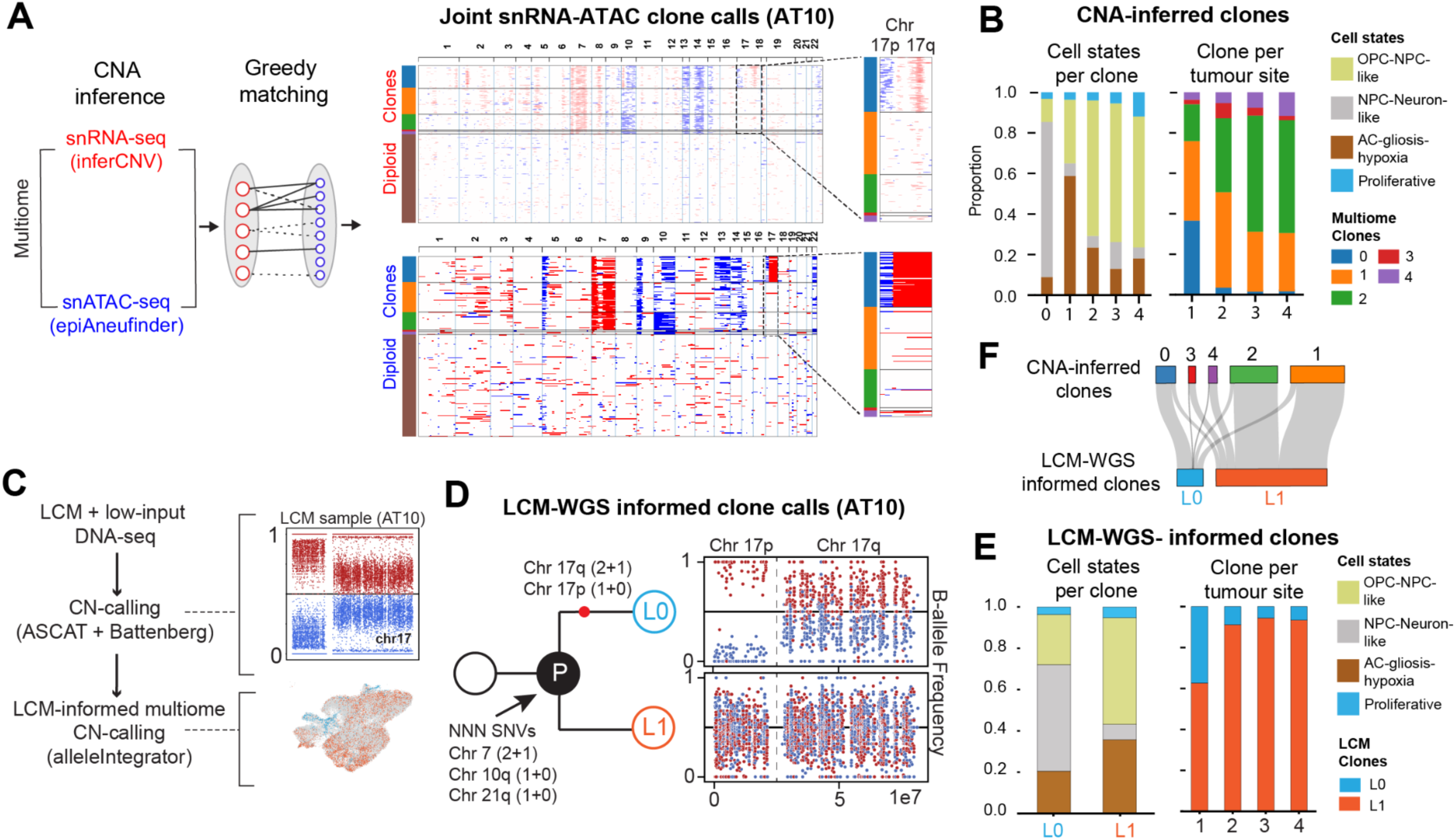
Developmental-like and gliosis-hypoxia states share subclonal origins. **A)** Multiome CNA-calling workflow. The joint clonal annotation in tumour AT10 is illustrated by heatmaps from paired snRNA- (top) and snATAC-seq (bottom) data showing putative copy gain (red) and copy loss (blue). The harmonised clone annotations are indicated by the left sidebar. Insets highlight a major lineage-defining CN changes to chr17. **B)** Barplots depicting cell state distributions across clones (right) and clone distribution across tumour sampling sites (left) in CNA-inferred clones. **C)** LCM-informed CNA calling workflow to predict snRNA-seq CN events via alleleIntegrator. The scatterplot (top right) represents the B-allele frequency (BAF) for sample PD60966a_lo0005. Colours correspond to SNP allele frequency of the major (red) and minor (blue) alleles. The UMAP (bottom right) shows alleleIntegrator clone calls superimposed on the visualisation such that cells with chr17 CNs (light blue) can be distinguished from cells without chr17 CNs (orange). **D)** LCM genotypes in the snRNA-seq data. Left: phylogeny illustrating lineage defining mutations. Right: BAF plot showing the LCM-based SNP-phasing applied to genotype the single nucleus data for two putative clones. **E)** Barplots depicting cell state distributions across clones (right) and clone distribution across tumour sampling sites (left) in LCM-WGS informed clones. **F)** Sankey plot depicting concordant clone assignment across multiome and LCM-informed CNA calling methods per cell.

This resulted in 58 tumour subclones across 12 patient tumours (5 per tumour on average) with chr7 and chr10 CNs distinguishing them from non-malignant diploid cells (**Fig. 4A and Extended Data Fig. 13A**). The majority of subclones contained both dev-like and gliosis-hypoxic cells (**Fig. 4B, Extended Data Fig. 13B,C, Table S9**), with the minority states making up an average of 15% (ranging from 3-49%) of each clone. The subclones within each tumour showed similar distributions of coarse malignant states (e.g. OPC-NPC vs NPC-neuronal) and tended to mirror the overall cellular composition of tumours, though we observed greater clonal variability in distributions of fine-grained GB states (**Fig. 4B and Extended Data Fig. 13B**).

Surprisingly, in three tumours, we identified individual subclones with highly distinct cellular compositions. In AT10, clones 0 and 1 were preferentially enriched for NPC-neuronal-like and AC-gliosis-hypoxia states, respectively (**Fig. 4B**). Similarly, AT15 clones 3 and 5 were enriched for AC-gliosis-hypoxia and OPC-like states, respectively, and AT6 clone 0 was enriched for OPC-like states (**Extended Data Fig. 13B**). Distinct CNAs marked these clones, such as chr17p loss and chr17q gain in AT10 clone 0 (**Fig. 4B**). While these clones still contained both dev-like and gliosis-hypoxia cells, these patterns imply that late genomic alterations in GB phylogenies can influence malignant cell states.To identify CNAs and subclones directly at the DNA level rather than inferring them from snRNA/ATAC-seq data, we performed whole genome sequencing (WGS) on tissue collected via laser capture microdissection (LCM)^37^. Each cut consisted of 100-200 μm^2^ sized regions of interest (ROIs) z-stacked across two adjacent H&E sections. We profiled multiple ROIs per primary tissue section across two sampling sites of two tumours (AT3, AT10), generating 61 high-quality WGS profiles (**Extended Data Fig. 14A**). In general, we found that most detected CN events are truncal, with only a minority of events appearing in specific tumour ROIs (**Extended Data Fig. 14B, Methods**), suggesting that GB subclonal diversification is a relatively late event. Notably, we detected a CN loss in chr17p and gain chr17q in one ROI from AT10 (**Fig. 4C and Extended Data Fig. 14B**), validating our multiome CN calls (**Fig. 4B**).

To understand whether the genotypes of subclones can influence their composite GB cell state phenotypes, we defined tumour subclones in the multiome data based on the CN events validated at the DNA level. We used the alleleIntegrator tool^38^ to leverage allelic imbalances identified in the LCM-WGS data to predict corresponding CNs in matched snRNA- seq profiles. In AT10, this approach validated chr7 gain and chr10 loss in malignant cells and distinguished two subclones based on chr17-alterations (**Fig. 4D**). Both subclones contained a mixture of dev-like and gliosis-hypoxia cells (**Fig. 4E**). Yet, strikingly, clone L0 was enriched for NPC-neuronal-like cells and marked by chr17p loss and chr17q gain (**Fig. 4D,E**). This mirrored the cellular composition and CN profile of clone 0 identified by our orthogonal multiome CNA inference (**Fig. 4A,B**). Correspondingly, the cell assignments to chr17-altered clones were highly consistent across our two CN calling approaches (**Fig. 4F**), validating our joint clone calls from multiome data by the WGS-derived “ground truth”.

Taken together, our paired genomic, epigenomic and transcriptomic data demonstrate shared clonal origins of dev-like and gliosis-hypoxia cell states, consistent with our trajectory hypothesis. Surprisingly, we also identified that late genomic alterations can alter malignant cell states and expand intratumour heterogeneity, the first such demonstration at cellular resolution in GB.

### Fine-grained spatial intermixing of GB subclones

Given the spatial segregation of GB cell states, we inquired whether tumour subclones are also spatially organised. Based on the shared clonal origins of dev-like and gliosis-hypoxia cell states (**Fig. 4**), we expected individual subclones to contribute to multiple spatial tissue niches (**Fig. 2**). While GB clonal distribution has been studied at coarse resolution using bulk tumour biopsies^5,7^, the fine-grained spatial organisation of clones and how they relate to tissue niches is poorly understood.

Initially, we investigated clonal organisation at centimeter-scale resolution from tumour sampling sites. Based on multiome CNA inference, we detected multiple subclones within each site (**Fig. 4B and Extended Data Fig. 15A**). The clone distribution was largely consistent across different tumour sites, with notable exceptions including AT10, where the chr17-altered clone 0 is predominantly located in site 1 of 4 (**Fig. 4B and Extended Data Fig. 15A**). Consistently, our orthogonal LCM-WGS approach found the corresponding clone L0 enriched in the same site (**Fig. 4E**). These observations suggest broad spatial intermixing of GB clones, as previously shown^5,7^, and that late genomic alterations can give rise to spatially restricted clones.

To investigate clonal organisation at higher spatial resolution and identify clones at the DNA level, we examined clonal distributions in our LCM-WGS data profiling ∼100x100 µm tissue areas. Considering the clonal intermixing described above and that our LCM cuts span multiple cells, we used PyClone-VI^39^ to infer clonal population structure of LCM ROIs based on single nucleotide variant (SNV) allele frequencies (**Fig. 5A, Methods**). We focused on tumour AT3 that yielded 35 high-quality genomes from LCM ROIs across two sampling sites, and identified 17 SNV clusters (i.e. putative subclones) in total (**Extended Data Fig. 14A, right**). Individual SNV clusters were present across multiple LCM ROIs and individual ROIs contained multiple SNV clusters, indicating clonal intermixing at fine spatial resolution (**Fig. 5B**).

**Figure 5:**
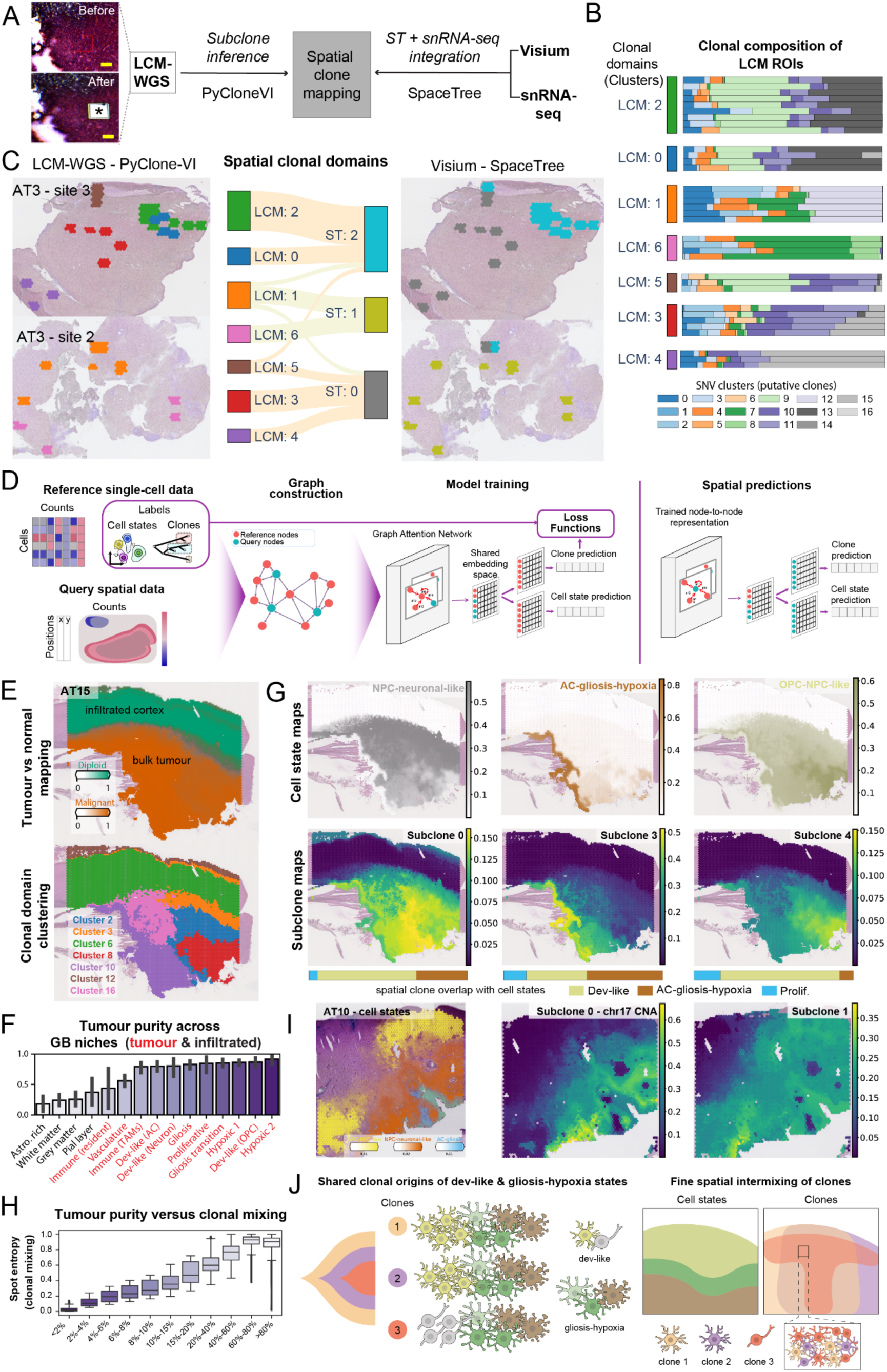
GB spatial tissue niches are polyclonal. **A)** LCM-WGS- and SpaceTree-based spatial clone mapping workflows. The H&E images show a tumour ROI before and after LCM cutting, marked by an asterisk. Scale bars: 100 µm. **B)** Spatially intermixed GB subclones in LCM-WGS data. Barplot summarises pyclone-VI clone distributions for each LCM ROI and clonal cluster. **C)** Spatial clonal domains compared across LCM-WGS pyclone deconvolution and Visium ST SpaceTree mapping across matching tissue locations. Domains were defined by Bray-Curtis clustering of clone distributions within each ROI across each modality H&E images depict tissue locations used for LCM colored by SpaceTree clone clusters (left) and pyclone-VI clone clusters (right). The sankey plot demonstrates ROI cluster assignment between modalities. **D)** SpaceTree workflow for joint spatial mapping of cell states and tumour subclones via integration of ST and snRNA-seq data. **E)** SpaceTree distinguishes tumour versus normal regions and resolves spatial clonal domains in tumours. Top: Visium section from tumour AT15 showing SpaceTree mapping of diploid versus malignant subclones to infiltrated and bulk tumour regions, respectively. Bottom: Spatial clonal domains identified by clustering SpaceTree-derived clonal composition of Visium spots. **F)** Tumour purity of infiltrated versus tumour tissue niches estimated by SpaceTree, summarised at the Visium spot level (n=97 Visium sections). **G)** GB subclones spatially overlap with both dev-like and gliosis-hypoxia cell states. SpaceTree derived cell state (top panel) and clonal (bottom panel) maps in tumour AT15 are shown. The barplots summarise the distribution of cell states across spots in which a given clone is the most abundant. **H)** Clonal entropy of Visium spots based on SpaceTree mapping binned across tumour purity fractions. High entropy (i.e. clonal intermixing) is observed in tissue regions with high tumour purity. **I)** SpaceTree cell state (left) and clonal (centre and right) maps of tumour AT10 resolves distinct regional distributions of chr17-CNA subclone. **J)** GB clonal organisation summary diagram.

Clustering ROIs based on their subclonal composition (**Fig. 5B**), we identified broad spatial clonal domains (**Fig. 5C**). Neighboring tissue regions had similar polyclonal composition, giving rise to several millimeter-wide clonal domains (e.g. LCM ROI domains 0 & 2 on site 3). Broad spatial gradients of subclones demarcated neighbouring domains, such as SNV cluster 9 spanning domains 0, 2 and 5 and SNV cluster 15 appearing in domains 3 and 4 (**Fig. 5C**). Different tumour sampling sites showed largely distinct polyclonal composition (**Fig. 5C**), displaying more extensive differences than we observed with multiome CNA inference (**Extended Data Fig. 15C,D**), likely owing to the better sensitivity of SNV-based clone calls from WGS data. We inferred phylogenetic trees of AT3 subclones and observed early SNV clusters to be widely spatially spread while late clusters showed variable spatial distribution, including those restricted to few sites (**Extended Data Fig. 15C**).

### Joint spatial modeling reveals GB tissue niches are polyclonal

Next, given their broad spatial distribution, we sought to validate that GB subclones contribute to multiple tissue niches spanning the putative trajectory from dev-like to gliosis- hypoxia states. Consistently, in the LCM-WGS data, we observed that individual clones (e.g. SNV cluster 9) and larger clonal domains (e.g. LCM cluster 0, 2 & 5) spanned tissue areas with diverse malignant cell states and tissue niches based on the adjacent Visium sections (**Extended Data Fig. 15D**). However, our LCM-WGS findings were limited to select ROIs from two tumours due to the high cost and low throughput of this methodology. Hence, we aimed to resolve spatial clonal patterns in our Visium ST dataset that comprehensively profiles 12 tumours (n=97 sections) at high resolution. The ST profiles in Visium data also enabled us to directly relate clonal domains to transcriptomic cell states and tissue niches (**Fig. 2**) on the same tissue section.

Previous studies have used CNA inference on Visium data to chart subclones in GB^9^ and other tumours^40^. However, these approaches assumed monoclonal composition of Visium spots, which could not resolve the finely intermixed GB subclones. To address this challenge, we developed SpaceTree, a computational framework to jointly deconvolve tumour subclones and cell states in ST data by leveraging a reference scRNA-seq dataset (**Fig. 5D**). SpaceTree utilises a multi-task graph neural network architecture coupled with label propagation to integrate the reference scRNA-seq and query ST datasets, enabling joint spatial mapping of clones and cell states annotated in the single cell data (**Fig. 5D**, described in detail in **Supp. Comp. Note**).

We applied SpaceTree to integrate our GB snRNA-seq and Visium ST data per tumour site, leveraging our granular cell annotation (**Fig. 1B,C**) and joint multiome clone calls (**Fig. 4B**). We optimised SpaceTree hyperparameters to match cell2location cell type mapping (**Supp. Comp. Note**). Initially, we validated that Spacetree can distinguish between infiltrated (i.e. largely normal cells) versus tumour regions. Consistently, SpaceTree mapped diploid clones containing TME cells to sparsely infiltrated regions, while malignant subclones were enriched in the tumour core (**Fig. 5E, top panel**). We then quantified the fraction of malignant cells per Visium spot (i.e. tumour purity) from SpaceTree estimates and summarised it by GB tissue niches across all tumours (**Fig. 2C**). Consistently, this showed predominant enrichment of malignant cells to malignant GB niches (e.g. *Dev-like, Hypoxic*) over infiltrated (e.g. *Grey matter*) or TME (e.g. *Vasculature*) niches (**Fig. 5F**).

SpaceTree revealed widespread spatial intermixing of subclones across all tumours (**Fig. 5G and Extended Data Fig. 16A**). To quantify the extent of clonal mixing, we calculated the clonal entropy per Visium spot across all tumours. Spots with low fraction of malignant cells (i.e. infiltrated and TME tissue niches with low tumour purity) showed low clonal entropy, suggesting they are dominated by diploid clones (**Fig. 5H**). In contrast, spots with high fraction of malignant cells (i.e. malignant tissue niches with high tumour purity) showed high clonal entropy, indicating fine-grained spatial mixing of GB subclones and the polyclonal nature of GB tissue niches (**Fig. 5H**).

SpaceTree also resolved broad spatial domains of subclones across GBs (**Fig. 5E and Extended Data Fig. 16B**). Individual clonal domains and their constituent subclones overlapped with diverse malignant cell states (**Extended Data Fig. 16D**). For example, AT15 subclones 0, 3 and 4 each overlapped with both dev-like and gliosis-hypoxia cell states (**Fig. 5G**). Yet, we also observed regional clonal enrichment, such as clone 3 contributing more heavily to gliosis-hypoxia (**Fig. 5G**). Notably, AT10 clone 0, distinguished by chr17 alterations (**Fig. 4**), was spatially enriched in tissue regions with NPC-neuronal-like cells (**Fig. 5I**).

Finally, we also validated SpaceTree by orthogonal LCM-WGS mapping of adjacent tissue sections (**Methods**). Across matching tissue regions, we observed concordance between clonal domains mapped by SpaceTree deconvolution of Visium ST data and those identified by SNV-based deconvolution of LCM-WGS data (**Fig. 5B, Methods**). While SNV-based mapping provided higher granularity, the spatial clonal domains broadly matched across the two modalities (**Fig. 5B**).

In summary, we present a new framework for the cellular and spatial organisation of GB subclones (**Fig. 5I**). We demonstrate that individual GB subclones give rise to both dev- like and gliosis-hypoxia states, and that they finely spatially intermix in tumours, organising into broad clonal domains cutting across GB tissue niches. We also show that late genomic alterations (i.e. chr17 CNA in AT10) can alter the cellular composition and localisation of GB subclones to expand intra-tumour heterogeneity.

### A recurrent trajectory from dev-like to gliosis and hypoxia states dominates GBs

Having validated the shared clonal origins of dev-like and gliosis-hypoxia states in single cell (**Fig. 4**) and spatial data (**Fig. 5**), we investigated whether they form a cellular trajectory tractable across GBs. Specifically, we examined whether the transition from dev-like to gliosis-hypoxia states represents the major transcriptomic trajectory across distinct GB subclones and tumours, and whether this involves conserved gene expression programs.

Given their spatial zonation observed throughout each tumour (**Fig. 2E,F**), we hypothesised dev-like malignant cells continuously transition into gliosis and hypoxia states during tumour expansion. The rapid expansion rate of GBs, which double in size every 50 days^41^ supported the likelihood that our “snapshot” snRNA-seq profiling can capture the different stages of these putative transitions. To infer cellular trajectories in an unbiased manner from snRNA-seq data, we used cell2fate, a probabilistic RNA velocity model^42^ (**Fig. 6A**).

**Figure 6:**
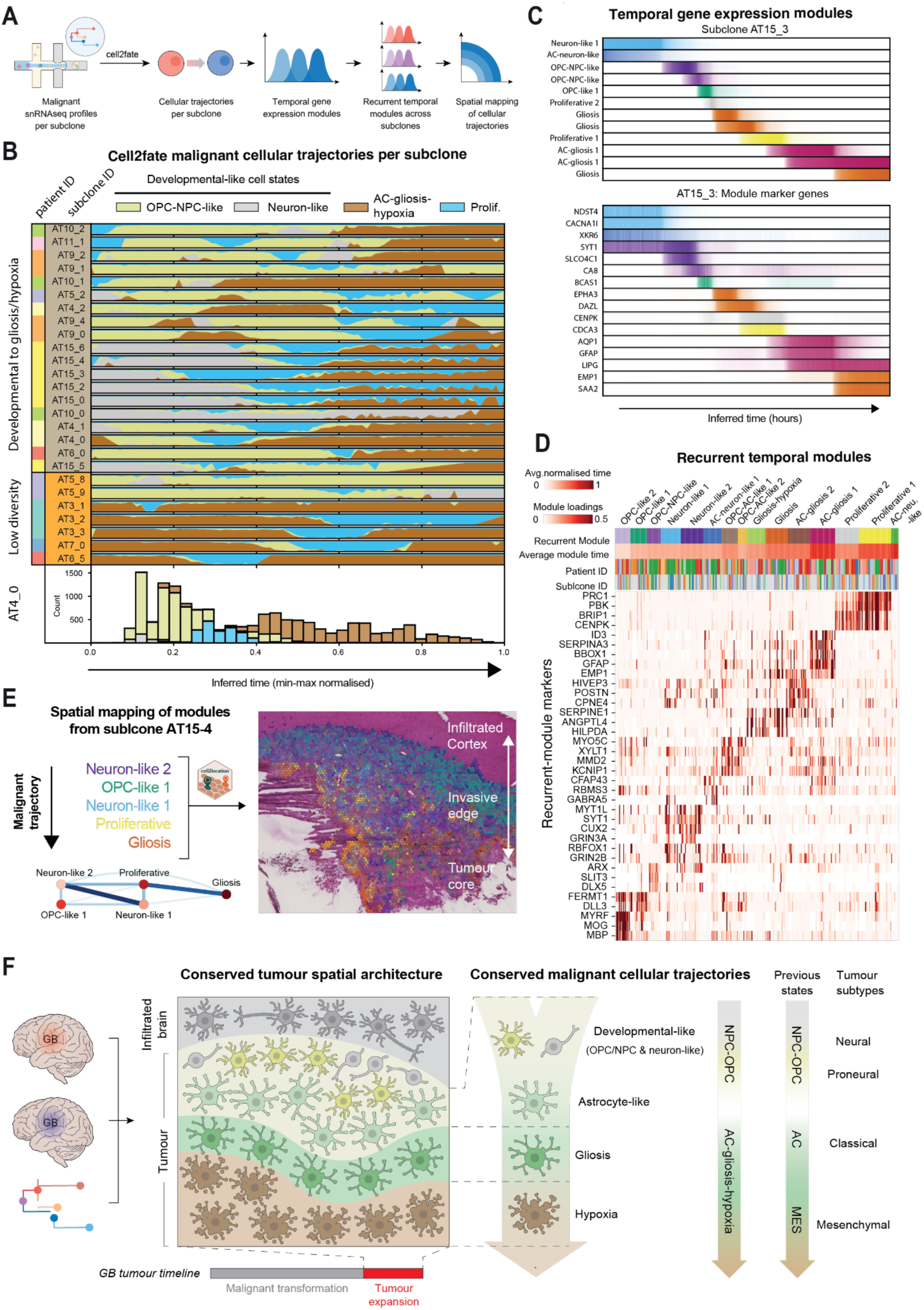
A recurrent malignant cell state trajectory across GB tumours. **A)** Cellular trajectory analysis workflow. **B)** Cell2fate analysis resolving a cellular trajectory from dev-like to AC-gliosis-hypoxia states across GB subclones and tumours. Top: Streamgraph plots show cell state occurrence across cell2fate-inferred trajectory time across 26 tumour subclones (rows). Clones are clustered based on the ranked temporal order of coarse-grained cell states, distinguishing those spanning diverse GB cell states (top panels) versus low diversity clones dominated by one of the aforementioned cell states (bottom panels). Bottom: Barplot showing the abundance of cell states in clone AT4_0 across inferred time. **C)** Activation patterns of genes (top) and meta-modules (bottom) for clone AT15_3 summarising the inferred cellular trajectory. Cell2fate derived activation plots of temporal gene expression modules (top) and top marker genes of each module (bottom) with matched colours. **D)** Recurrent temporal gene expression modules are conserved across the trajectories of 26 tumour subclones. Heatmap displays the module loadings for top module markers across each individual subclone. Columns are clustered by Jaccard similarity index of top genes (n=200). Colour bars correspond to recurrent temporal module annotations, average latent time across each recurrent module, tumour and clone labels. **E)** Spatial mapping of key temporal modules (colours) from subclone AT15_4 onto its matching Visium section (right). Network diagram (bottom) shows the mean pairwise euclidean distance between temporal modules for the associated Visium section. **F)** Summary diagram of the spatiotemporal malignant cell trajectory identified across GBs. The diagram relates different stages of this trajectory to GB cell states and tumour subtypes described by previous studies.

We applied cell2fate to the snRNA-seq profiles of individual tumour subclones, avoiding confounding genetic differences across clones and tumours. Out of 58 subclones, 26 subclones from 9 tumours showed robust cellular trajectories based on their inferred duration^42^, where GB cell states segregated across distinct latent time windows (**Fig. 6B and Extended Data Fig. 17A, Methods).** The remaining subclones showed poorly discernible trajectories based on their inferred duration, likely due to either small numbers of cells or low state diversity captured per clone (**Extended Data Fig. 17B**). Across the reconstructed trajectories, the majority of subclones (19/26, >73%) displayed temporal transitions from dev- like to AC-gliosis-hypoxia (**Fig. 6B**). We observed this trajectory across 7 out of the 9 tumours (e.g. AT4, AT10, AT15) and quantified temporal ordering of GB states from dev-like to AC- gliosis-hypoxia (**Fig. 6B, bottom panel, Extended Data Fig. 17D**). The minority of clones that did not show this pattern (7/26) were dominated by an individual state and displayed relatively lower cell state transition quality scores (**Fig. 6B, Extended Data Fig. 13B, Extended Data Fig. 17A, Methods**).

At a finer-grained cellular level, both NPC-neuronal-like (AT15_0) or OPC-NPC-like (AT6_0) states led to AC-gliosis-hypoxia (**Fig. 6B**). Proliferating cells distributed across this trajectory and were most frequently observed mid-latent time (**Extended Data Fig. 17D**). Within dev-like or AC-gliosis-hypoxia classes, fine-grained malignant states did not show highly distinct temporal ordering and consistently displayed low cell state transition scores (**Extended Data Fig. 17B**).

Importantly, we observed the transition from dev-like to AC-gliosis-hypoxia states commonly across tumours of all presumed GB subtypes, including the OPC-NPC-like enriched “proneural” (e.g. AT5), the NPC-neuronal-like enriched “neural” (i.e. AT15) and the AC-gliosis- hypoxia enriched “classical-mesenchymal” (e.g. AT6) tumours (**Fig. 6B** and **1D**). Within each tumour, multiple subclones also shared this trajectory (e.g. AT15_0 and AT15_3) (**Fig. 6B)**. This included tumour AT10, where both NPC-neuronal- and OPC-NPC-enriched clones distinguished by chr17 alterations (**Fig. 4A,B**), converged on AC-gliosis-hypoxia (**Fig. 6B)**. Taken together, these findings identify a dominant malignant cellular trajectory across GBs, manifesting across tumour subtypes as well as genetically distinct subclonal lineages.

Next, we examined whether this cellular trajectory is molecularly conserved across subclones and tumours, leveraging temporal gene expression modules (i.e. RNA velocity modules) identified by cell2fate^42^. Malignant cells in each subclone progressed through successive gene expression modules that were sequentially activated and temporally overlapped (**Fig. 6C**). Clustering modules based on their marker gene similarity (**Methods**), we identified 15 meta-modules representing recurrent temporal transcriptional programs across subclones and tumours (**Fig. 6D, Table S10**). Based on their GB cell state (**Extended Data Fig. 18A**) and MSigDB enrichment (**Table S10**), we annotated these meta-modules using a nomenclature similar to GB cell states (**Fig. 6D**). Developmental-like modules occurred early in the GB cellular trajectory across subclones and tumours, and shared marker genes with OPC-like (e.g. *MBP, MOG*) and NPC-neuronal-like (e.g. *MYT1L*, *SYT1*) cell states. In contrast, AC-gliosis-hypoxia modules occurred late and displayed corresponding marker genes (e.g. *GFAP, SERPINE1, ANGPTL4, HILPA)* (**Fig. 6D**). Proliferative modules frequently preceded or followed AC-gliosis-hypoxia. Hence, these modules demonstrate a shared malignant gene expression trajectory across GBs.

While the recurrent temporal gene expression modules broadly reflected GB cell state phenotypes and included known regulators of dev-like (e.g. *ASCL1, SOX10)*^43,44^ and gliosis- hypoxia (e.g. *FOS*^45^) states (**Extended Data Fig. 18C, Table S10**), they also pointed at novel candidate regulators and effectors of GB trajectories. For example, *MYRF*, which transcriptionally co-regulates oligodendrocyte maturation and myelination with SOX10^46,47^, was enriched in the earliest developmental modules (**Fig. 6D**). GBM cells upon white matter invasion have been shown to acquire mature oligodendrocyte-like states induced by SOX10^48^. In contrast, *HIVEP3*, which regulates transcription mediated by nuclear factor kappa-B (NF- κB)^49^ and c-Jun^50^, was enriched in late gliosis-hypoxia modules (**Fig. 6D**). Accordingly, c-Jun is implicated in regulation of mesenchymal-like states^51^ and *HIVEP3* is also implicated in Interleukin signalling in glioma^52^.

To relate temporal malignant gene expression trajectories to clinical outcomes, we performed survival analysis on bulk RNA-seq data from TCGA and CPTAC cohorts (n=235 GBs)^53–55^ using gene sets from the recurrent gene expression modules. We found that patients with high relative expression of late-occurring Gliosis-hypoxia and Proliferative module markers showed worse survival than those enriched for early-occurring OPC-like 1 module (*p*<0.007) (**Extended Data Fig. 18D**), consistent with poor prognosis of “mesenchymal-like” GBs1,56.

To spatially validate the GB cell state trajectory identified above, we mapped clone-specific temporal gene expression modules to our Visium ST data. As expected, we observed Dev-like modules adjacent to the infiltrated cortex, followed by proliferative and gliosis-like modules near to the necrotic and hypoxic core (**Fig. 6E**). Pairwise distance computation also suggested a trajectory in which dev-like modules transition into proliferative states and terminate in gliosis-like gene expression programs (**Fig. 6E**).

Taken together, we identify a spatially patterned and molecularly conserved malignant cell trajectory across GBs. Combined with our fine-grained mapping of AC-gliosis-hypoxia trajectories (**Fig. 3E**), these data indicate stepwise transitions of malignant cells from dev-like towards gliosis and hypoxia-defined states throughout tumour expansion (**Fig. 6F**).

### The malignant cell trajectory is coupled to a regionalised myeloid microenvironment

Finally, we investigated whether the GB TME is specialised to accommodate the trajectory of malignant cells from dev-like towards gliosis and hypoxia states. We focused on myeloid cells that were the most abundant TME cell type residing in tumour niches (**Table S3, Fig. 2B**). We annotated myeloid subclusters in our snRNA-seq atlas by their expression of 1) canonical marker genes, 2) MSigDB gene sets and 3) gene programs distinguishing functional myeloid subtypes across gliomas from Miller et al.^16^. We classified 11 myeloid subtypes across 3 major classes, with tumour associated macrophages (TAMs) making up the majority (**Fig. 7A**).

**Figure 7:**
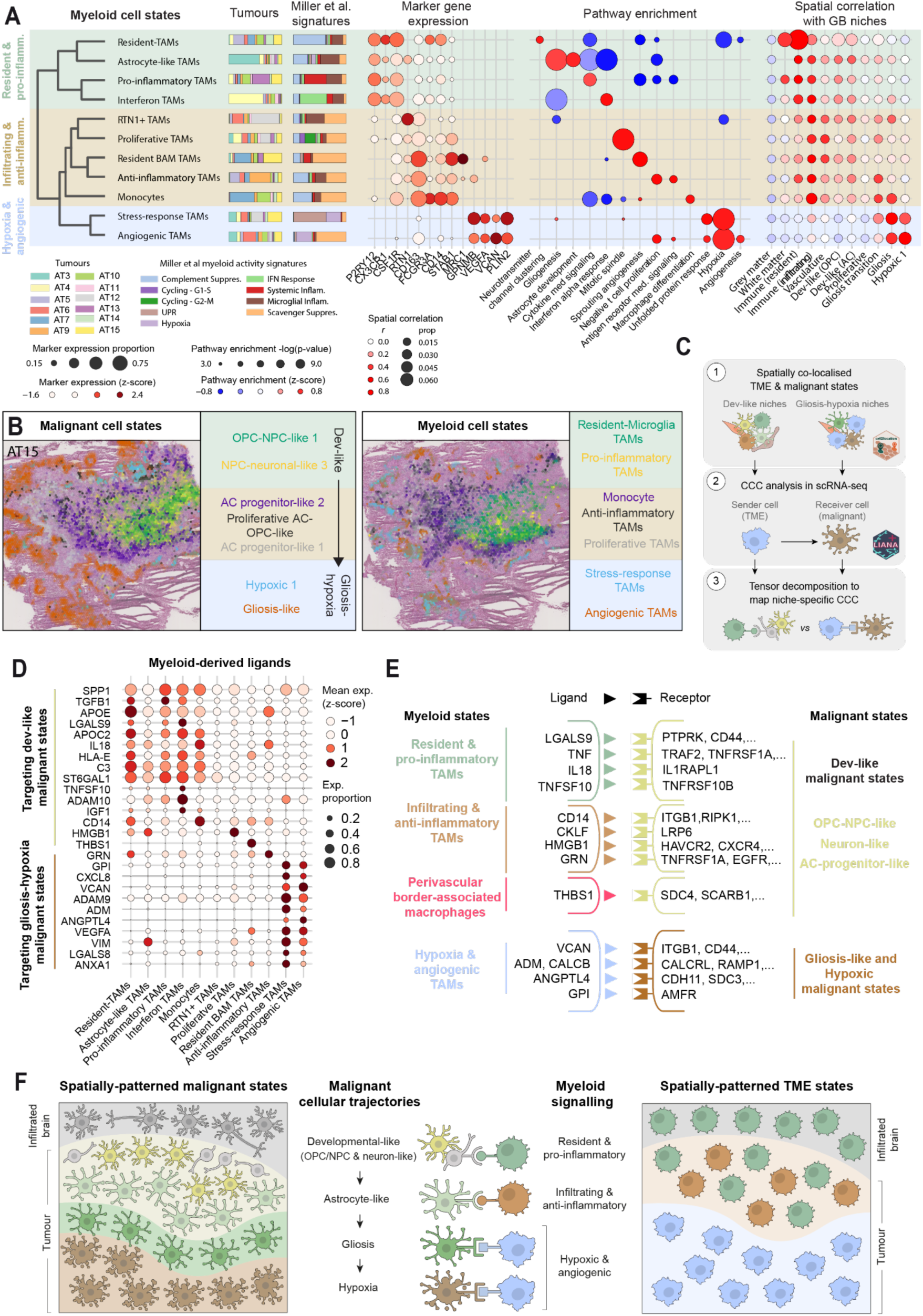
The malignant cell state trajectory is coupled to a regionalised myeloid TME and signalling. **A)** Myeloid heterogeneity in our multimodal atlas. Dendrogram depicts transcriptomic clustering of myeloid subcluster snRNA-seq profiles annotated to three major phenotypic classes (background colors). Stacked bar plots depict the relative distribution of myeloid states across tumours and the proportion of top myeloid activity signatures from Miller at al. per state. Dotplots show key marker gene expression and GOBP term enrichment across states. The spatial correlation dotplot depicts the sum cell2location abundance of a given myeloid cell state relative to total cell2location abundance across spots belonging to a given histopathological feature (prop) and the point biserial correlation between cell state abundances and histopathological feature (r). **B)** Spatial co-localisation of distinct malignant and myeloid states. Visium sections (left) show cell2location abundances of top cell states in tumour AT15 with colours mapping to corresponding labels highlighted in the boxes to the right. Malignant cell states are grouped according to their trajectory and myeloid cell states are grouped according to their phenotypic classes and their spatial organisation. **C)** Schematic detailing the spatially resolved cell-cell communication analysis strategy to detect signalling from distinct myeloid subtypes to their spatially co-localised malignant states that recur across multiple tumours. **D)** Myeloid ligands predicted to signal to distinct malignant cell states by spatially-resolved cell-cell communication analysis. Genes correspond to top ligands identified in myeloid signalling factors identified by tensor decomposition analysis. Dotplot represents the proportion of myeloid cells expressing the ligand (size) and relative expression across cell states (colour). **E)** Overview of major cell-cell interactions identified between regional myeloid (sender) and distinct malignant (receiver) cell states **F)** Summary diagram illustrating spatially organised myeloid signalling to malignant cell states across their cellular trajectory.

The first major class consisted of resident and pro-inflammatory myeloid cells, distinguished by expression of microglial marker genes (*P2RY12, CX3CR1*), and related gene sets and programs (**Fig. 7A**). Resident TAMs appeared closest to homeostatic microglia, whereas pro-inflammatory and interferon TAM populations showed elevated cytokine-mediated signalling and IFN-α signalling, respectively, associated with a strong anti-tumour response^57,58^.

The second class consisted of infiltrating and anti-inflammatory myeloid cells, distinguished by expression of monocyte-derived macrophage (MDM) marker genes (*CD163, STAB1, CD14*) (**Fig. 7A**). Monocytes expressed classical (*CD14*) and intermediate markers (*FCGR3A/CD16*), and macrophage differentiation gene sets. The remaining populations showed a scavenger immunosuppressive transcriptional phenotype as described by Miller et al^16^, including anti-inflammatory TAMs marked by negative regulation of T cells. Perivascular-border associated macrophages (L*YVE1*+ Resident BAM-TAMs) were included here based on their elevated expression of scavenger receptors (*MRC1*) (**Fig. 7A**)

The final class consisted of stress-response and angiogenic TAMs, marked by hypoxia and angiogenesis related gene expression (*VEGFA, VCAN*). These populations also expressed markers of “lipid-laden” macrophages (*GPNMB, PLIN2*) that were recently shown to reside in hypoxic compartments of GB tumours^15^ and promote proliferation of hypoxic malignant cells^15,59^.

Previous studies have identified spatial segregation of myeloid subtypes in GBs^16,17^. To precisely relate the spatial patterning of myeloid and GB cell states, we compared their distributions across GB tissue niches in our integrated snRNA-seq and Visium ST data (**Fig. 2B**). The three classes of myeloid cells established spatially distinct immune compartments across tumours (**Fig. 7B**). Resident and pro-inflammatory populations were enriched in the *Immune (resident)* niche (**Fig. 2B,C**, **Fig. 7A**), forming immune hotspots in the CT that spatially overlapped with dev-like malignant niches such as *Dev-like (OPC)* (**Fig. 2D,E, and Extended Data Fig. 19B-D**). These cells also extended into neighbouring white matter (**Fig. 7A**), consistent with their microglial phenotype. Infiltrating and anti-inflammatory populations formed the distinct *Immune (infiltrated)* niche in the CT (**Fig. 7A**, **Fig. 2B-D**) and were located in closer proximity to *Dev-like (AC)* and *Gliosis transition* niches (**Fig. 2E and Extended Data Fig. 19B-D**). Consistent with their infiltrating phenotype, these cells also occurred near *Vasculature*. Finally, stress-response and angiogenic TAMs were highly enriched in the *Gliosis* and *Hypoxia* niches in the PNZ, PAN and necrotic compartments (**Fig. 7A**, **Fig. 2B, and Extended Data Fig. 19B-D**), as previously described^15,59^.

These observations indicate distinct myeloid niches for dev-like and gliosis-hypoxia GB cell states. To determine whether these malignant states are exposed to distinct signals from their myeloid neighbours, we performed cell-cell communication (CCC) analysis in snRNA-seq data amongst spatially co-localised TME (including vasculature and lymphoid cells) and malignant states, assigning cells to either dev-like or gliosis-hypoxia niches (**Fig. 7C, Tables S11 and Extended Data Fig. 12**). We then used tensor decomposition to identify myeloid ligands that target receptors on either dev-like or gliosis-hypoxia malignant cells in a conserved manner across tumours (**Fig. 7C, Extended Data Fig. 20A-B, Methods).**

Our spatially refined CCC analysis identified distinct myeloid signalling environments across the GB cellular trajectory (**Fig. 7D,E, Table S13**). In dev-like niches, we predicted diverse myeloid signals targeting dev-like malignant states. TRAIL (TNFSF10), expressed predominantly by interferon TAMs, showed a putative interaction with TRAIL receptor 2 (TNFRSF10B) (**Fig. 7E and Extended Data Fig. 21A**) that can induce apoptosis in glioma cells^60^. Conversely, progranulin (GRN), highly expressed in anti-inflammatory TAMs, has been shown to promote glioma growth and therapeutic resistance ^61,62^. Several other predicted TAM ligands such as Interleukin-18 (IL18), Galectin-9 (LGALS9) and high mobility group box 1 (HMGB1) are well known regulators of tumour immune cells^63,64,65^, but how they influence malignant cells is not well understood.

In contrast, in gliosis-hypoxia niches, angiogenic and stress-response TAMs expressed ligands such as *VEGFA* and *VIM* that are known to be associated with hypoxia and angiogenesis in GBs^14,66^. These TAMs were also predicted to target gliosis-hypoxia malignant cells via expression of extracellular matrix proteins such as versican (VCAN) that has multiple roles in glioma^67^ including regulation of cell adhesion via β1 integrin (ITGB1)^68^. Additional TAM-expressed genes in these niches include wound healing regulators such as annexin-1 (ANXA1) and angiopoietin-like 4 (ANGPTL4), with the latter linked to wound healing through cadherin-11 (CDH11)^69^, and peptide hormones such as adrenomedullin (ADM) and calcitonin B (CALCB) that promote leukemia relapse through the CALCRL receptor^70^ (**Fig. 7D,E, and Extended Data Fig. 21A**).

In summary, signals from diverse TAM populations are linked to pro and anti-tumour functions across dev-like niches, whereas TAM signals are linked to wound healing, hypoxia and angiogenesis in gliosis-hypoxia niches. Taken together, we find that the myeloid TME is regionalised across the malignant cells trajectory and exposes malignant cells to distinct signals (**Fig. 7F**).

## Discussion

The extensive tumour heterogeneity and plasticity of GB has undermined efforts to pinpoint the cellular trajectories of cancer cells and develop targeted therapies. Here, we applied a holistic multi-modal mapping approach to jointly characterise the genetic, cellular and tissue architecture of GB tumours and dissect cancer cell trajectories. We created the most comprehensive GB atlas to date, combining large-scale single cell transcriptomics and epigenomics with spatial transcriptomics of each tumour as well as spatial whole genome sequencing.

Our findings show remarkable intratumour heterogeneity of GBs, extending previous surveys. Yet, our multimodal atlas and integrated analysis approach enabled us to trace GB heterogeneity to a spatiotemporally-patterned cancer cell trajectory from developmental-like towards gliosis and hypoxia states. This cellular trajectory is molecularly conserved across tumours, including multiple presumed GB subtypes, as well different subclonal lineages within each tumour. Furthermore, it is coupled to spatially patterned TME signalling from distinct myeloid populations. Hence, this trajectory represents a fundamental feature of cancer cell and TME organization across GB tumours.

Our multimodal atlassing approach facilitated the biological reinterpretation of spatiotemporal cancer cell states in GB. For example, using new transcriptomic and spatial characterisation, we found that the previously reported mesenchymal cell state^1,2^ was in fact a malignant cell gliosis and hypoxia response. Our integrative analysis demonstrated the stepwise transitions of malignant cells from developmental-like states resembling neuronal and oligodendrocyte progenitors into those resembling astrocytes, followed by the gliosis and hypoxia response. This trajectory is compatible with observations in mouse studies using GB patient-derived xenografts (PDX)^2,6^, yet we are the first human study to map this trajectory in patient tumours considering both intrinsic cancer cell and TME characteristics. Moreover, wound healing-like states and hypoxia are present across multiple cancers including pancreatic^71^ and colon^72^ cancer, suggesting the transitions of developmental-like cancer cells into these states may be a conserved feature across multiple solid tumours^73^. While hypoxia is a prominent feature of the GB cancer cell trajectory, whether other cell intrinsic and extrinsic cues push developmental-like cancer states into gliosis remains to be identified.

The spatiotemporal trajectory of cancer cells challenges the stratification of GB tumours into distinct subtypes. We captured remarkable regional heterogeneity of cancer cell states within each tumour. This suggests that GB tumour subtypes, commonly classified from single tumour biopsies, rather represent region-specific cancer cell states and tissue niches. We profiled one tumour in whole (AT15), ruling out such sampling confounders, and found it highly enriched for neuronal progenitor-like states. While this could represent a genuine tumour subtype as recently proposed^74^, we speculate that the temporal trajectory of cancer cells across tumour progression poses another challenge. For instance, tumours enriched for developmental-like or gliosis-hypoxia states may represent early or advanced stages of GB, respectively, possibly accounting for the worse survival outcomes associated with “mesenchymal” GBs^1^.

Previous studies have shown that genetic alterations can promote distinct cancer cell states across GB tumours^2^. While our results suggest that the GB malignant trajectory is shared across the GB genetic hierarchy, we also observed that cell states can be modulated across genetically distinct subclonal lineages, exemplified by the association of chr17 CN events with the neuronal progenitor-like states in one tumour. These results show that intratumour genetic heterogeneity can expand cell state diversity in GB. Furthermore, they imply that mutations can bias the GB malignant cell trajectory towards specific states, presumably through perturbations of key cancer cell state regulators.

We identify multi-scale spatial architecture of GB cancer cell subclones using complementary spatial whole genome sequencing and spatial transcriptomics approaches. We developed SpaceTree, a new computational approach to jointly map cancer cell states and clonal lineages in spatial transcriptomics data. These identified that, while GB subclones parcellate tumours into broad spatial domains, they finely spatially intermix within each domain in an intimate manner. This pattern contrasts with the spatial segregation of clones observed in breast^75^ and prostate^76^ cancer. We speculate this reflects the lack of physical barriers in the brain, such as ductal structures in the breast^75^, that can restrict clonal outgrowth, as well as the migratory nature of GB cancer cells.

Here, we defined GB cancer cell trajectories at a transcriptomic level. In a companion study, we analysed the joint single nuclei RNA- and ATAC-seq data generated in our atlas to define GB cancer cell states and their cellular transitions at the level of gene regulatory networks^20^. Beyond identifying transcriptional regulators and plasticity potential of distinct GB cell states, this orthogonal approach suggested highly specific transition routes between cancer states and validated the GB trajectory from developmental-like states towards gliosis and hypoxia.

We found that the spatial tumour niches and signalling of myeloid cells correlates with the malignant cell trajectory. These results present TME heterogeneity as a distinct axis of regulation for the broad spectrum of cancer cell states in GB. Whether these myeloid states, given their plasticity^16^, are induced by cues from cancer cells or represent independent drivers of tumour biology remains to be resolved. Nevertheless, our results predict highly regionalised myeloid functions tightly linked to cancer cell phenotypes as well as processes such as angiogenesis and immune privilege in GB.

Our study has several limitations. First, we do not validate the GB cancer cell trajectories using lineage tracing or experimental perturbations and instead focus on identifying conserved cellular patterns across human GB. Second, while our study examines the dominant cellular trajectories of cancer cells in primary GB, we do not examine their plasticity potential. Specifically, we do not demonstrate whether GB cancer cells can revert from gliosis and hypoxia into developmental-like states. Yet, we note that published lineage tracing studies using PDX models recapitulate the trajectory demonstrated here^2,6^ and our companion study predicts limited cell state transitions of gliosis-hypoxia cells based on gene regulatory networks^20^. Together, these observations suggest the possibility of limited cancer cell plasticity in GB beyond the trajectory illustrated here.

The significant intratumoural heterogeneity of GB underscores the challenges to therapeutic intervention. Our findings propose targeting the spatiotemporal trajectory of cancer cells and destabilising the myeloid TME associated with this trajectory, two conserved features of GB tumours identified here, as potential avenues for exploration.

## Data and code availability

The processed single cell and spatial cell atlassing datasets can be explored and downloaded from our interactive webportal (www.gbmspace.org/). The raw datasets will be deposited on EGA and the ST datasets will be deposited on the BioImage Archive. SpaceTree code is available at https://github.com/PMBio/spaceTree and tutorials are provided at https://pmbio.github.io/spaceTree/. ATAC datasets will be made accessible on the data portal upon peer review. All code necessary for re-analysis of the data presented in this paper is available upon request.

## Supporting information

Supplementary Computational Note

## Acknowledgements

We thank Jason Swedlow and Josh Titlow for their unwavering support and advice, Dave Adams, Muzz Haniffa and Aidan Maartens for comments on the manuscript, Koen Rademaker for assistance with Xenium data analysis, and Martin Prete for assistance with the GBM-space data portal. This work was supported by Wellcome Leap as part of the Delta Tissue programme (to J.B., D.H.R., S.B., O.S. and O.A.B.) and funded by the Wellcome Trust Grant institutional grants to the Wellcome Sanger Institute (206194 and 220540/Z/20/A), the Chan Zuckerberg Initiative DAF, an advised fund of the Silicon Valley Community Foundation (2023-323357, to O.S. and O.A.B.), a Wellcome Trust personal fellowship (223135/Z/21/Z, to S.B.) and investigator award (108139/Z/15/Z, to D.H.R), an ERC Advanced grant (789054, to D.H.R), the NIHR Cambridge Biomedical Research Centre (NIHR203312), the Medical Data Scientist Program of Heidelberg Faculty of Medicine, Heidelberg University (to A.S.R.), and the National Institute for Health and Care Research (NIHR) Academic Clinical Fellowship (to M. J.). J.S., M.J. J.B. were in part supported by the Francis Crick Institute, which receives its core funding from Cancer Research UK, the UK Medical Research Council and the Wellcome Trust (all under CC001051). The views expressed are those of the authors and not necessarily those of the NHS, the NIHR or the Department of Health.

## Author contributions

*Conception*: G.D.J., F.M., T.G., O.L., S.B., O.S., and O.A.B. conceived the study. *Data generation:* R.M. provided GB tissue samples, aided by T.S., A.Y. and H.B.. F.M. led experimental design, histology, Visium ST data generation and tissue processing for multiomic profiling. E.T, S.R., I.Mu. and M.P. contributed to histology and Visium ST data generation. T.G. coordinated multiomic profiling, led single nuclei multiome data generation, sample logistics and management. S.Ec., D.Z, A.O., H.P., I.Ma. Z.K. and E.P. contributed to single nuclei multiome data generation. O.G. led sequence library generation for multiome and Visium ST data, aided by Y.W., S.Ec. and T.P. M.D. generated LCM-WGS data. K.R. and A.L.T. generated Xenium data. J.D.BS. generated immunohistochemistry data. *Analysis/interpretation:* G.D.J. led multiome, ST and LCM-WGS data processing and analysis, and interpreted results. M.J. performed histopathological annotations of Visium ST data, validated by A.Q. and M.B. O.L. developed the SpaceTree model, applied the model to Visium ST data and interpreted results. A.A. and S.Er. performed cell2fate trajectory analysis and interpreted results. Q.Z. and A.S.R. contributed to myeloid cell state annotation and CCC analysis. H.M. analysed Xenium data. J.T.H.L. and S.M. assisted with raw Visium ST data processing. R.P. developed the GBM-space data portal for multiomic data with contributions from J.C.. O.G. coordinated raw data submission to data repositories. L.R., M.S. and M.M. contributed to interpretation of GBM cell states. *Supervision:* J.S.R. supervised CCC analysis. J.B. supervised histopathological annotation and immunohistochemistry validation. D.H.R. supervised myeloid data interpretation. S.B. supervised LCM-WGS data generation, analysis and interpretation. O.S. supervised SpaceTree model development, application and data interpretation. O.A.B. supervised cell atlassing data generation, analysis and interpretation. *Manuscript preparation*: G.D., F.M. and O.A.B. prepared figures. G.D. and O.A.B. wrote the manuscript with feedback from all authors.

## Extended data figures and tables

### Extended data figures

Extended Data figures 1-21 are provided in an accompanying file.

## Supplementary information

### Supplementary Computational Note

Supplementary note is provided in an accompanying file.

### Supplementary tables

Supplementary tables are provided in an accompanying file.

|  |  |
| --- | --- |
| Table S1 | Tumour sample and patient cohort summary |
| Table S2 | Data modality summary |
| Table S3 | Annotation hierarchy and associated cell counts |
| Table S4 | Malignant cell state distribution per tumour |
| Table S5 | Significantly enriched pathways and significantly up-regulated genes per granular cell state |
| Table S6 | Spapros marker gene identification of malignant and TME clusters |
| Table S7 | Recurrent spatial niche factor annotation |
| Table S8 | Curated Xenium probe panel addition |
| Table S9 | Cell state counts per tumour subclone |
| Table S10 | Cell2fate recurrent module annotation |
| Table S11 | CCC malignant niche overview |
| Table S12 | CCC malignant niche cell state identities |
| Table S13 | Myeloid ligands identified across dev-like and gliosis-hypoxia niches |
| Table S14 | Serial sectioning and multiomics plan of tumour tissue samples |

## Methods

### Human Subjects

Patients with suspected GB were identified pre-operatively and consented for entry into the study. Surgery was performed at Cambridge University Hospitals NHS Foundation Trust. Written and informed consent was obtained in accordance with the guidelines in The Declaration of Helsinki 2000. Ethical approval for the use of these tissues was obtained from the Cambridge Local Research Ethics Committee (REC 18/EE/0172). All patients underwent 5-ALA guided tumour resection as per local protocols. During tumour debulking, regions of high fluorescence were identified, their spatial location recorded and the tissue samples were collected for this study. Matched whole blood was taken during surgery for germline characterisation. The clinical characteristics of the patient cohort and samples are detailed in **Table S1**.

### Serial tissue sectioning for paired multi-omic analysis

Tissue was sampled from multiple sites of each GB tumour, targeting superior, anterior, posterior and middle regions where possible. Each tissue sample was immediately washed in saline buffer and embedded in OCT medium (Scigen OCT Compound, #4586) using a dry ice-cooled bath of isopentane at −75 °C. OCT-embedded samples were sectioned using a cryostat (Leica CX3050S). The fresh frozen tissue blocks were trimmed until the tissue surface was fully exposed. Two to three 10 µm thick sections were collected to check RNA integrity and another section was processed for Haematoxylin and Eosin (H&E) staining to assess tissue morphology. The RNA quality of each sample was evaluated by Tapestation (Tapestation RNA ScreenTape, Agilent) after isolating RNA (Qiagen EZ2 Automated RNA Extraction using EZ2 RNA/miRNA Tissue Kit Cat. No.959035). Only samples with RNA integrity number (RIN) values >7 were used for omic profiling. Blocks with good morphology and RIN were subsequently cryosectioned serially for different profiling applications as listed below (see **Table S14** for the sectioning plan of each tumour tissue block).

1. *Sectioning for single nuclei isolation*: A series of 50 µm thick sections, totalling 500 to 900 µm thick volume depending on the size of each tumour block, were collected in pre-chilled homogenization glass tubes and kept on dry ice until processing.
2. *Sectioning for Visium and Visium TO*: Sectioning for Visium ST was done according to the manufacturer’s guidelines. The Visium Spatial Gene Expression (GE) (catalog no. 2000233, 10x Genomics) slide was pre-chilled in a cryostat and four 10 µm sections were mounted onto the 0.42 cm^2^ capture areas. 2 sections were processed for Visium from each tumour block. For tumour samples larger than the capture areas, the blocks were scored and each half was placed on an individual capture area. Sections from two different blocks were collected on one Visium GE slide. Sectioning for Visium TO was done similarly, with four sections collected per block and two blocks per slide (catalog no. 3000394, 10x Genomics) were collected. Slides were then placed in a slide mailer and stored in a bag with desiccant in −80°C. Adjacent tissue sections were assessed with H&E staining.
3. *Sectioning for Visium CytAssist:* 10 µm cryosections were mounted on a Superfrost™ Plus Microscope Slide (Fisherbrand™) and stored in −80°C. Adjacent tissue sections were assessed with H&E staining.
4. *Sectioning for Xenium ST*: 10 µm cryosections were mounted onto Xenium slides (10x Genomics 3000941) and stored at −80°C for up to 4 weeks.
5. *Sectioning for LCM-WGS*: For LCM, sections were collected on a PEN MembranSlide (cat 11505158, Leica) and the slides were kept in the cryostat chamber until moved to −80°C. Adjacent tissue sections were assessed with H&E staining.
6. *Sectioning for Immunohistochemistry*: 10 µm cryosections were mounted onto Superfrost™ Plus Microscope Slides (Fisherbrand™) and stored at −80°C.

### H&E staining and imaging

Tissue sections were removed from −80°C and air dried before fixation in 70% ethanol for 3 minutes (min). After rinsing with deionised water, slides were stained in Gill II haematoxylin solution (Leica) for 7 seconds (s). Slides were completely rinsed in 2 washes of tap water for 20 and 25 s. Slides were then stained with aqueous eosin 1% (Leica) for 10 s and rinsed twice with deionised water for 20s and 25s, followed by dehydration through an ethanol series (70%, 70%, 100%, 100%; 20 s each) and cleared twice in 100% xylene for 10 s. Slides were coverslipped and allowed to air dry before being imaged on a Hamamatsu Nanozoomer 2.0HT digital slide scanner.

### Single-nuclei extraction

Nuclei were extracted from fresh frozen tissue sections that were homogenised using a glass Dounce homogenizer (Sigma) in nuclei isolation buffer (3mM MgCl2, 10mM NaCl, 10mM Tris (buffer pH7.4), 1 mM DTT, 0.1% Tween-20, 0.1% Nonidet P40, 1% BSA and 0.01% Digitonin) in the presence of Protector RNase Inhibitor (Roche) at 0.2 U/μl. Tissue was homogenised using 10 strokes with pestle A and then 10 strokes with pestle B. Nuclei were then filtered through a 40 μM filter, collected at 500 RCF and resuspended in 0.25 ml of storage buffer (PBS containing 1% BSA and Protector RNase Inhibitor (Roche) 1 U/μl). An aliquot of the nuclei suspension was incubated with Trypan Blue (Gibco 15250061) for counting and purified from debris using a Percoll gradient. The cleaned nuclei suspension was stained with Trypan blue and counted.

### 10x Genomics Chromium GEX and ATAC library preparation and sequencing

For the snRNA-seq experiments, two to three 10x reactions were prepared per tumour site and loaded onto the 10X chromium controller according to the manufacturer’s protocol for the Chromium Next GEM Single Cell Multiome ATAC + Gene Expression assay. Post-GEM-RT cleanup, cDNA amplification and 3′ gene expression library construction were carried out as per the Chromium Single Cell 3’ Reagent Kits v3 User Guide, to obtain between 5000-10,000 nuclei per reaction. Libraries were paired end-sequenced on a NovaSeq 6000 System (Illumina) using the Novaseq S4 Flowcell, targeting a minimum coverage of 100,000 read pairs per nuclei per modality. The following sequencing formats were employed for GEX and ATAC respectively:

- GEX: <u>Read 1</u>: *28 cycles* <u>i7 Index</u>: *10 cycles* <u>i5 Index</u>: *10 cycles* <u>Read 2</u>: 9*0 cycles*
- ATAC: <u>Read 1N</u>: *50 cycles* <u>i7 Index</u>: *8 cycles* <u>i5 Index</u>: *24 cycles* <u>Read 2N:</u> *49 cycles*

### Visium tissue optimisation

Tissue sections on TO slides were fixed in chilled methanol and H&E-stained according to the Visium Spatial Tissue Optimization User Guide (catalog no. CG000238 Rev A, 10x Genomics). Initial optimization experiments assessed 7 different permeabilization times at 6 min intervals and consistently indicated 12 to 30 min as optimal pre-treatment. Hence, subsequent TO experiments assessed 4 different permeabilization times (12, 18, 24 and 30 min) per tissue block, testing 2 blocks per TO slide. Brightfield histology images were taken using a 40× objective on a Hamamatsu Nanozoomer 2.0HT digital slide scanner and Fluorescent images were taken with a Cy3 filter using a 20× air objective and 200-ms exposure time on an Opera Phenix™ Plus.

### 10x Genomics Visium library preparation and sequencing

Libraries were generated according to the manufacturer’s instructions (Visium Spatial Gene Expression User Guide, CG000239 Rev A, 10x Genomics). Briefly, sections were fixed with cold methanol, stained with haematoxylin and eosin and imaged on a Hamamatsu NanoZoomer 2.0HT before permeabilization (most blocks processed for 12-24 mins), reverse transcription and cDNA synthesis using a template-switching protocol. Second-strand cDNA was liberated from the slide and single-indexed libraries were prepared using a 10x Genomics PCR-based protocol. Sequencing was performed on the NovaSeq 6000, aiming for a minimum of 50000 read-pairs per spot, with the following sequencing format; read 1: 28 cycles, i7 index: 10 cycles, i5 index: 10 cycles and read 2: 90 cycles.

### Visium CytAssist

Visium CytAssist slides were dried at RT for 5 min and fixed in cold methanol at −20°C for 30 min, then H&E-stained and imaged on Hamamatsu NanoZoomer 2.0HT. After destaining, human whole transcriptome probe pairs were hybridised and ligated to the tissue RNA. The CytAssist instrument was used to facilitate the transfer of transcriptomic probes from the standard glass slide to the Visium CytAssist Spatial Gene Expression Slide (v2, 11 mm capture area). Probe hybridization, probe ligation, release, extension, pre-amplification, and library preparation followed the Visium CytAssist Spatial Gene Expression Reagent Kits User Guide (CG000495).

Libraries were sequenced with paired-end dual-indexing (28 cycles Read 1, 10 cycles i7, 10 cycles i5, 90 cycles Read 2) on a Illumina NovaSeq 6000, aiming for a minimum 70.000 read pairs per spot. The Space Ranger v2.1.0 (10x Genomics) and the GRCh38-2020-A reference were used to process FASTQ files.

### Xenium Spatial Transcriptomics

Samples were processed using the Xenium Sample Preparation Kit (10x Genomics 1000460) as per the manufacturer’s protocols (CG000581 for pre-treatment including fixation with formaldehyde and permeabilization with SDS and methanol, followed by CG000582 for probe hybridisation and rolling circle product generation). Following autofluorescence quenching and nuclei counterstaining as per the manufacturer’s instructions (CG000582), slides were transferred onto a Xenium Analyzer instrument alongside Decoding Reagents (10x Genomics 1000461) and Decoding Consumables (1000487) all prepared according to the manufacturer’s protocol (CG000584). The DAPI-stained nuclei were exported from the instrument in the format of ome.tif images.

### Laser capture microdissection and DNA library preparation

Fresh-frozen tissue samples were fixed using PAXgene fixative (Qiagen) and embedded in paraffin. Sections were cut at 10 µm thickness, with additional reference slides constructed at 4 µm thickness to ensure the correct region of the tumour was identified. All slides were stained with H&E (Leica). Regions of interest were isolated by laser capture microdissection (LCM) and lysed using the Arcturus PicoPure Kit (Applied Biosystems) in accordance with the manufacturer’s instructions. Low-input, whole-genome sequencing libraries were created from each LCM lysate^77,78^. Sequencing was performed on the Illumina NovaSeq platform to generate 150 bp paired-end reads to a target coverage of 30X.

### Immunohistochemistry and microscopy

Fresh frozen GB tissue sections were fixed in 10% neutral buffered formalin (NBF) for 30 min. Slides were washed and incubated at 4°C for 10 min in solution of 2% H2O2 in methanol to block endogenous peroxidase. Slides were then washed and rinsed in 100% IMS and air dried before loading onto an automated staining platform (Leica Bond Rx). Antigen retrieval/stripping steps between each antibody were performed with Epitope Retrieval Solution 1 (pH9) (Leica, AR9961) for 20 min at 95°C. Slides were then incubated with 0.1% BSA-PBST solution for 5 min at RT for protein blocking.

For multiplexed immunofluorescence, primary antibodies (**Table 1**) were labelled with Opal reagents (Akoya Biosciences). Anti-rabbit Polymer HRP (Leica, RE7260-CE, RTU) was used as a conjugated secondary antibody. The antibody order, dilution and incubation time of primary antibody, secondary antibody and Opal dyes are detailed on **Table 2**. Slides were counterstained with DAPI (Thermo Scientific, 62248; 1:2500), mounted with Prolong Gold Antifade reagent (Invitrogen, P36934), and scanned following the PhenoImager whole slide workflow, Akoya PhenoImager HT (formerly Vectra Polaris) using PhenoImager HT 2.0 software.

**Table 1:**
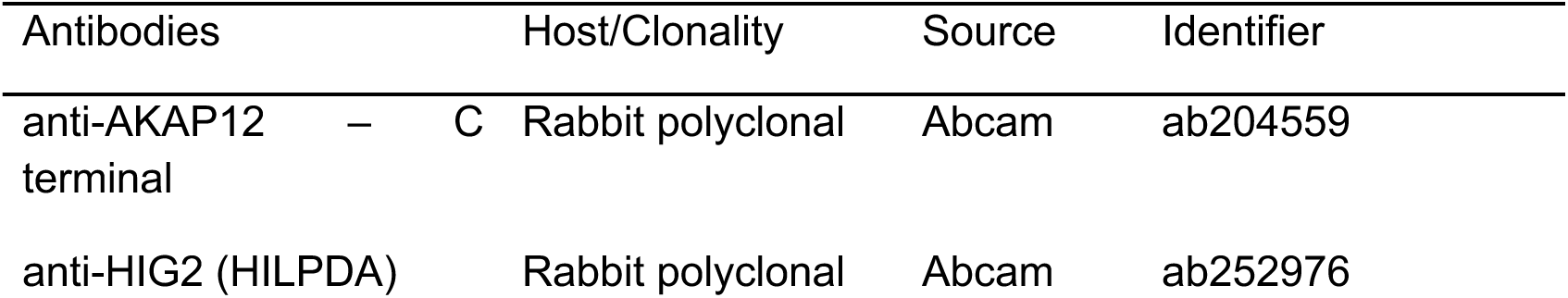
Primary antibodies.

**Table 2:** Multiplex immunofluorescence antibody parameters.

| Position | Ab | AR | Primary dilution | Primary incubation (min) | Secondary | Secondary incubation (min) | Opal | Opal dilution | Opal incubation (min) |
| --- | --- | --- | --- | --- | --- | --- | --- | --- | --- |
| 1 | AKAP12 | ER1 20 | 1 in 750 | 15 | Rabbit Polymer | 8 | 690 | 1 in 150 | 10 |
| 2 | HILPDA | ER1 20 | 1 in 400 | 15 | Rabbit Polymer | 8 | 520 | 1 in 300 | 10 |

Diluents (Akoya)

OPAL diluent – 1xPlus Automation Amplification Diluent cat number FP1609 Antibody diluent - 1X Antibody Diluent / Block, 100ml cat number ARD1001EA

### Pathological annotation

A pathologist annotated digitised images of the H&E stained tissue sections with histologically- distinct anatomic features. Histological annotations were defined according to the IVY Glioblastoma Atlas Project reference; an existing taxonomy system with established inter- neuropathologist agreement and clinical relevance^26^. Histological annotations included:

1. *Leading edge (LE)*: the outermost boundary of the tumour where the laminar architecture of the cortical layers is frequently evident.
2. *Infiltrating tumour (IT)*: the intermediate zone between the leading edge and cellular tumour, frequently marked by perineuronal satellitosis.
3. *Cellular tumour (CT)*: the major part of the tumour core.
4. *Necrosis*: dead or dying tissue, marked by the presence of karyorrhectic or cellular debris and the absence of crisp cytological architecture.
5. *Pseudopalisading cells around necrosis (PAN)*: the narrow boundary of cells arranged like pseudopalisades along the perimeter of necrosis in the tumour core.
6. *Perinecrotic zone (PNZ)*: a boundary of tumour cells in the tumour core along the edge of necrosis that lacks a clear demarcation of pseudopalisading cells around necrosis.
7. *Microvascular proliferation (MVP)*: two or more blood vessels sharing a common vessel wall of endothelial and smooth muscle cells typically in the tumour core, and arranged in the shape of a glomerulus or garland of multiple interconnected blood vessels.
8. *Hyperplastic blood vessels (HPV):* blood vessels with thickened walls found anywhere in the tumour.

Two additional pathologists independently validated the histological annotations.

### Single nuclei multiome data processing and quality control

We aligned reads from each snRNA-seq and ATAC-seq library to a custom-made genome consisting of 10X Genomics’ GRCh38 3.0.0 pre-mRNA reference genome and 10X Genomics Cell Ranger ARC 2.0.1 ATAC genome. To perform quantification and initial quality control, we used the default parameters in the Cell Ranger ARC software (v2.0.1; 10X Genomics)^79,80^. This was followed by CellBender^81^ (v0.2.0), which was applied to the Cell Ranger output to correct for background noise and identify empty droplets. The parameters --expected-droplets and --total-droplets-included were identified based on sample UMI curves. Quality control of the RNA data was performed using Cell Ranger ARC filtered count matrices with Scanpy^82^ (v1.9.3). This involved removing nuclei with total gene counts <500, total counts UMI <1000, and nuclei with >10% of reads mapping to mitochondrial content. We then applied Scrublet^83^ (v0.2.3) on each library individually and filtered our data based on a two-step method adapted from previously described median absolute deviation (MAD) thresholding. Individual cells were first filtered based on methods described in Pijuan-Sala et al.^84^ (FDR < 0.05) and then clusters were subsequently filtered using similar methods^84,85^ to ensure the removal of probable subthreshold doublets (FDR < 0.1). Following doublet detection and removal, any remaining cell with a total UMI count >75,000 was removed from the dataset. Finally, we calculated cell cycling scores using Scanpy based on the gene list described by Tirosh et al.^86^. ATAC data were processed according to the ArchR^87^ workflow (minimum fragments >1000 and >4 transcriptional start sites). These barcodes were then filtered by the list of barcodes passing RNA quality control filters, effectively removing the majority of low-quality ATAC barcodes. Finally, the RNA data was subset by barcodes passing ATAC filters to ensure symmetry between modalities for downstream applications.

### snRNA-seq integration

snRNA-seq libraries were concatenated into a single dataset and log-normalised for highly variable gene (HVG) selection. HVGs were identified with Scanpy using tumour ID as a batch key and dispersion-based feature selection with following non-default parameters: (min_mean=0.0075, max_mean=4, min_disp=0.1). The resulting list of 10,701 HVGs was then used for batch integration with scVI^21^ (v0.19.0), for which we used each 10x reaction as the batch and included tumour ID, tumour site, 10x reaction date, and cell cycle phase as model covariates. Model hyperparameters included 50 latent space dimensions and 2 hidden layers, with 1024 nodes per layer. After training the model, leiden clustering was performed using the nearest neighbours graph constructed from the resulting scVI embeddings. Leiden clusters corresponding to putative and broadly defined TME populations were then annotated (e.g. oligodendrocytes, myeloid cells, neurons, lymphocytes) based on marker gene expression.

### snRNA-seq CNA inference

The detection of cell-specific copy number aberrations (CNAs) was performed on each tumour individually using inferCNVpy (https://github.com/icbi-lab/infercnvpy), a python implementation of the inferCNV^88^ method, with parameters consistent with previous GB studies^89^ and a window size of 250 genes. InferCNV also requires a set of non-malignant reference cells, for which we used the least ambiguous TME populations identified based on marker gene expression: macrophages, oligodendrocytes, neurons, and lymphocytes.

### Clustering and cell state annotation

Following integration and coarse-grained clustering, the first step in atlas annotation was accurately delineating between malignant and non-malignant cells. This followed an approach adapted from previous GB studies^2^, in which two separate CNA metrics were used to score cells: CNA signal and CNA correlation. We calculated the absolute CNA signal by averaging absolute CNA scores originating from the diagnostically relevant chromosome 7 gain and chromosome 10 loss events. CNA correlation was computed by calculating the correlation between the CN profile of a given cell and mean CN profile of cells identified as non-malignant through marker gene expression. Malignant cell signal was defined as a CNA signal > 0.02 and a CNA correlation > 0.3. As static malignant cell thresholds may not identify all malignant cells and are dependent on the coverage in expression-based CN inference methods, we then annotated clusters as putatively malignant on the basis of the percentage of malignant cells. The majority of malignant clusters exhibited a malignant percentage > 20% but malignant clusters with a percentage ≥ 3% were flagged as putatively malignant.

We then reintegrated and sub-clustered each major TME population and all malignant cells, individually. This was followed by a subsequent CNA evaluation to ensure there were no misannotated subclusters. TME cell types were then identified manually using lists of well-established human brain marker genes (**Extended Data Fig. 6A**). TME subclusters with a malignant cell percentage ≥ 3% were also flagged. Malignant cells subclusters were annotated using composite of four different approaches: (1) We calculated a series of gene module scores based on the workflow outlined by Neftel et al.^2^ in conjunction with the *score_genes* function in Scanpy, which is analogous to their methodology. (2) We ran mapped our GB data onto a developing brain atlas^22^ using scPoli^90^ with default parameters to identify developing brain analogs to malignant cell state identities. (3) Markers specific to each cluster were identified using scanpy.tl.rank_genes_groups based on Wilcoxon rank-sum. (4) Gene set enrichment analysis was performed on differential gene expression results using decoupler^91^ with MGSigDB signatures^92,93^.

### Visium spatial cell type mapping and tissue niche identification

Visium spatial RNA-sequencing data were mapped to the official Cell Ranger GRCh38 using 10x Genomics Space Ranger Software Suite (v2.0 for standard Visium, v2.1 for Visium Cytassist). To spatially map malignant and TME cell states in Visium data, we used the cell2location model^25^ to deconvolve the Visium spot transcriptomes to cell state abundances based on paired snRNA-seq reference signatures. We generated unique reference signatures for each tumour to better reflect malignant cell gene expression heterogeneity. This was done using the negative binomial regression model packaged with cell2location and included the following changes to default training parameters: max_epochs=400, batch_size=10000, and lr=0.002. The cell2location model was run on each individual tumour (i.e. batching multiple Visium sections) with the following modifications to default parameters: model parameters were adjusted to N_cells_per_location=30 and detection_alpha=200; training parameters adjustments included max_epochs=6000, and a batch_size set to approximately 25% of the total number of spots profiled for a given tumour.

To identify tissue niches, we applied the scikit-learn NMF model packaged with cell2location to our cell state abundance results^25^. This allowed us to decompose matrices of inferred cell state abundances into components representing patterns of cell state co-localisation, analogous to microenvironmental niches. The model was trained using 16 factors for each individual tumour. Of the resulting matrices, we normalised the matrices composed of factor loadings per cell state and performed a cross-tumour annotation of associated factors. Factors were grouped into meta-factors, described as niches, using agglomerative hierarchical clustering with cosine distance. These grouped factors were annotated according to the most frequently observed cell states in each factor, in addition to the correlation between cell state and factor abundances.

To define a measure of spatial intermixing we calculated Shannon entropy based on cell2location abundance frequencies for each Visium spot. To ensure spots were highly representative of a given cell state, only spots in the 98^th^ percentile of abundance for each cell state were considered.

To quantify the relative spatial organisation of cell states, niche factors, and cell2fate modules, we calculated the minimum pairwise spot distance between all component pairs for each of these features independently using k-d trees. As the inferred abundance of these features are continuous variables, we imposed thresholds based on feature abundance to quantify spots associated with high feature abundance. This included two filters: abundance in a spot must be greater or equal to the median abundance of a feature, and, to remove abundances with poor signal, the abundance must also be greater than a base threshold. This allowed us to focus on comparing regions of high density while also accounting for biologically relevant differences in abundance. The base threshold varied depending on the application and was based on the distribution of abundances (cell2location = 4, niche factor = 10, and cell2fate modules = 2). As some spatial features have inherently different spatial distributions – e.g. the compactness and density of vasculature compared to malignant niches – we summarised these distances by calculating the bottom 25^th^ percentile of nearest neighbour distances.

### Xenium panel design and data processing

Our Xenium panel included an additional 62 probes alongside the standard 266-gene Xenium human brain panel (https://www.10xgenomics.com/products/xenium-panels). The additional custom probes were largely selected by applying a probe selection method, Spapros^94^, to our snRNA-seq data. Spapros was run without selecting a feature number cutoff (n=None) and default parameters. Malignant and TME cells were run inclusively to ensure useful cell state delineation; however, the atlas was downsampled for each tumour and cell state to reduce the computational burden. We then selected high-quality markers based on the spatial expression according to our Visium data. For each cell state associated with a given marker, we calculated Pearson correlations between the marker voxel expression and cell state abundance inferred via cell2location. Markers with the highest median correlation and strongest inter-tumoural concordance were selected. Markers that were already present in the human brain panel were omitted. This resulted in 44 Spapros-derived cell state markers which were further supplemented by 17 known GB markers which were expressed in our Visium data.

Default on-instrument nucleus segmentation and expansion was performed on Xenium datasets (Xenium Onboard Analyser versions xenium-1.5.0.3 and 1.6.0.8). Nuclei were segmented from DAPI images, followed by the definition of cell boundary as either a 15 µm expansion or the presence of another cell boundary. Single cell Xenium gene expression data were pre-processed using standard Scanpy workflow^95^. Cell-by-gene matrices were filtered to exclude cells with low numbers of transcripts (total gene count < 10) and genes that expressed in fewer than 10 cells across the dataset. Gene counts were normalised and log-transformed prior to downstream analysis. Tangram^27^ (version 1.0.4) was used to transfer coarse cell state labels from our snRNA-seq atlas onto Xenium sections using default parameters. Patient matched snRNA-seq profiles were used as reference for each Xenium dataset. Intersecting and non-zero expressed genes from both modalities were used as training sets (315 genes). Tangram was trained with default learning rate (0.1) for 500 epochs, in ‘clusters’ mode with a uniform density prior, and most probable cell type labels were transferred to Xenium cells from output mapping.

### Joint snRNA- and snATAC-seq subclone calling

We leveraged multiome data to jointly infer subclones from multiome data. For snRNA-seq, we used the inferCNV results described above. For snATAC-seq data, we detected CNs directly on the Cell Ranger ARC ATAC output using epiAneufinder^96^ with a 1MB window size and blacklisted regions of hg38 provided by the authors. Cells with total fragment counts <20,000 were filtered to ensure sufficient chromosomal coverage.

Clones were called by applying leiden clustering on a neighbourhood graph generated from genome-wide CN calls for each modality. We observed that scATAC-seq-based CNAs exhibited greater variability than RNA-based CNAs, likely due to the higher sensitivity of ATAC-seq in detecting open chromatin regions, resulting in a larger number of events. To be more conservative and reduce false positives, we matched clones from scATAC-seq to scRNA-seq clones. To reduce clone redundancy, we assessed the similarity between scATAC-seq clones using correlation distance. Clones with pairwise distances below a predefined threshold were merged to eliminate highly similar clones.

To mitigate differences in resolution between modalities, we applied interpolation to the RNA-based CNV profiles, aligning them with the larger genomic windows of scATAC-seq data. For matching clones across modalities, we computed two metrics: cell overlap and cosine similarity between cluster centroids. Cell overlap was defined as the proportion of cells shared between scRNA-seq and scATAC-seq clones, while cosine similarity was calculated based on the mean CNV profile for each clone. These metrics were averaged to create a composite similarity score.

Next, we retained the top 20% of clone matches based on this composite similarity score. For each RNA clone, we greedily selected the most similar scATAC-seq clones, assigning cells shared between matched RNA and ATAC clones to a new joint clone identifier. Finally, we applied label spreading to propagate clone labels to classified RNA cells based on their CNV profiles. Each cell was assigned a probability of belonging to a clone, allowing flexibility in downstream analyses by using different confidence thresholds. For most analyses, we applied a confidence threshold of 0.9 to ensure high-confidence clone assignments. Overall, the methodological concordance with LCM-based WGS orthogonally validated the efficacy of counts-based methods at identifying core clone populations. Furthermore, while there may be minor clones with small artifactual signals introduced by inferential nature of this method, we reasoned that redundant granularity would allow us to conservatively control for genetic heterogeneity when applied to downstream trajectory analyses.

### LCM WGS data variant calling, filtering and alleleIntegrator analysis

Following sequencing, all low-input WGS data was mapped to the hg38 reference genome using BWA-MEM^97^. All mutations were called against matched normal blood samples using the Wellcome Sanger Institute variant calling pipeline.

Copy number variants were called using ASCAT^98^ and Battenberg^99^. Battenberg copy number results were used for subsequent analyses. For each patient, LCM libraries with sufficient DNA concentrations (10 μg/mL) and copy number diversity were used as the basis for alleleIntegrator^38^. CNAs were called in the snRNA-seq data using alleleIntegrator. The standard workflow was applied using Battenberg CN segments alongside the standard alleleIntegrator workflow. Clones were identified by first subsetting cells with support for a chr7 gain, which is diagnostically relevant to GB, followed by support for putative lineage-specific CNAs (maxPostProb > 0.7).

CaVEMan^100^ was used to call substitutions. Variants were filtered to account for artefacts consistent with other low-input WGS studies^37,78^, which involved applying a filter designed to remove false positive variants such as erroneously processed cruciform DNA (https://github.com/MathijsSanders/SangerLCMFiltering). Subsequently, variants were retained if they met a minimum median alignment score (ASMD) of ≥140, fewer than half reads were clipped (CLPM = 0), were supported by at least 4 variant reads, and had a total sequencing depth of 10x or greater. The minimum thresholds for mapping quality and base quality were set to 30 and 25, respectively. Finally, we used a beta-binomial model adapted from the Shearwater variant caller by Coorens et al.^101^ designed to remove both germline variants and consistently observed low-quality variants. Site-specific error rates were calculated for each variant and only retained if the FDR-correct *p*-value was less than 0.001. Sample phylogenies were generated using Seqouia^102^ with default parameters and no application of the binomial mixture model. To ensure Sequoia compatibility with our tumour data, substitutions in CN variable regions were omitted.

Following filtering, PyClone-VI^39^ was used to cluster variants outside of patient-associated CNAs into putative subclones for each patient, separately. Each patient was run with the following non-default parameters: number of grid points = 150, number of restarts = 100, number of maximum expected clusters = 30. Relationships between inferred clone populations, including phylogenetic analysis, was performed using PairTree^103,104^ with, apart from default parameters, a burn-in of 0.4 and the RPROP method for fitting subclones to trees. LCM samples were grouped into clusters based on clone population frequencies with agglomerative clustering using Bray-Curtis dissimilarity and complete linkage.

### Spatial clone deconvolution with SpaceTree

To assess the spatial distribution of multiome-inferred clones (i.e. called from joint RNA and ATAC) in Visium data, we developed the SpaceTree model for joint clone and cell state deconvolution (**Supp. Comp. Methods**). This method follows a multi-step process, beginning with graph construction and followed by the use of a multi-task GNN to predict labels for each Visium spot. We used cell2location cell state mapping results to further evaluate the performance of SpaceTree, showing a high degree of concordance (see **Supp. Comp. Methods**, section 2.5.3). Additional orthogonal validation of this method was performed through comparison with LCM-derived analyses, including spatial clone cluster concordance (**Fig. 5C**). Cluster concordance evaluation involved summarising SpaceTree voxel probabilities that were histologically consistent with LCM ROIs into equivalent ROIs. These were hierarchically clustered using the same methodology as LCM samples (agglomerative clustering with Bray-Curtis dissimilarity and complete linkage). Tumour purity was calculated according to SpaceTree clone mapping results and represents the sum probability of all malignant clones in a given Visium spot. Shannon entropy was calculated on clone probabilities per spot and defined more generally as clonal entropy.

### Cell2fate cell trajectory analysis

Cell fate trajectories were inferred using a Bayesian generative model, cell2fate^42^ from snRNA-seq data with default parameters. This method infers temporal expression modules resulting from the factorisation of RNA velocity data and then applies associated models of gene activation to infer cell-specific time along a trajectory. A key advantage of the Bayesian approach is that this method also returns posterior uncertainty estimates for every parameter, allowing for the evaluation of the confidence of cell state transitions. As input, we generated spliced and unspliced counts using STARsolo^105^ v2.7.10a. Cell2fate analysis was run for each tumour clone separately using 10x reaction as a batch covariate, with 20 minimum shared genes, and 3000 HVGs in total. Individual cell2fate runs were evaluated first based on maximum cell-specific time (maximum(Tc) > 20), and then flagged as high-quality if average cell state transition scores exceeded 0.25. To quantify cell state transition scores, posterior distributions of cell-specific times were compared between two cell states, with the score equalling to the percentage of cells in state B that have a greater inferred time than the 90^th^ percentile of cells in state A (**Extended Data Fig. 17A,C**) . For applications involving inter-clonal comparison across tumours, inferred times were min-max normalised.

RNA velocity modules were subsequently hierarchically clustered using Jaccard distance of the top 200 genes in each module for the purposes of inter-clone module annotation. The “meta”-modules were then manually annotated according to (1) the identity of shared top genes, (2) the results of over-representation analysis of MGSigDB signatures for this gene list, and (3) enrichment of a given meta-module across malignant cell state annotations. Meta-module markers were compiled by taking the top 50 genes associated with each module, and sorting them by the degree of overlap across all modules in a given meta-module.

Using the standard cell2location workflow and the methods outlined in Aivazidis et al^42^, we also spatially mapped RNA velocity modules from cell2fate for a subset of tumours. Sections were selected if the multiome clone associated with the cell2fate run had a frequency of at least 30% in the associated site of sampling.

### Survivorship analysis of bulk RNA-seq samples

Count matrices from CPTAC^53^ and TCGA^54,55^ bulk RNA-seq samples were downloaded from the Genomic Data Commons (GDC). Samples were aggregated and log-normalised using Scanpy^82^. Relative expression scores were calculated for each meta-module based on commonly observed top genes (n=200) and samples were annotated as “enriched” for a meta-module if the expression scores were greater than the 75^th^ percentile. Kaplan-Meier analysis followed by the log rank test were performed using the python package lifelines^106^ between meta-modules with early and late inferred times.

### Cell-cell communication analysis

We performed two sets of cell-cell communication (CCC) analysis to predict TME-derived ligands targeting receptors expressed on malignant cell states across different dev-like versus gliosis-hypoxia niches, focusing on myeloid signalling.

In step 1, we identified TME cell states spatially co-localised with dev-like versus gliosis-hypoxia malignant cell states from Visium ST data (**Fig. 2**). These included the entire GB TME including myeloid, vasculature and T cells, and was summarised across all tumours (**Table S11**). We then selected malignant cell states present within dev-like versus gliosis-hypoxia niches per each individual tumour (**Table S12**).

In step 2, we performed CCC analysis using the LIANA+ framework (version 1.2.0)^107^, which integrates multiple computational methods and prior knowledge resources into a consensus-based approach to identify ligand-receptor interactions. We employed the “consensus” approach, combining five different methods implemented in LIANA+: CellPhoneDB, Connectome, log2FC, NATMI, SingleCellSignalR, and CellChat. Consensus scores across these methods were calculated using the robust rank aggregation method^108^, with an ‘expr_prop’ (filtering criteria based on the proportion of cells per cell type) value of 0.05 and ‘n_perms’ (number of permutations) set to 1000. Here, TME cell states were set as the sources (ligands) and the malignant cell states as the targets (receptors). An expression threshold of 0.05 was applied to filter out ligands and receptors below this threshold. We applied the LIANA analysis separately to the dev-like and gliosis-hypoxia niches.

In step 3, the resulting ligand-receptor pairs were analyzed using the Tensor-cell2cell method^109^ (version 0.7.4), which utilised the LIANA+ results as input to identify shared CCC events across tumours as factors. This method automatically determined the number of factors using elbow analysis and employed robust tensor factorization to ensure consistent results across donors. The main parameters and their values used in this analysis were as follows: ‘score_key = magnitude_rank’ (the LIANA+ score used for factor analysis), ‘tf_optimization = robust’ (the robustness setting for tensor factorization), ‘upper_rank = 15’ (the upper limit for the number of factors), and ‘tf_svd = numpy_svd’ (the singular value decomposition method used). We then identified factors shared across tumours that involve myeloid cells, specifically Factors 1 and 2 from the dev-like analysis and Factor 1 from the gliosis-hypoxia analysis (**Extended Data Fig. 20**).

In step 4, we sorted cell interactions associated with each factor based on their loadings and selected interactions in the top 50^th^ percentile. We then focused on ligands from the myeloid populations that are predicted to target receptors on their respective malignant cell states. We filtered the ligand-receptor pair list to ensure at least one relevant cell state showed mean gene expression ≥ 0.5 and had a total expression fraction ≥ 0.1 for each associated gene pair. This resulted in the identification of 80 unique interactions in the dev-like niches (involving 37 ligands), 35 unique interactions in the gliosis analysis (involving 18 ligands), and a total of 63 shared interactions shared across both niches (involving 28 ligands).

