## Extended Data figures (1-21) for "A spatiotemporal cancer cell trajectory underlies glioblastoma heterogeneity"

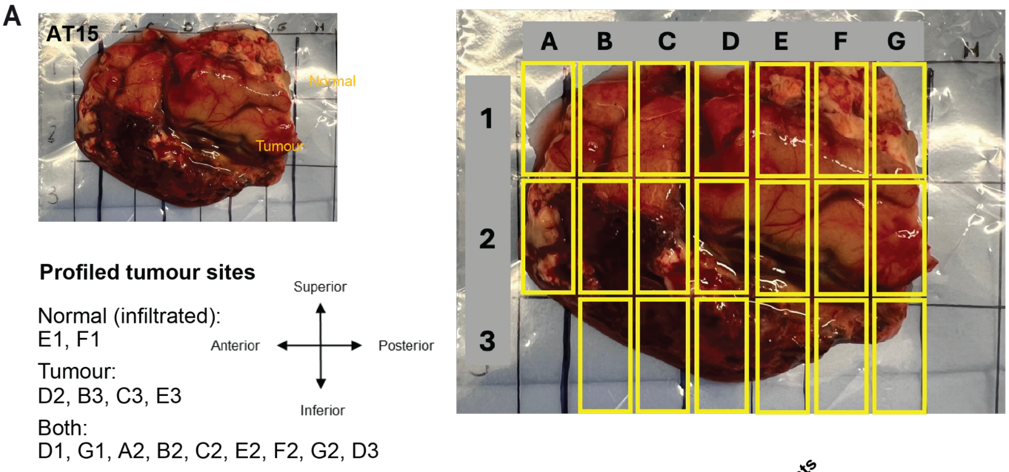


B

| Tumour | Age | Sex | MGMT methyl. | Survival | Total sites | Multiome sites | Nuclei counts | Visium sites | Visium spot counts | Xenium sites | Xenium cells | Spatial WGS profiles |
|---|---|---|---|---|---|---|---|---|---|---|---|---|
| AT3 | 62 | M | 43% | 7 | 4 | 4 | 76845 | 4 | 14589 | 0 | 0 | 35 |
| AT4 | 54 | M | 2% | 7 | 4 | 4 | 83500 | 4 | 20638 | 1 | 126738 | 0 |
| AT5 | 82 | F | 5% | 4 | 4 | 4 | 65665 | 4 | 16566 | 0 | 0 | 0 |
| AT6 | 64 | F | 38% | 5 | 4 | 4 | 93636 | 4 | 19695 | 1 | 547340 | 0 |
| AT7 | 59 | M | 1% | 16 | 4 | 4 | 45884 | 4 | 23322 | 0 | 0 | 0 |
| AT9 | 62 | M | 2% | 6 | 4 | 4 | 130063 | 4 | 19792 | 2 | 632741 | 0 |
| AT10 | 70 | M | 20% | 4 | 4 | 4 | 85983 | 4 | 26183 | 4 | 1138158 | 26 |
| AT11 | 63 | M | 35% | 5 | 4 | 4 | 52538 | 4 | 33252 | 0 | 0 | 0 |
| AT12 | 58 | F | 42% | 10 | 4 | 4 | 47065 | 4 | 29587 | 1 | 269486 | 0 |
| AT13 | 73 | F |  | 1 | 4 | 4 | 119787 | 4 | 30443 | 2 | 180851 | 0 |
| AT14 | 68 | F | 2% | 5 | 2 | 2 | 32488 | 2 | 13387 | 1 | 164489 | 0 |
| AT15 | 64 | M | 2% | 13 | 15 | 15 | 191875 | 7 | 129695 | 5 | 1293269 | 0 |

***Extended Data Fig. 1: Multi-region multi-modal profiling of GBs.***
***A)*** Overview of grid-like sampling strategy for tumour AT15.
***B)*** Summary of each data modality.

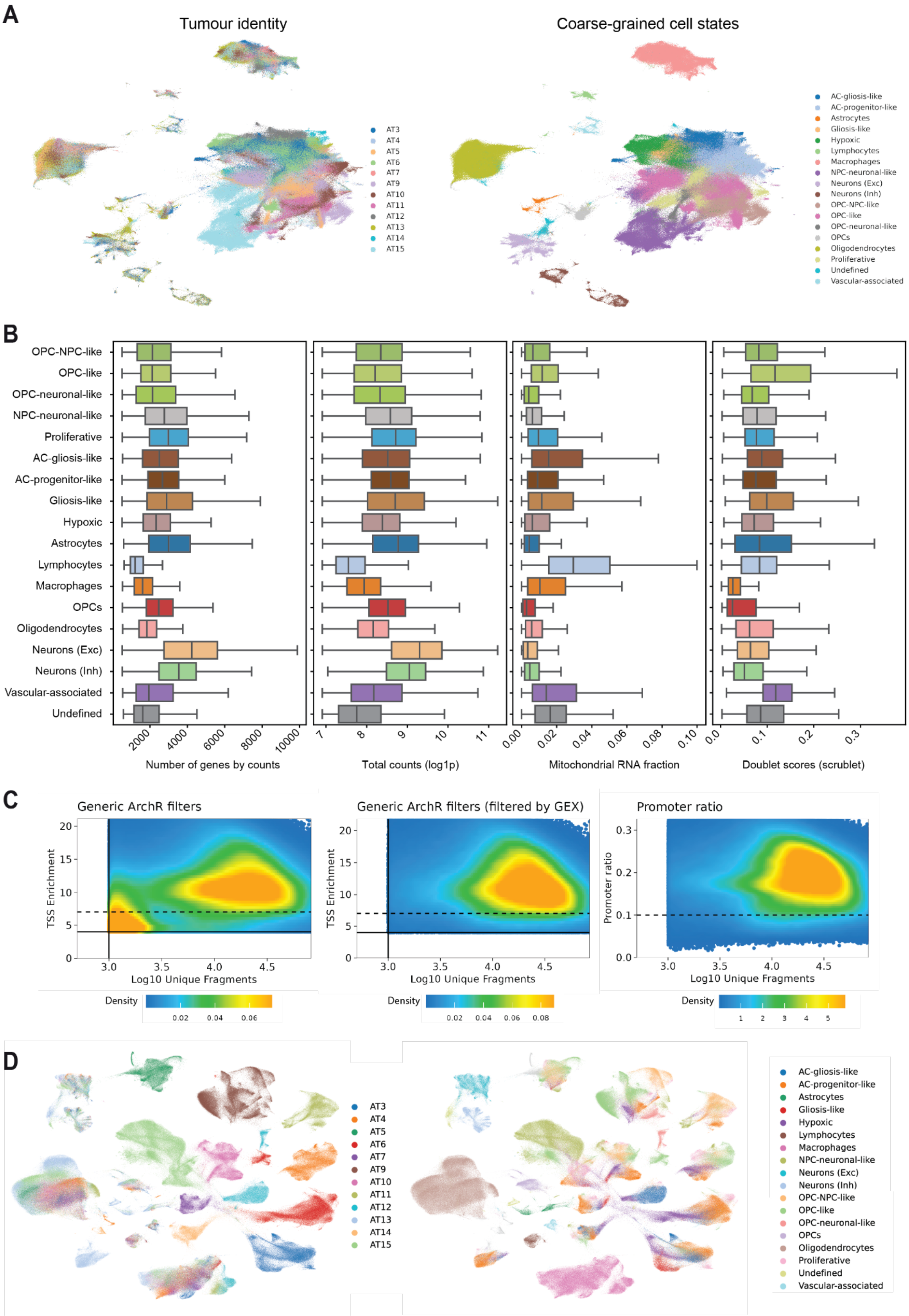


***Extended Data Fig. 2: Summary of snRNA and ATAC QC results.***

***A)*** UMAPs derived from scVI embeddings indicating tumour identity (left) and coarse-grained cell state identity (right) across the main RNA atlas.

***B)*** Distributions of common RNA QC metrics (number of genes by counts, total counts, mitochondrial read fraction, and doublet scores) post-filtering across major coarse-grained cell states.

***C)*** Summary of ATAC QC metrics and applied QC filters. From left to right: nuclei passing generic ArchR filters, nuclei passing both generic ArchR filters and RNA filters, and promoter ratio compared against minimum fragments.Generic ArchR filters indicated by solid lines (minimum unique fragments $\geq$ 1000 and TSS enrichments $\geq$ 4). Dashed lines represent target minimum values for the respective metrics.

***D)*** UMAPs showing the topology of tumour (left) and coarse-grained cell state (right) identities without the application of integration methods.

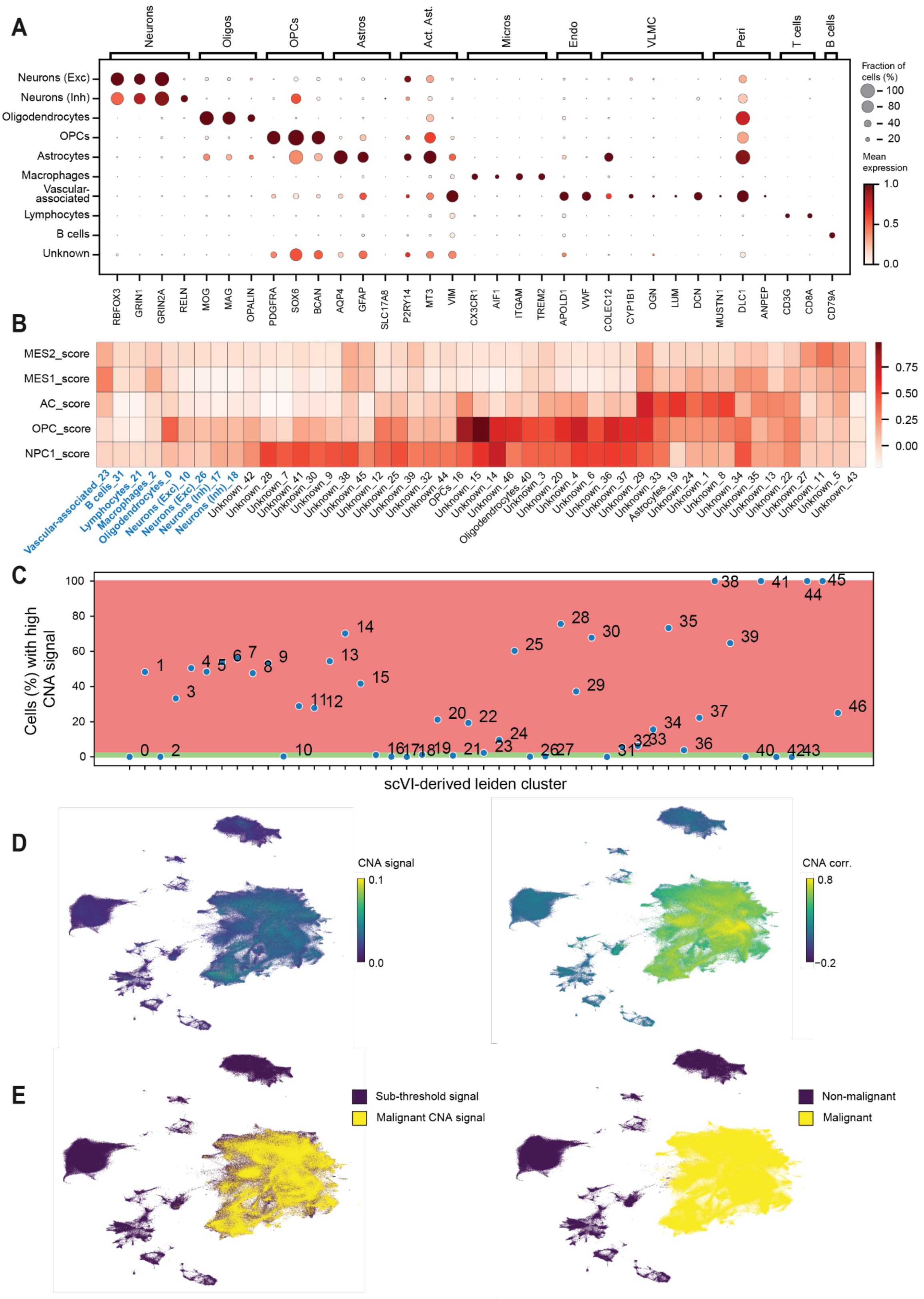


***Extended Data Fig. 3: Annotation and characterisation of malignant cell clusters.***
***A)*** Dotplot depicting marker gene expression among provisional leiden cluster annotations. Clusters are consolidated into low-resolution non-malignant cell classes (depicted along the y-axis). Colour corresponds to gene expression standardised per-gene, and dot size reflects the proportion of cells in a group expressing a given gene (expression > 0).

***B)*** Neftel et al. expression signature enrichment across all provisional clusters. Heatmap colours depict the median relative expression score for each Neftel gene set. Provisional clusters are listed along the x-axis with cluster annotations (A) listed as a prefix. Clusters annotated as non-malignant based on marker gene expression are coloured blue.

***C)*** Percentage of cells in each provisional cluster (annotated text) with an inferCNV-derived malignant CNA signal greater than minimum CNA thresholds (see Methods). Coloured patches indicate a malignant signal greater than or equal to 5% and green patches indicate a malignant signal percentage lower than 5%.

***D)*** UMAPs depicting metrics for malignant signal cell assignment: CNV signal (left) and CNA correlation (right) (see Methods).

***E)*** UMAPs demonstrating cells with malignant vs. non-malignant CNA signal assignment (left) and clusters with above-threshold (≥5%) malignant cell proportions (right).

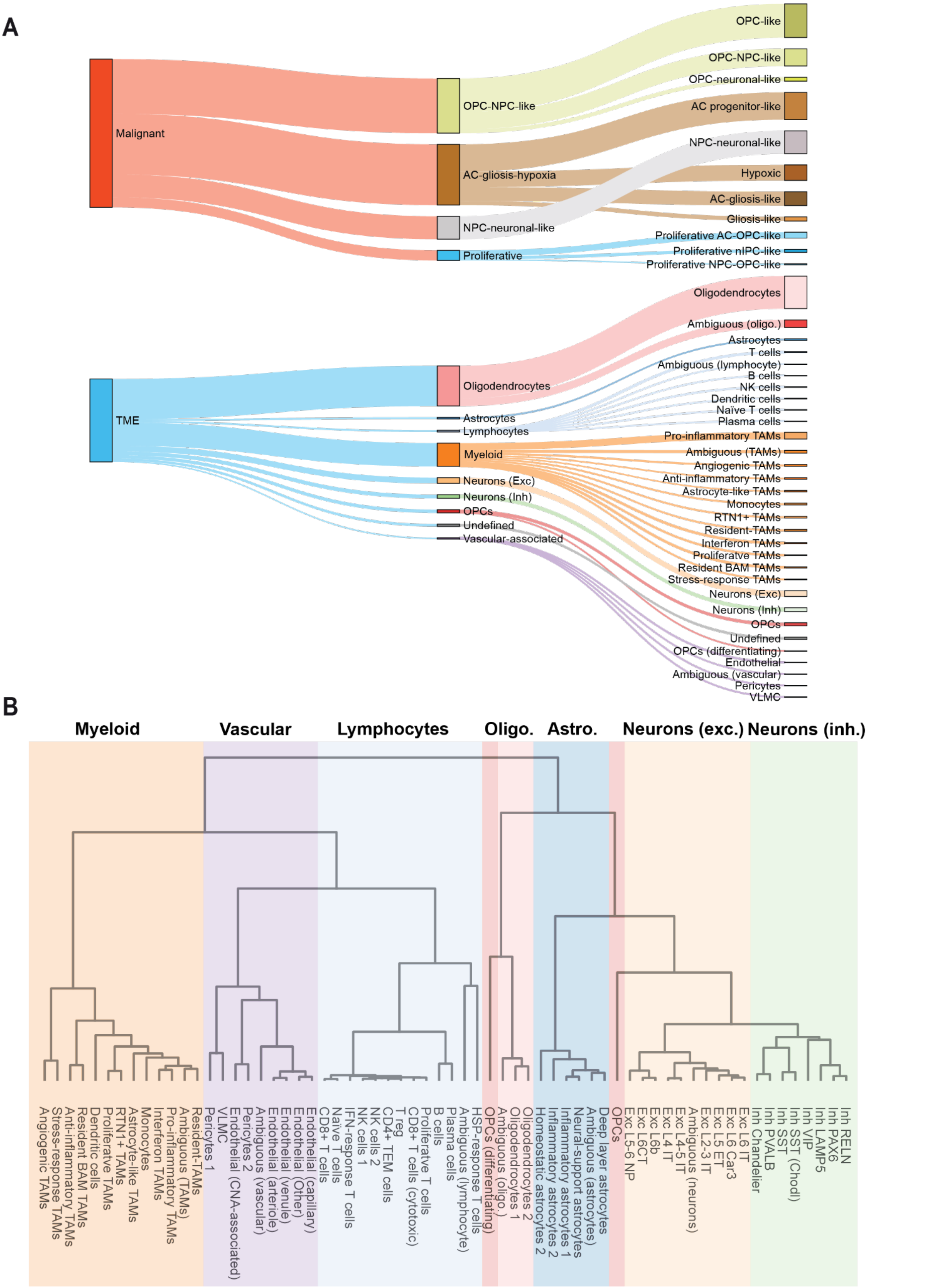


***Extended Data Fig. 4: Hierarchical annotation of malignant and TME states in the multiome data.***

***A)*** Sankey plot demonstrating frequently referenced annotation levels and their hierarchical relationship. This includes annotation levels referred to as malignant status (left) and cell class (middle), and coarse-grained (right, malignant-only) cell state annotations. High-resolution TME states belonging to key elements of the GB immune compartment are also referenced (i.e. lymphocytes, myeloid, and vascular-associated cells).

***B)*** Hierarchical clustering of all granular TME cell states identified in this study. Colours correspond to the coarse-grained annotation label associated with each cell state (see A).

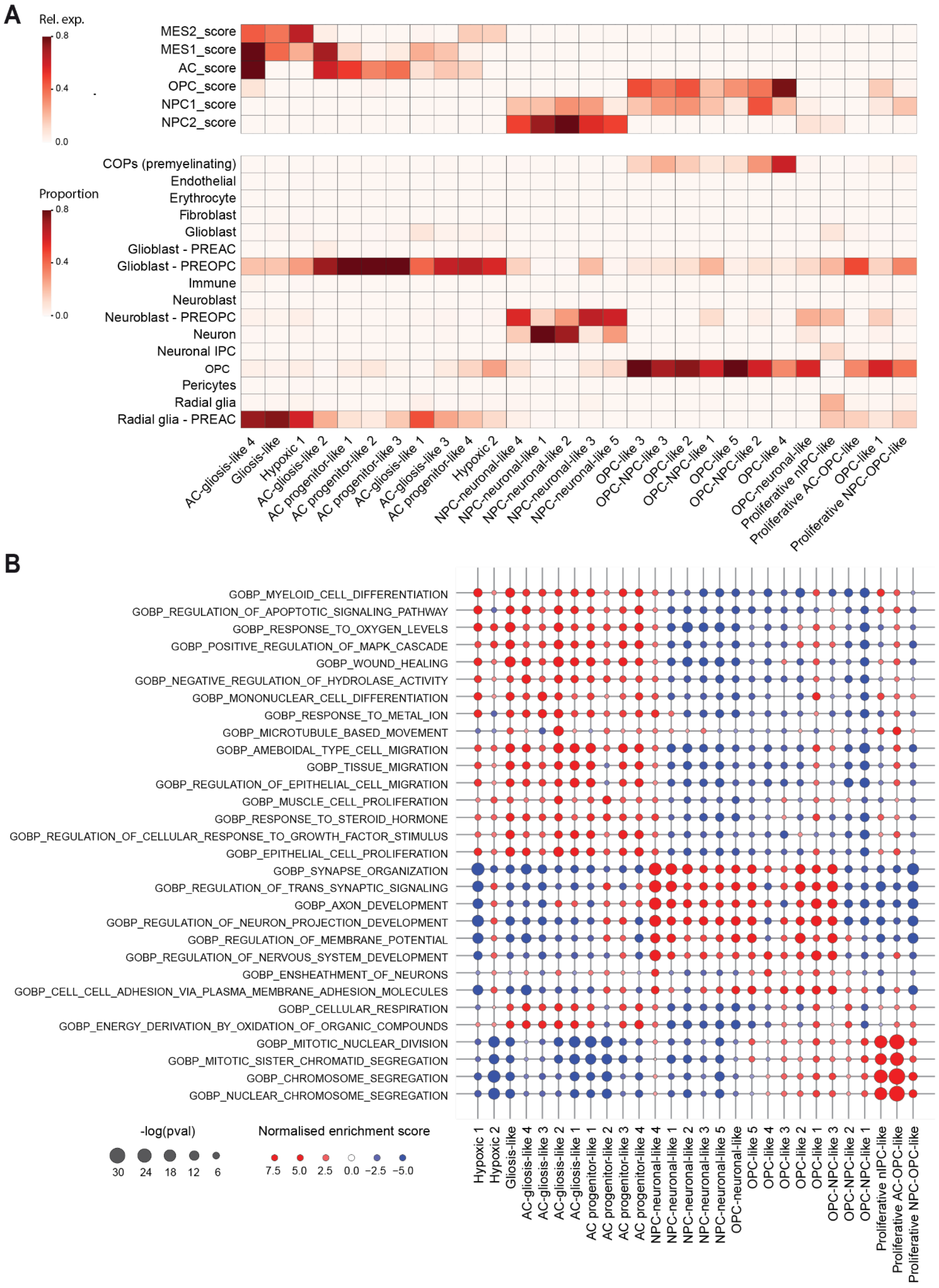


***Extended Data Fig. 5: Annotation of malignant cell states based on developing human brain label transfer and pathways analysis.***

***A)*** Comparison between granular malignant cell states based on relative expression of Neftel et al. expression signatures (top) and cluster-specific proportions of cell types labels transferred from Braun et al. using scPoli (bottom).

***B)*** Dotplot summarising the highest-scoring biological process GO terms identified through gene set enrichment analysis for each malignant cell state. Normalised enrichment score (within cluster) is indicated by colour, and significance of the observed enrichment score is reflected by dot size.

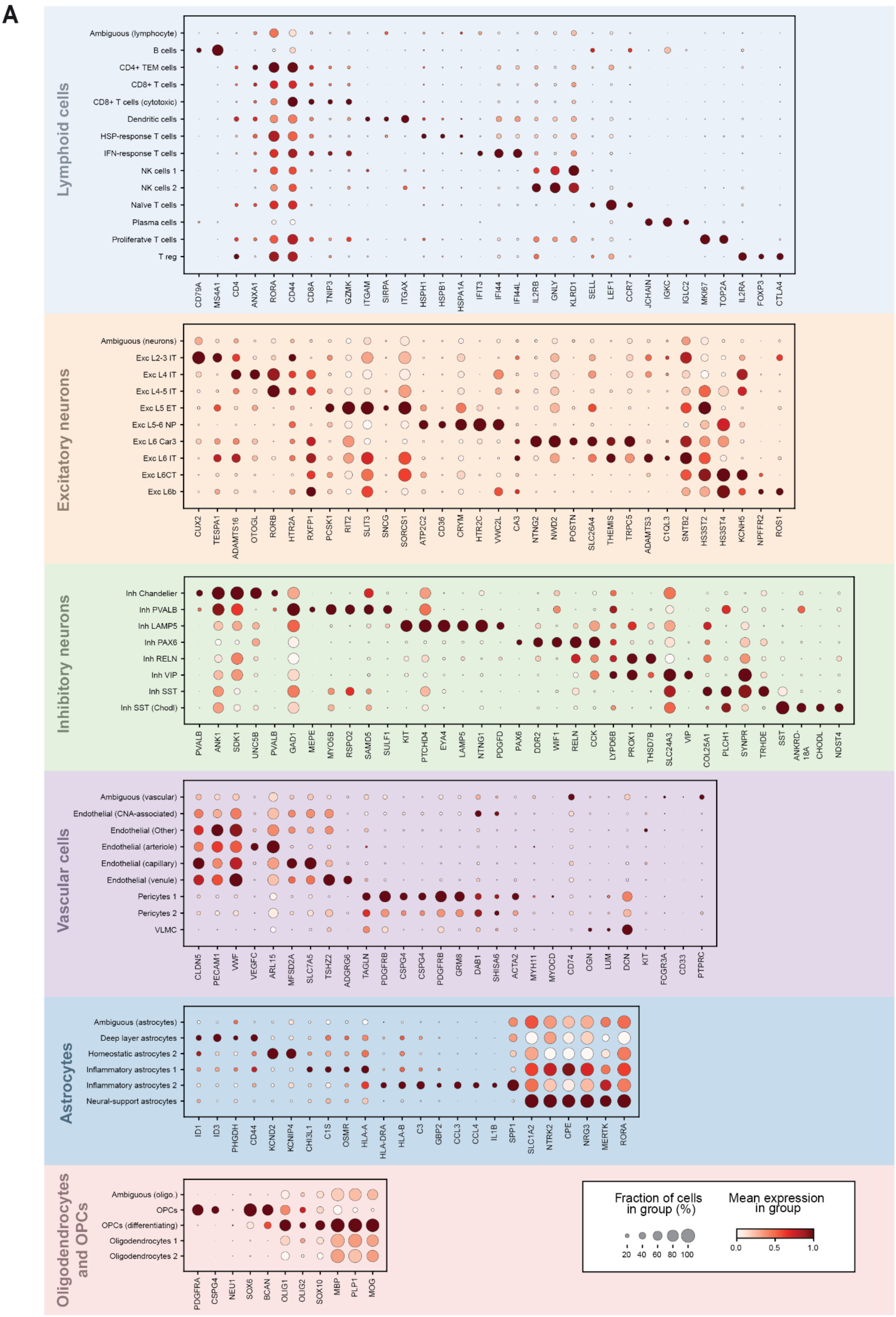


***Extended Data Fig. 6: Gene expression markers of high-granularity TME populations.***
***A)*** Dotplots summarising TME marker expression. Dotplots correspond to broadly defined populations as indicated by banner colour and title (left). Marker size reflects the fraction of

cells in a given group expressing a gene, whereas colour corresponds to mean expression within each group and is min-max normalised for each gene. Legend is shared across all plots (bottom right).

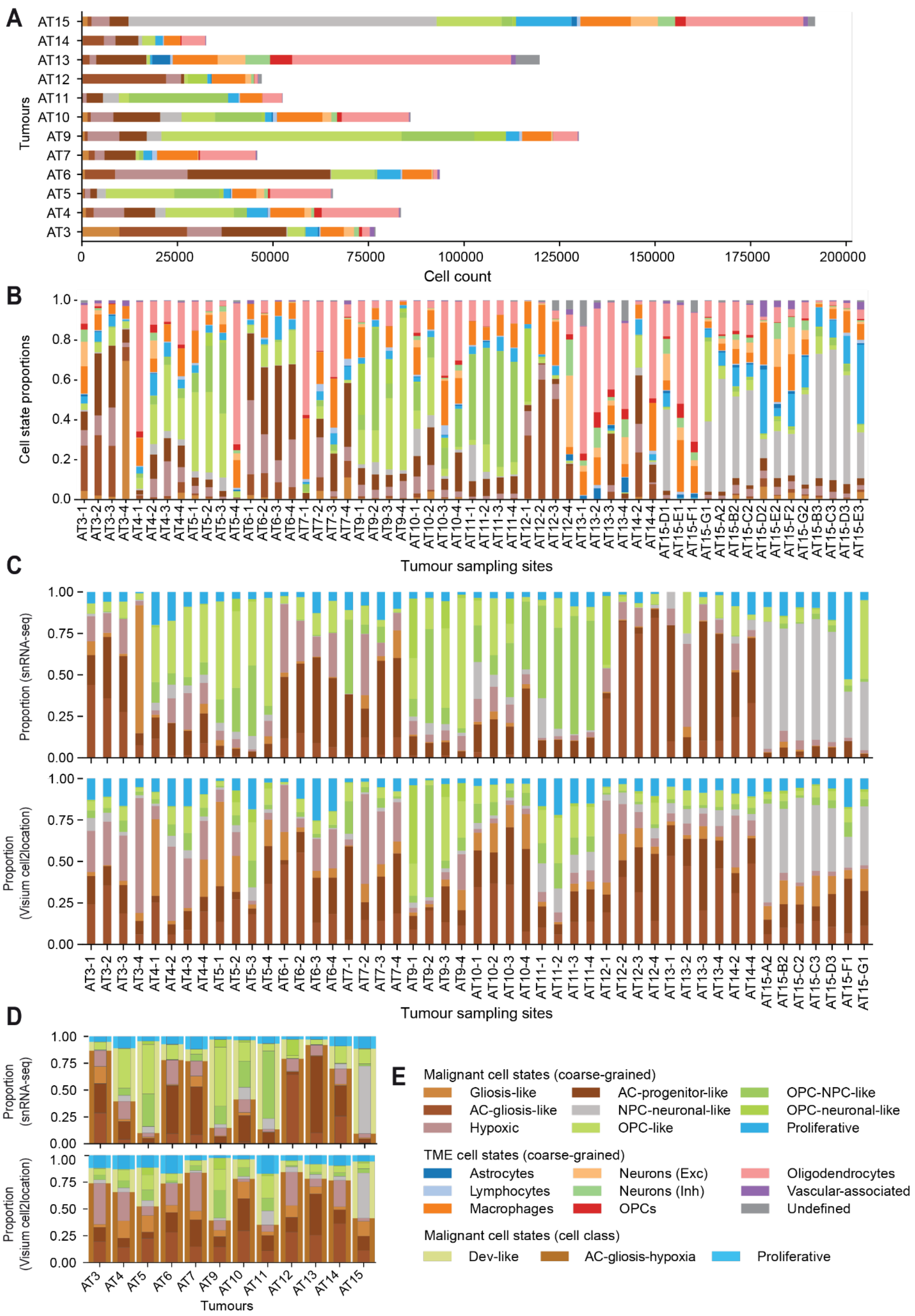


**Extended Data Fig. 7: Cross-modal malignant cell state distributions.**

***A)*** Counts of coarse-grained cell states for each tumour computed using snRNA-seq data. Colours correspond to individual coarse-grained malignant and TME cell states (E).

***B)*** Proportions of coarse-grained cell states for each tumour site computed using snRNA-seq data. Colours correspond to individual coarse-grained malignant and TME cell states (E).
***C)*** Modality comparison of cell state distributions across sampling sites. Proportion of coarse-grained malignant cell states for each tumour site based on both snRNA-seq data (top) and cell2location abundances from the Visium data (bottom). Colours correspond to coarse-grained malignant indicated below (E).
***D)*** Modality comparison of cell state distribution across each tumour. Barplots show the relative abundance of coarse-grained malignant cell states superimposed on broader malignant cell classes for snRNA-seq data (top) and cell2location abundance data (bottom), respectively..
***E)*** Legend mapping colours to cell state identities previously referenced (A-D).

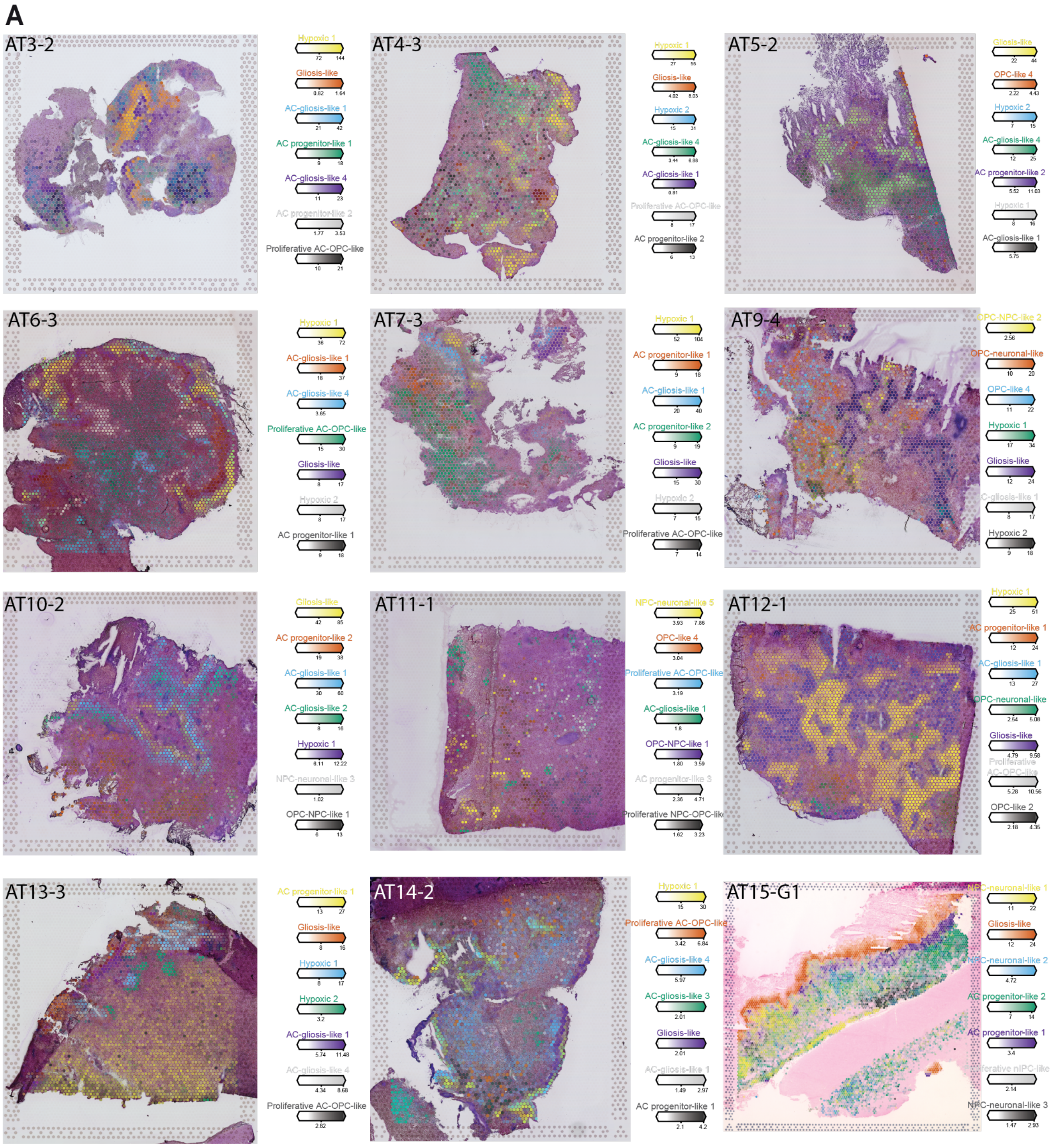


***Extended Data Fig. 8: Examples of malignant cell state spatial maps across all tumours.***

***A)*** Images depict overlayed cell2location-derived cell state abundances (colour intensity) for the highest-abundance cell states (colour) across each tumour. The top 7 cell states were selected on the basis of ranked maximum abundance. One section from a single site was included (top left).

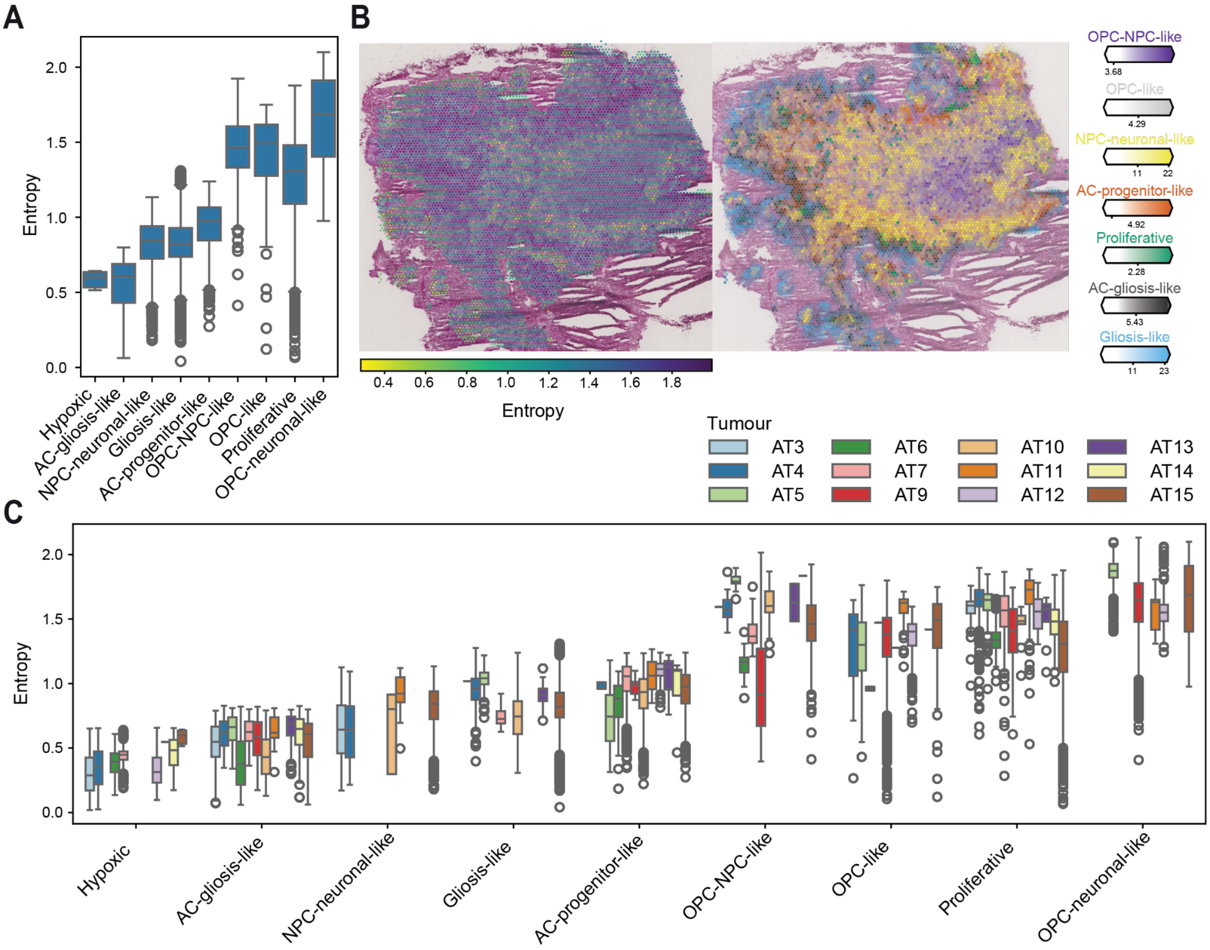


***Extended Data Fig. 9: Regionalised differences in spatial entropy consistent across tumours.***

***A)*** Boxplot summarising the Shannon entropy of spots with the highest abundance of a given cell state (abundance > 98$^{th}$ percentile) across tumour AT15.

***B)*** Visual comparison of entropy (left) and cell state abundance (right) in AT15 site C3. Low entropy (yellow) corresponds to spots with low cell state diversity, and high entropy (purple) reflects high cell state diversity spots. Cell state colour maps to individual cell states and the intensity is representative of relative abundance.

***C)*** Boxplot summarising the Shannon entropy of high-abundance (abundance > 98$^{th}$ percentile) spots for coarse-grained cell state across all tumours. Colour corresponds to individual tumour.

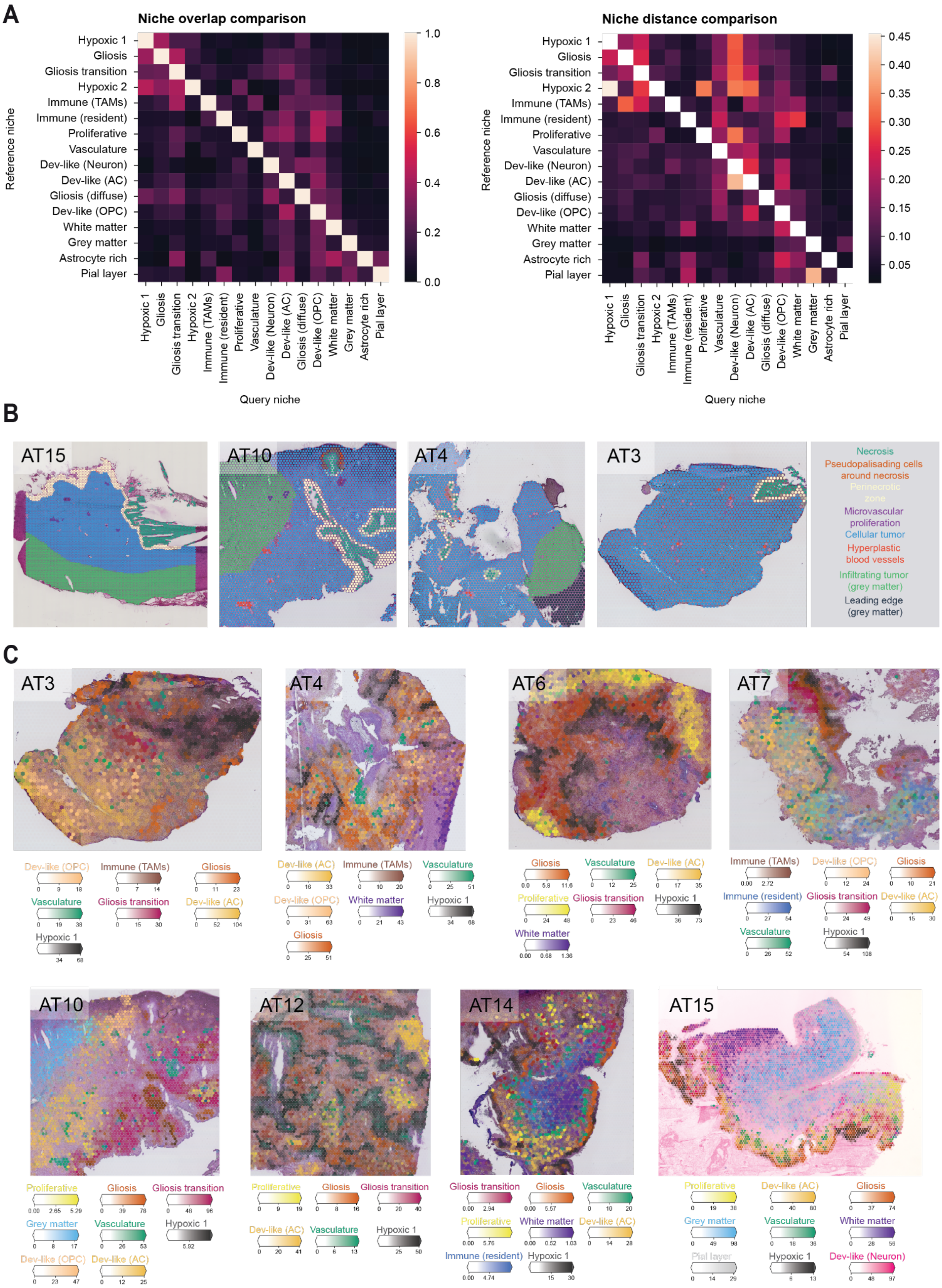


***Extended Data Fig. 10: Recurrent tissue niches of GB and their pathological annotation.*** ***A)*** Spatial relationship between each niche according to the extent of niche overlap (left) and euclidean distance (right). Niche overlap represents the fraction of spots annotated as a given niche (reference niche) that are also annotated as other niches (query niche). Niche distance is the average minimum euclidean distance between spots annotated as a given niche (reference niche) compared to other niches (query niche). Spots were assigned to a niche if the niche abundance was greater than the median or an absolute abundance threshold of 10.

***B)*** Histopathological annotations of Visium H&E images transferred to Visium voxels. Colour corresponds to histopathological features (right) and colour intensity reflects the degree of overlap, with partially overlapping features showing less intensity.

***C)*** Spatial niche (colour) abundance (colour intensity) overlaid on Visium H&Es for a subset of patients, demonstrating niche regionalisation. Niches with the highest max abundance visualised.

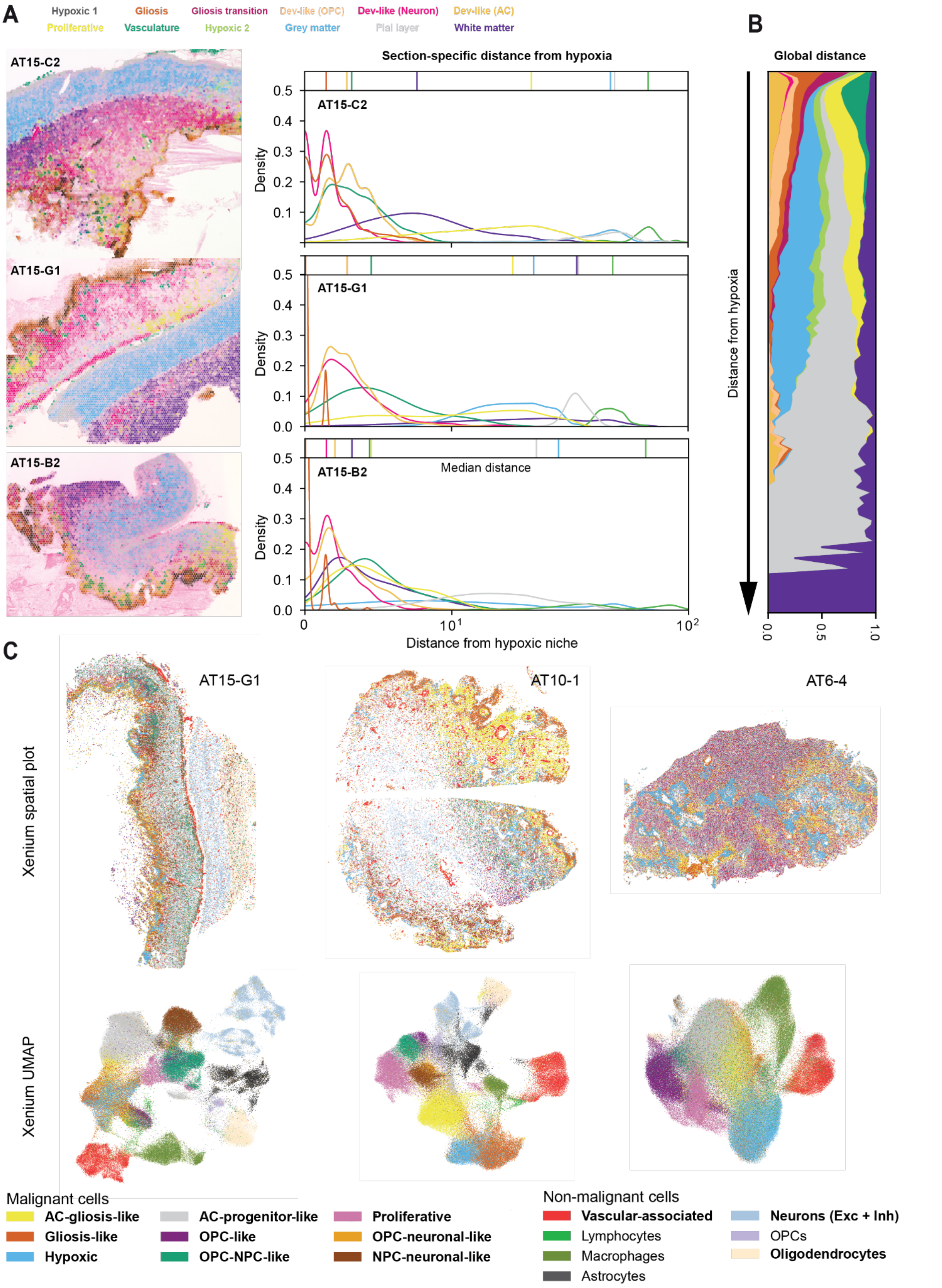


***Extended Data Fig. 11: Conserved spatial patterning of niches throughout a tumour and Xenium validation of malignant spatial organisation.***

***A)*** Spatial patterning of niches, from regions of hypoxia and necrosis to diffusely infiltrated normal brain cell types. Visium sections (left) include three sites originating from tumour AT15 showing overlaid niche abundances (colour). Density plots (right) illustrate the distribution of minimum euclidean distances from a spot of a given niche to the Hypoxic 1 niche, the latter representing necrosis. Colours of both plots map to niche identity presented in the legend (top).

***B)*** Streamgraph depicting the proportion of niches associated with a given distance from Hypoxic 1 across all tumours. Colours correspond to niche identities (A).

***C)*** Xenium sections (top) from three separate tumours coloured according to coarse-grained snRNA-seq annotations transferred using Tangram. UMAPs from Xenium cells depicting the topological relationship of transferred annotations. Colours map to the coarse-grained cell state listed in the legend below. To avoid overcrowding, Xenium spatial scatterplots were plotted using a subset of cell states, which are listed in bold font.

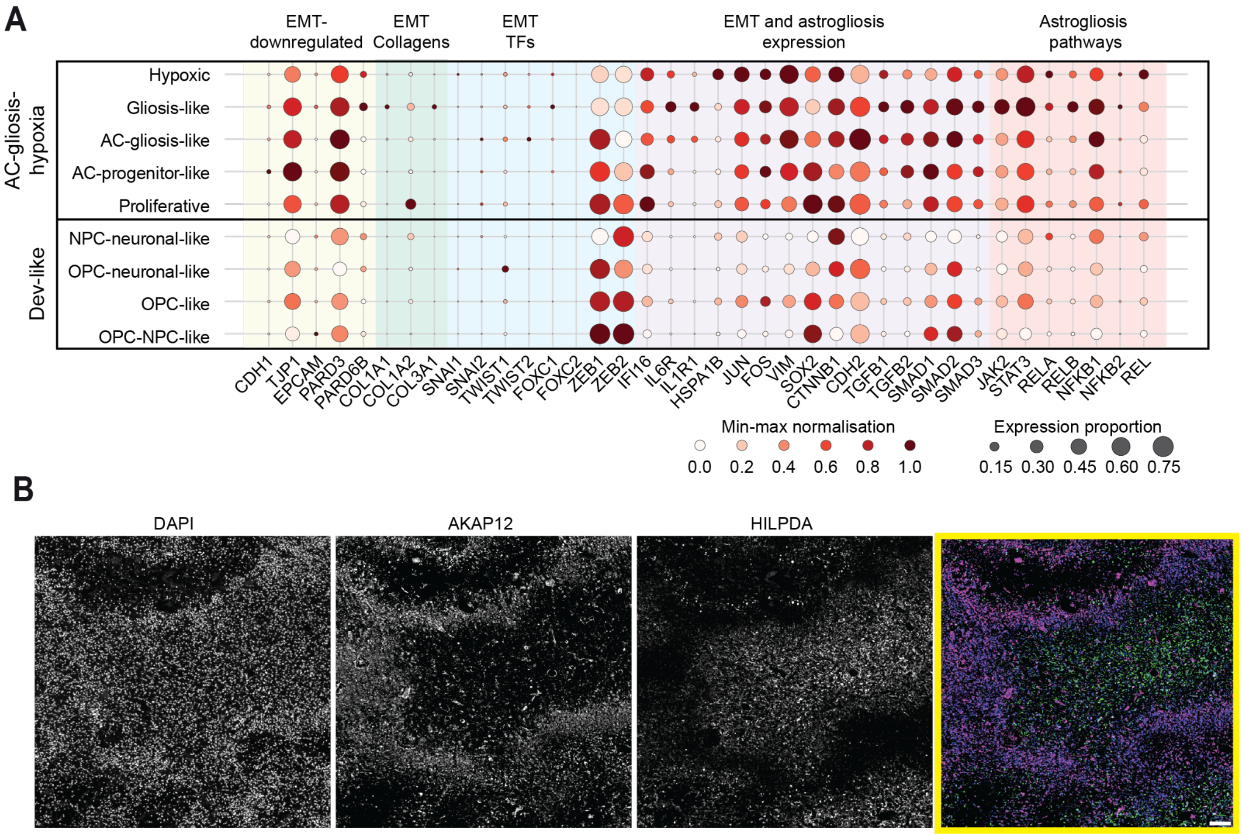


***Extended Data Fig. 12: Additional validation of spatial and functional partitioning of the AC-hypoxia anatomical axis.***

***A)*** Dotplot depicting common EMT markers compared to markers associated with astrogliosis. Colour intensity is the mean expression across a cell state with min-max normalisation applied across each gene. Dot size corresponds to the proportion of cells in a group expressing a given gene (expression greater than 0). Colours correspond to groups of EMT-associated genes and genes associated with both EMT and astrogliosis (including "Astrogliosis pathways").

***B)*** Immunofluorescent staining of GB tissue section revealing distinct patterns of AKAP12 (AC-gliosis) and HILPDA (hypoxic) protein expression. Panels show single channels of a region of interest (scale bar: 100µm).

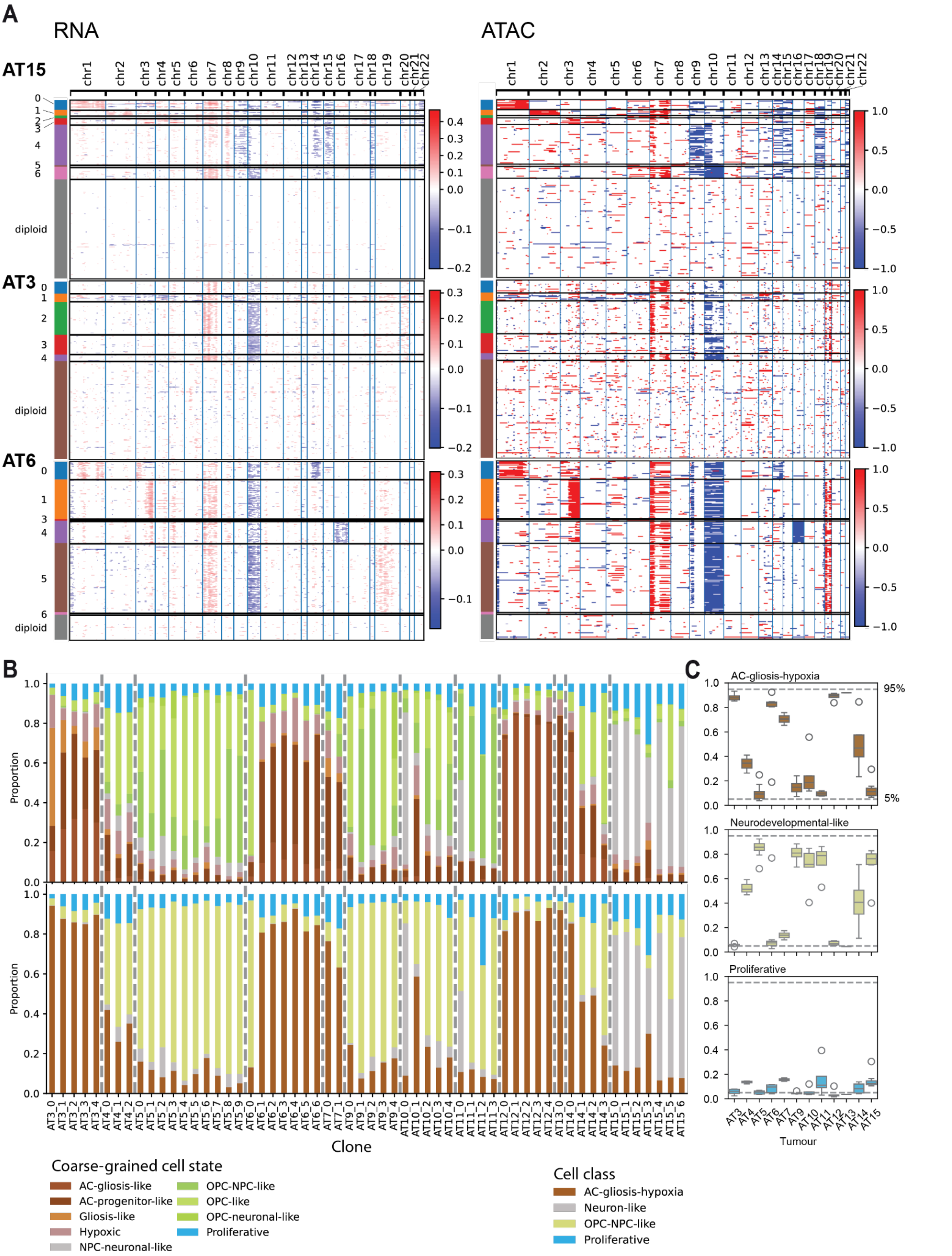


***Extended Data Fig. 13: Cross-modality CNA calling and subcone identification using RNA and ATAC multiome data.***

***A)*** Chromosomal CNAs inferred from both snRNA (left) and snATAC (right) data for three tumours. Rows map to individual cells and are clustered by putative subclones identified from the joint RNA-ATAC subclone calling (see Methods). Cell clusters representing subclones are indicated by the banner to the left of the plot.

***B)*** Proportions of cell states comprising each subclone for both coarse-grained cell states (top) and GB cell classes (bottom).

***C)*** Boxplots illustrating the spread of cell state proportions across all subclones in each tumour for each major cell class. The dashed lines highlight frequencies of 5% and 95%, representing a guideline indicating substantive frequencies of a given cell state.

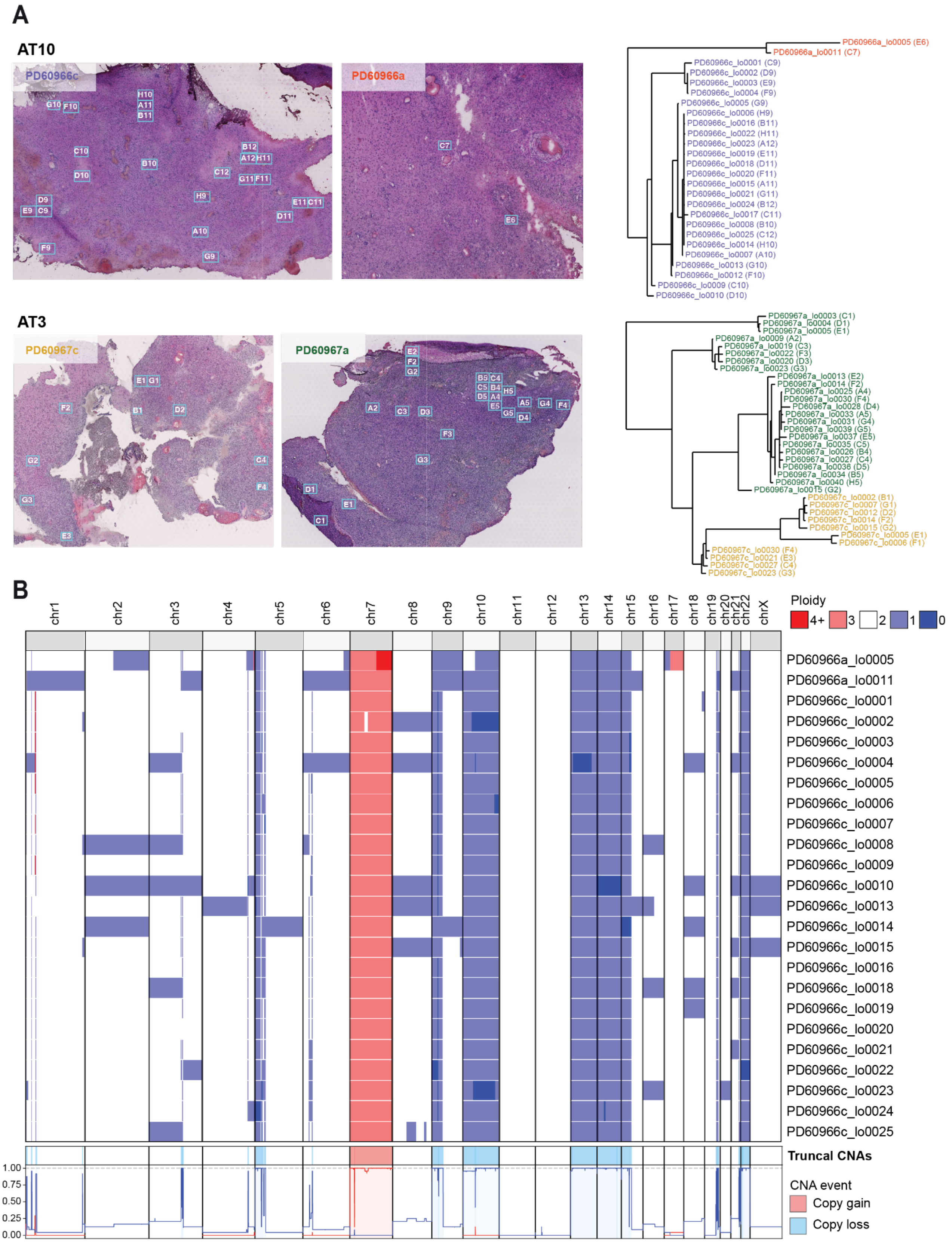


***Extended Data Fig. 14: Relationship between LCM cuts and identification of truncal mutations.***

***A)*** Spatial and genomic relationship between LCM ROIs. H&E images show the LCM ROIs superimposed on the adjacent Visium H&E section (top) for each tumour. Phylogenies (right) show genomic relatedness between ROIs and were constructed using somatic variants outside of regions with major CNs.

***B)*** Copy number results for ROIs from tumour AT10. The heatmap (top) shows Battenberg CN calls for each AT10 ROI. The line plot (bottom) summarises CN event concordance across samples and highlights highly concordant CNs suggestive of truncal CN events.

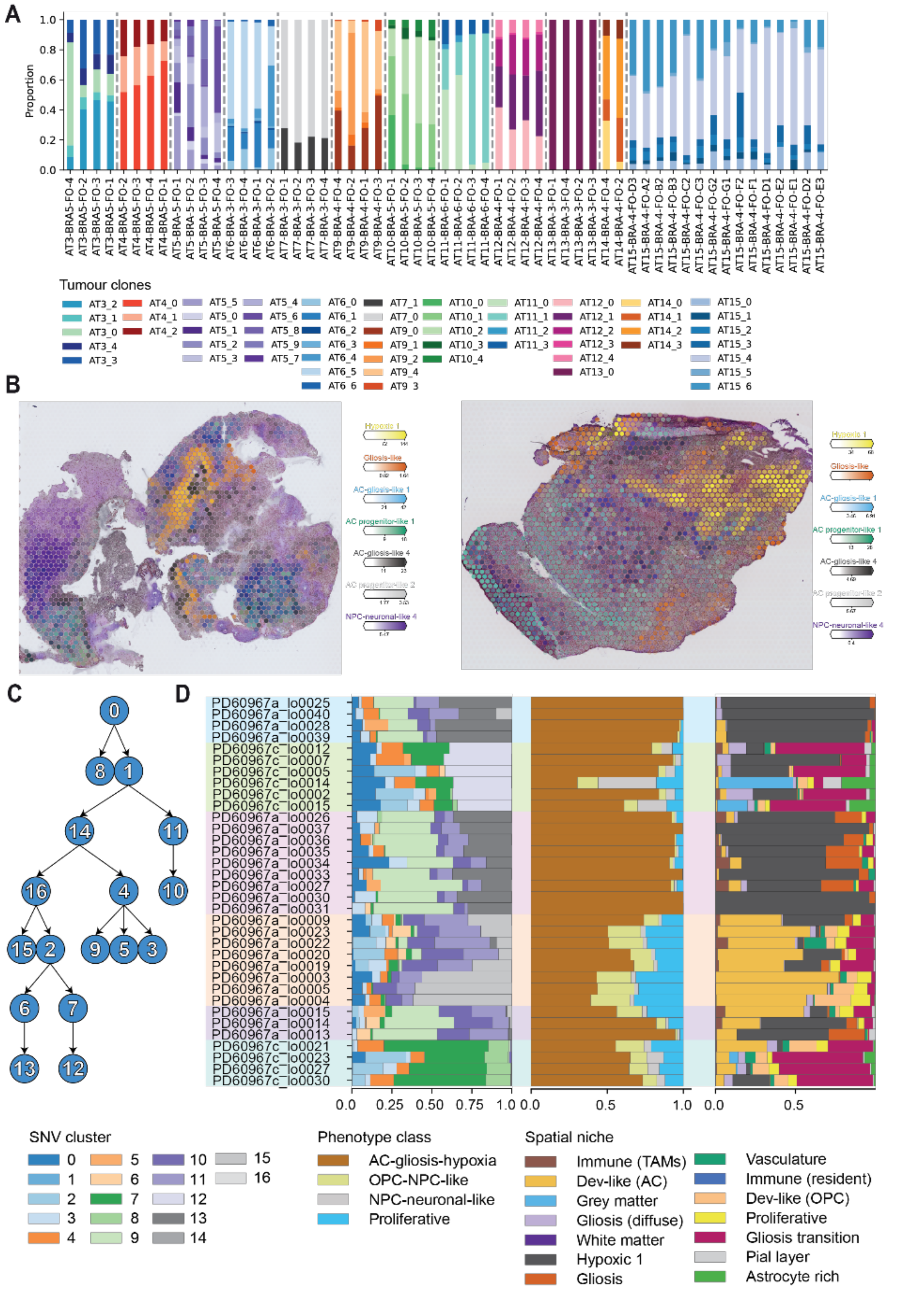


***Extended Data Fig. 15: Spatial context of tumour clones derived from transcriptomic and genomic data.***

***A)*** Proportion of SpaceTree-derived tumour clones within each tumour sampling site. Colours map to clone identity.

***B)*** Cell state (colour) abundance (colour intensity) inferred via cell2location for tumour AT3. Colours correspond to the top 7 high-abundance cell states across the tumour. Sections are histologically consistent with AT3 LCM ROIs (Figure S14).

***C)*** Phylogeny depicting possible evolutionary relationship between SNV clusters in tumour AT13. SNV clusters were identified with pyclone-VI and the phylogeny was inferred using PairTree.

***D)*** Relative frequency of SNV clusters per LCM sample (left) compared to the proportions of cell class (middle) and spatial niches (right). LCM samples are clustered and coloured based on composition clusters derived from SNV cluster distributions.

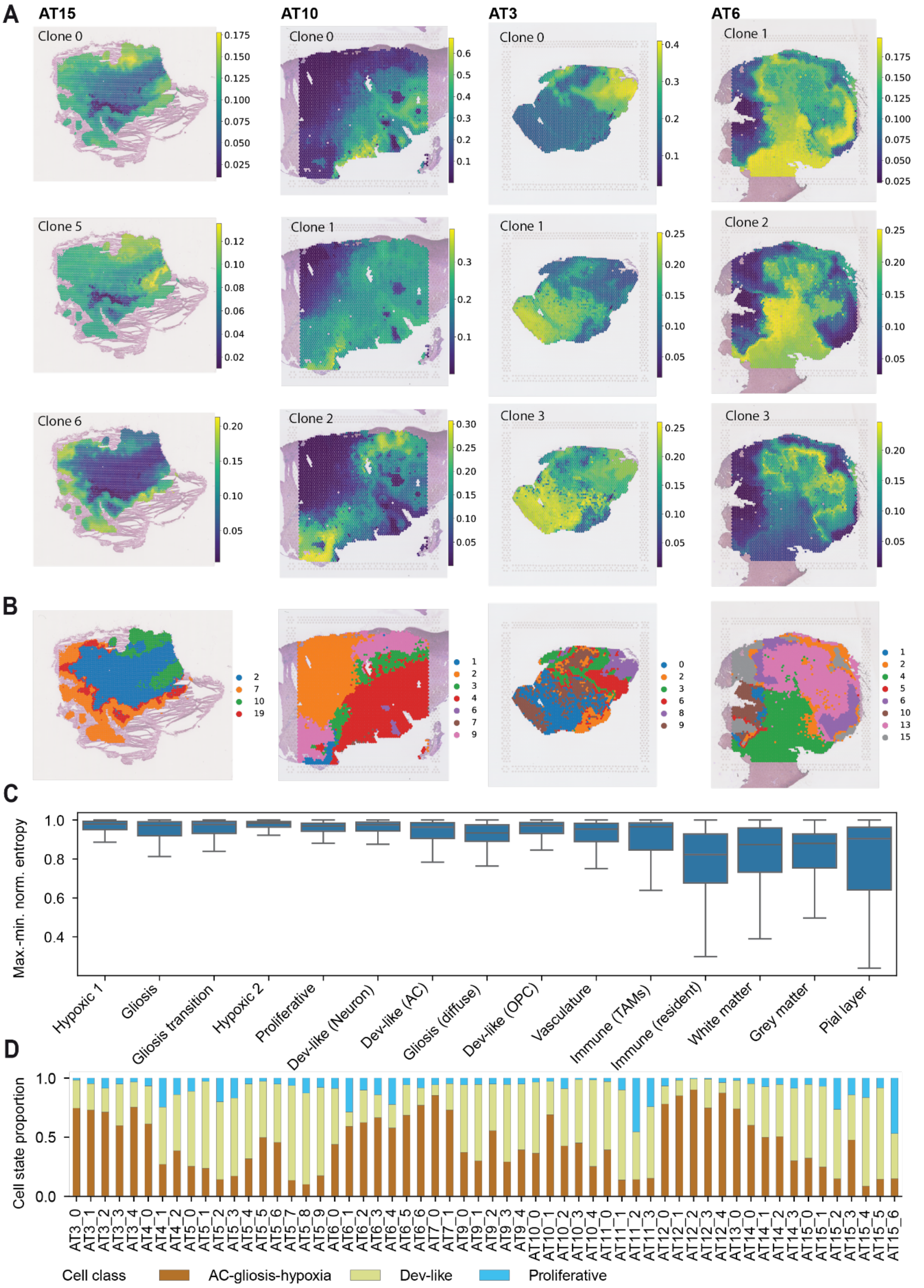


***Extended Data Fig. 16: Spatial mapping of clones with SpaceTree.***

***A)*** Spatial maps of SpaceTree clones for a subset of clones across four tumours. Colour intensity corresponds to the inferred frequency of each clone in a given Visium voxel.

***B)*** Compositional clusters derived from relative frequency of clones in each voxel. Colours map to composition cluster. Tumour identity is consistent with tumours listed above (A).

***C)*** Boxplot summarising clonal entropy across spatial niches for all patients. Entropy calculated for spots with high abundance of a given niche (abundance > $98^{th}$ percentile). Data points corresponding to outliers (fliers) omitted from plot for visual clarity.

***D)*** Cell class (colour) proportions for spots associated with each clone. Clone identities were defined according to the clone with the highest frequency in a given spot.

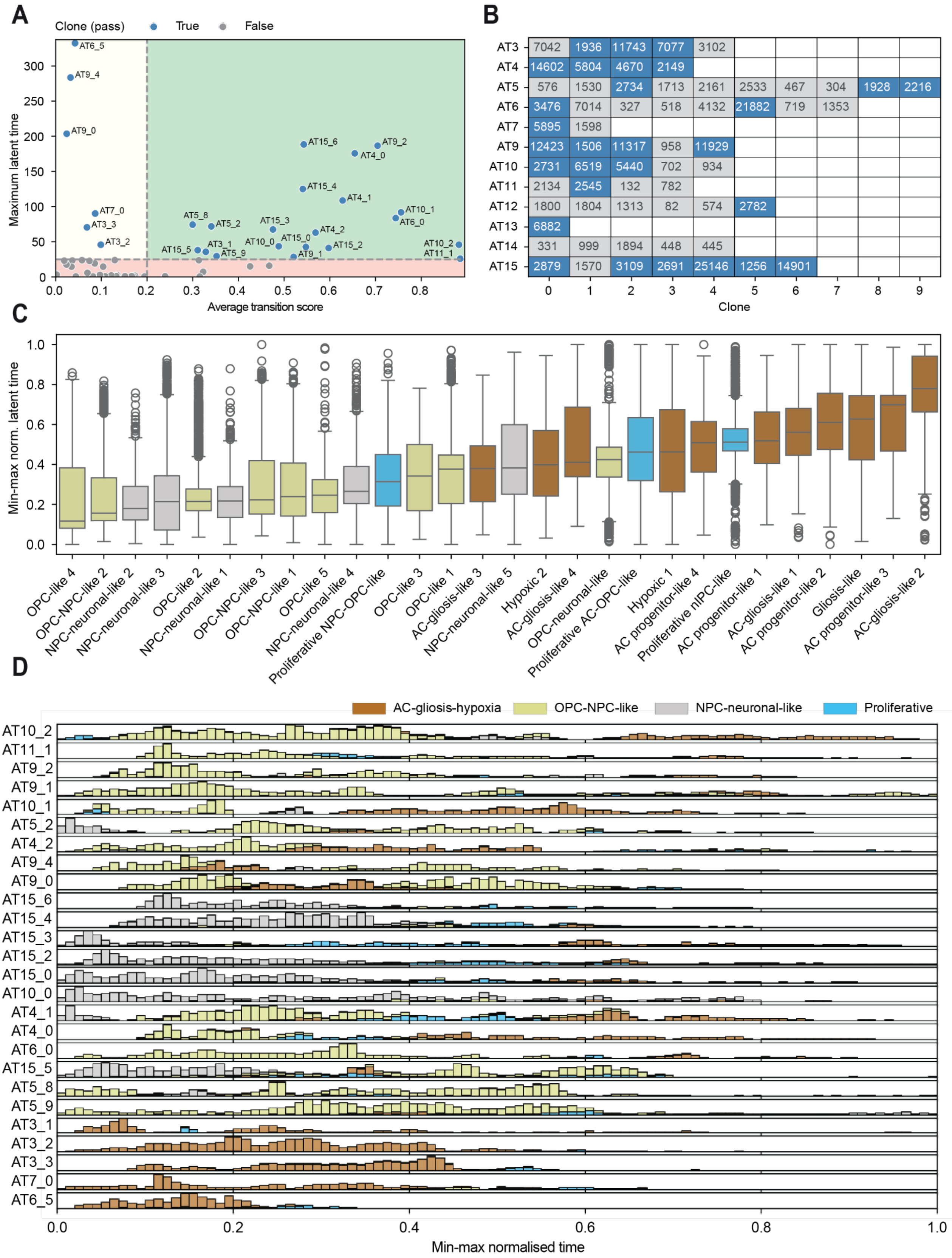


***Extended Data Fig. 17: Clonal trajectory inference with cell2fate.***

***A)*** Scatterplot depicting major QC metrics for cell2fate trajectory results. Patches correspond to QC thresholds: maximum latent time > 25 (red), and average transition score > 0.2 (yellow). Trajectories below the maximum latent time threshold (grey) were omitted from analysis, and those passing were used (blue) but flagged if they fell beneath the transition score threshold.

***B)*** Clones which passed trajectory QC thresholds (blue) compared to those which were omitted (grey) for each tumour.

***C)*** Boxplot summarising transition scores between granular cell states in among good clones (Figure S17A). Transitions (x-axis) are denoted by “>” and are ordered such that the first cell state transitions into the second. Colours (legend above) correspond to the cell classes involved in the transition. The red dashed line represents the threshold for transition scores used during QC.

***D)*** Boxplot summarising min-max normalised latent time for each cell state across all clones passing QC thresholds (A). Box colour corresponds to cell class. Legend shared with Figure S17E.

***E)*** Histograms showing cell class distribution across latent time for each clone passing QC thresholds. Colours correspond to cell class. The x-axis represents min-max normalised latent time.

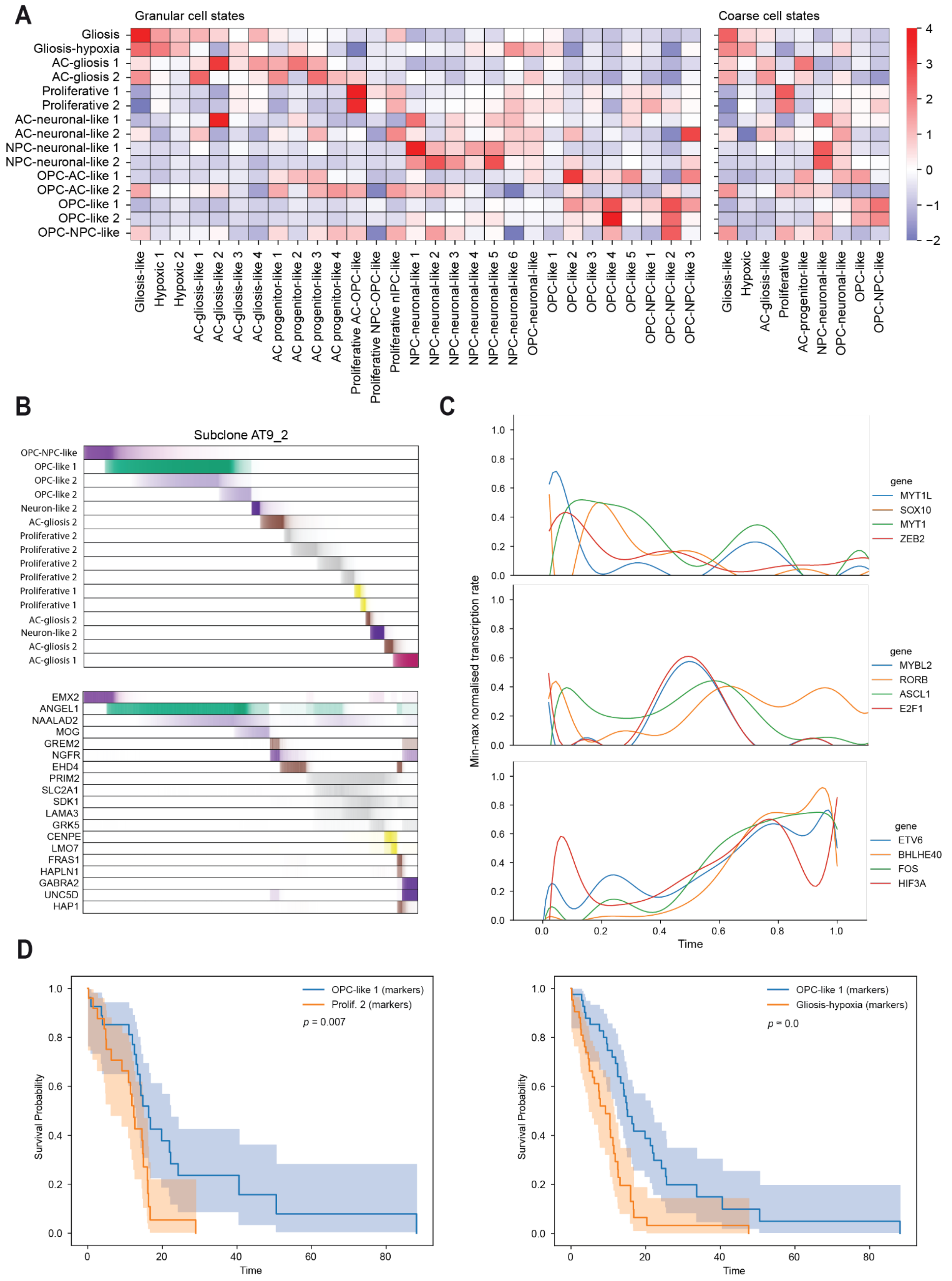


***Extended Data Fig. 18: Inference of meta-modules derived from clonal cell2fate trajectories.***

***A)*** cell2fate meta-module enrichment among granular cell states (left) and coarse-grained states (right). Colour corresponds to z-score normalised mean relative expression scores for each cell state.

***B)*** Annotated meta-module activation across inferred time (top) and top markers associated with each meta-module (bottom) for clone AT9_2. Colours correspond to recurrent meta-module and colour intensity maps to the activation pattern.

***C)*** Example of TF expression rate over time for TFs implicated in three different stages of inferred latent time across three clones (AT9_2, AT15_3, and AT4_1): early time (top), mid-range time (middle), and late time (bottom) TFs. Line plots are summarising TF rate across all stated clones. Colours correspond to TF identity.

***D)*** Kaplan-Meier plots comparing early time OPC-like 1 meta-module compared against markers of mid-range Proliferative 2 (right) and late time Gliosis-hypoxia. CPTAC and TCGA samples were categorised based on higher relative expression scores of one set of meta-module markers over another. The *p*-value reflects the results of a log-rank test comparing samples associated with each meta-module.

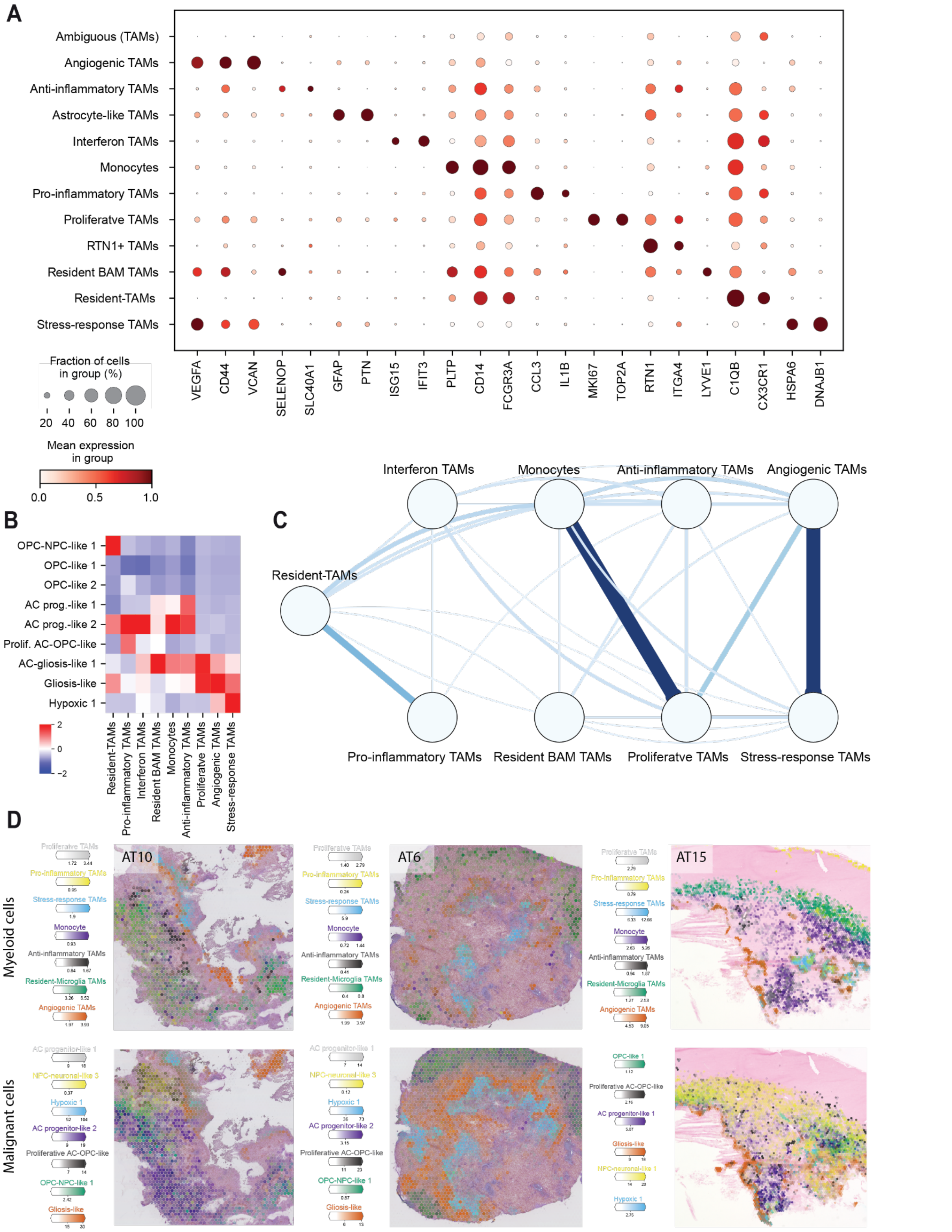


***Extended Data Fig. 19: Myeloid cells form distinct spatial immune compartments across GB tumours.***

***A)*** Dotplot highlighting myeloid cell state expression markers involved in cell state annotation. Colour represents the mean expression of markers with min-max normalised applied per gene. Dot size reflects the proportion of cells expressing a certain gene.

***B)*** Proximity of common malignant cell states to myeloid populations. Mean euclidean distance between a given myeloid-associated Visium spot and its nearest neighbour for each malignant cell state. Distances are z-score normalised within each myeloid population.

***C)*** Spatial proximity between myeloid populations. Edge width and colour represent the mean minimum pairwise euclidean distance between spots associated with a given population. Mean values calculated based on the closest 25% of spots to account for differences in spatial dispersion.

***D)*** cell2location abundances across sections from three different tumours summarising the spatial distribution of most common myeloid cell states (top) and malignant cell states (bottom). Colours map to cell state identity and intensity reflects the abundance.

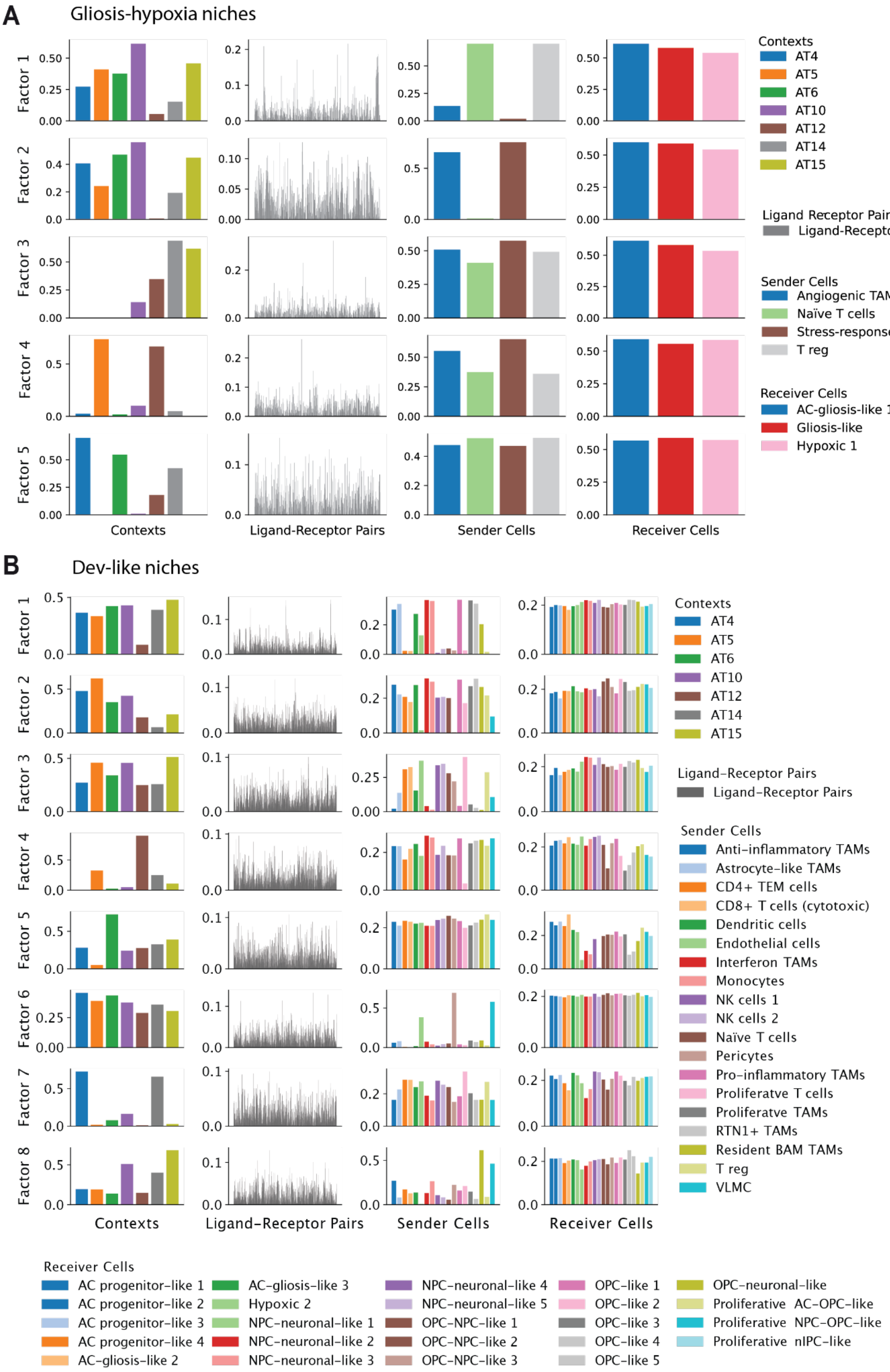


***Extended Data Fig. 20: Cell-cell communication analysis of within major tumour niches using Tensor-cell2cell.***

***A)*** Cell-cell communication analysis for hypoxic and peri-nectoric niches. Factors generated from the decomposition of a 4D communication tensor derived from a single-nucleus dataset of GB donors using the selected sender and receiver cell states. Each row represents a factor, while each column corresponds to the loadings for a specific tensor dimension. Five factors were used for this analysis where the context corresponds to donors. The columns represent donors, ligand-receptor pairs, sender cell state (TME cell states) loadings, and receiver cell state (malignant cell states) loadings, respectively. Bars are color-coded according to the categories assigned to elements in each tensor dimension as shown in the legend.

***B)*** Cell-cell communication analysis for the cellular tumour niches. Similarly, factors were obtained from the decomposition of the communication tensor created based on selected sender and receiver cell types. Eight factors were selected for this analysis. Sender and receiver cell states are available in the legend.

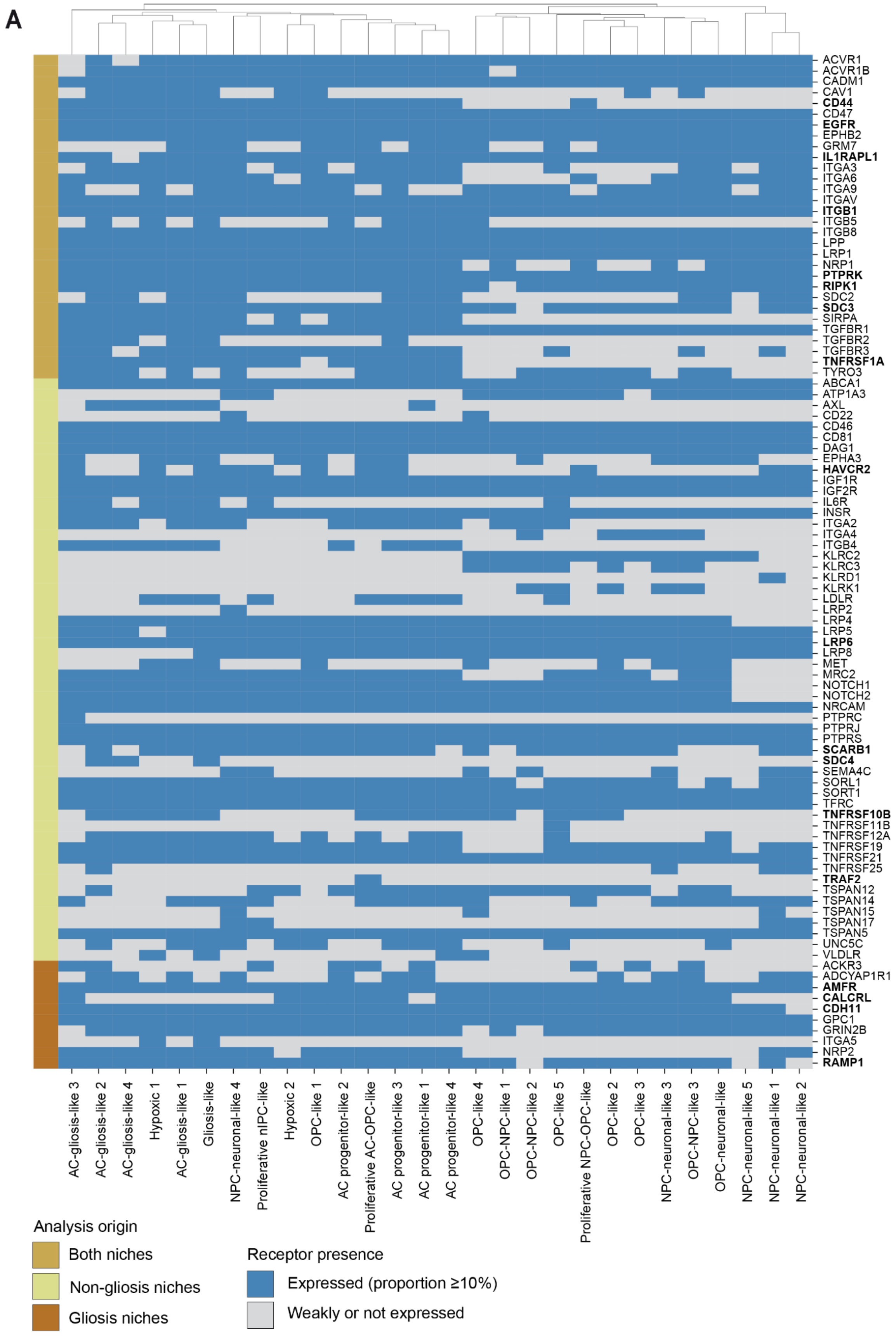


***Extended Data Fig. 21: Receptor expression in malignant cell populations of each CCC analysis.***

***A)*** The presence of a receptor (expressed in ≥ 10% of cells) across malignant cell states for CCC analysis of perinecrotic niches (gliosis) and cellular tumour (non-gliosis) niches. Analysis origin is highlighted in the coloured banner (left). Bolded receptors were listed in Figure 7E.
