## Supplementary Computational Note for "A spatiotemporal cancer cell trajectory underlies glioblastoma heterogeneity"

### spaceTree: Deciphering Tumor Microenvironments by joint modeling of cell states and genotype-phenotype relationships in spatial omics data

#### 1. Introduction

Spatial omics technologies have revolutionized the study of tumor evolution by providing spatially resolved molecular data, enabling researchers to investigate the intricate relationships between cellular composition and spatial architecture within tumors [1]. These technologies offer unprecedented opportunities to explore tumor heterogeneity, which manifests not only at the genetic and cellular levels but also across distinct spatial niches that govern cell type distribution and clonal composition within the tumor microenvironment.

While many existing methods have provided valuable insights into cell type composition and spatial mapping, accurately deciphering the clonal structure of tumors remains a complex challenge. Genetic subclones are closely intertwined with cellular phenotypes and are influenced by the spatial organization of the tumor, making it difficult to fully capture these relationships. Current approaches, though effective in certain domains, often focus primarily on cell type decomposition or mapping single-cell RNA sequencing (scRNA-seq) data onto spatial platforms. However, these approaches may not always be optimized for the joint modeling of both cell types and clonal profiles, potentially limiting the depth of integration between genetic and cellular information [2]. As a result, there remains a need for methods that can comprehensively capture both cellular and clonal states while accounting for their spatial interdependencies.

To address this gap, we propose spaceTree, an end-to-end framework built on a multi-task graph neural network architecture coupled with label propagation. spaceTree is designed to jointly model spatially smooth cell type and clonal state compositions, offering a more integrated perspective on tumor heterogeneity. By employing Graph Attention mechanisms[3], it selectively incorporates information from spatially adjacent regions, enhancing both interpretability and quantitative accuracy, particularly in cases where reference mappings may be incomplete or insufficient.

A significant strength of spaceTree is its technology-agnostic design, making it adaptable for clone mapping across both sequencing-based and imaging-based assays at a variety of resolutions. The model outputs can be used to characterize spatial niches with consistent cell type and clone compositions, offering a more holistic view of tumor architecture and providing deeper insights into the complex spatial dynamics of tumor heterogeneity.

#### 1.1 Background

Projecting genomic features onto spatial data has been a challenge within the spatial transcriptomic (ST) community. Several experimental protocols, such as BaSISS[4] and slide-DNA-seq[5], attempt to provide direct DNA readouts in spatially resolved contexts. However, these technologies remain niche and are not as widely democratized or accessible as other spatial platforms like 10x Genomics. Due to these limitations, solutions for integrating genomic information with spatial data primarily lie in the computational domain.

When direct DNA readouts are not available—particularly since scDNA-seq is also not widely adopted—a common strategy is to infer genomic features through copy number variation (CNV) inference tools applied to scRNA-seq data. Methods such as InferCNV[6], as well as other approaches for haplotype phasing and clonal subpopulation delineation[7, 8, 9], are widely used in the community. To bridge the gap between spatial transcriptomics and clonal inference, recent studies have adapted these RNA-based CNV inference methods for spatial data analysis[10, 11]. While these adaptations offer potential, they also present challenges. For spot-based ST technologies, interpreting results can be complicated due to the absence of a deconvolution step. Meanwhile, in cellular or subcellular resolution technologies, the sparse nature of the data can introduce ambiguity, complicating the identification of clones.

Several methods have been developed specifically for spatial transcriptomics data (Table 1), focusing on different spatial technologies. Many of these methods were designed for spot-based technologies, such as Clonalscope[12], Tumoroscope[13], CVAM[14], CalicoST[15], SlideCNA[16], and STARCH[17]. However, their applicability to other platforms is often not demonstrated or remains unclear. Among these, Tumoroscope[13] stands out by addressing the challenge of deconvolving the proportions of clones in spatial transcriptomics spots. This method requires bulk DNA-seq data with estimated clone genotypes, providing a novel approach to uncovering clonal architecture.

Although the above-mentioned methods do not require single-cell RNA-seq data, spatial transcriptomics studies frequently incorporate matched scRNA-seq data to enhance analysis resolution. For spot-based technologies, such data is crucial for performing spot deconvolution, as spots typically contain mixtures of multiple cells. Consequently, there has been significant focus on developing methods for cellular deconvolution in spatial transcriptomics. These methods aim to disaggregate the molecular signatures of mixed

| Method | Reference | Spatial information | Deconvolution | ST platforms |
|---|---|---|---|---|
| Clonalscope[12] | WGS/WES | No | No | Visium(demonstrated, but possibly not limited) |
| Tumoroscope[13] | WGS/WES | H&E images used | Yes | Any spot-based ST |
| SlideCNA[16] | No | Spatial locations | No | Slide-seq-like ST data |
| CVAM[14] | No | Spatial locations | No | Spot-based (demonstrated, but possibly not limited) |
| CalicoST[15] | No | No | No | Visium (demonstrated, but possibly not limited) |
| STARCH[17] | No | Yes | No | Visium (demonstrated, but possibly not limited) |

Table 1: Overview of methods for genomic feature mapping in spatial transcriptomics

cell types within low-resolution spatial spots.

In a comprehensive benchmarking study, Gao et al.[18] evaluated 18 deconvolution methods and identified Cell2location[19], CARD[20], and Tangram[21] as the top-performing approaches. Notably, each of these methods approaches the problem in a distinct way. For instance, unlike Cell2location and CARD, Tangram does not directly perform deconvolution but instead aligns sc/snRNA-seq data to spatial data. CARD stands out by incorporating spatial proximity information to improve accuracy. Interestingly, the benchmark shows that no single method consistently outperforms the others across all datasets, highlighting the context-dependent nature of their effectiveness.

Drawing inspiration from these deconvolution approaches, spaceTree expands their use to scenarios that require the simultaneous deconvolution of both cell types and clonal compositions. spaceTree relies on established scRNA-seq CNV inference methods to assign clone labels to cells and then projects both cell type and clone labels onto spatial data, enabling a comprehensive understanding of the spatial organization of cell types and clonal populations within a tissue.

### 2. Methods

spaceTree is a multi-step approach that performs a graph construction first and then employs multi-task GNN to predict labels for all spatial loci. Please see **Figure** 1 for a detailed illustration of the process.

#### 2.1 Input data

The initial step of the algorithm is the construction of the graph $G$ based on the following components:

- **Reference Count Matrix ($M_R$):** Transcriptome counts for reference nodes.
- **Reference labels ($L_R$):** Labels for reference nodes. In principle, any labels can be used, but here we assume they consist of clone IDs and cell types.
- **Spatial Count Matrix ($M_S$):** This matrix represents the transcriptome counts of spatial nodes.

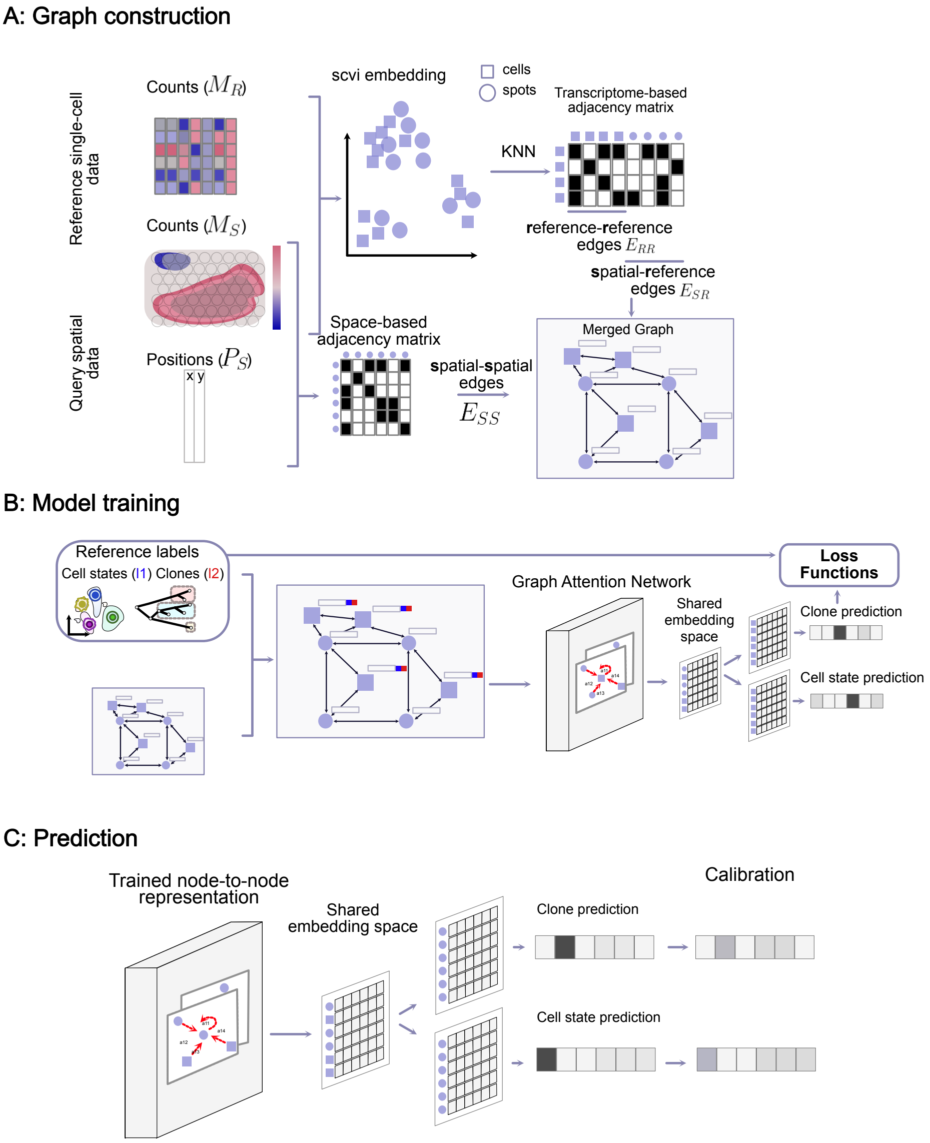


Figure 1: The complete workflow of spaceTree method. A: Construction of a merged graph that combines features from spatial and reference data in a single structure. B: Training of spaceTree model. The graph is used in a graph attention network that learns to predict node labels based on a combination of spatial and reference node features. C: Predictions are performed on spatial nodes based on the obtained node embedding and later calibrated to account for necessary mixing effects relevant to individual technologies.

- **Spatial Position Matrix ($P_S$):** This matrix specifies the physical positions of spatial nodes within a tissue, with each row corresponding to a spatial node in $V_S$. A row in $P_S$ contains the $(x, y)$ (or $(x, y, z)$) coordinates that define the node's position in the spatial domain.

The modalities mentioned above are the only requirements that are needed to execute spaceTree. All other steps mentioned below are a part of the spaceTree package and are demonstrated in our tutorials.

#### 2.2 Graph construction

spaceTree is a GNN-based framework that requires an input graph $G = (V, E)$, where $V$ represents the set of vertices (nodes) and $E$ represents the set of edges. The vertices are categorized into two distinct types: spatial nodes, denoted as $V_S$, and reference nodes, denoted as $V_R$. Thus, the entire set of vertices can be expressed as $V = V_S \cup V_R$, with $V_S \cap V_R = \emptyset$ indicating that the sets of spatial and reference nodes are mutually exclusive. Edges in the graph are similarly categorized based on the node types they connect, with $E_{SS}$ representing edges between spatial nodes, $E_{SR}$ representing edges between spatial and reference nodes, and $E_{RR}$ representing edges between reference nodes. Consequently, the set of all edges is $E = E_{SS} \cup E_{SR} \cup E_{RR}$. The algorithm operates in two steps: initial construction of $G = (V, E)$, and GAT-based label propagation.

To construct $G$ out of $M_S$, $M_R$ and $P_S$, we need to define procedures to obtain $E_{SS}$,$E_{RR}$ and $E_{SR}$:

- **Spatial-spatial edges ($E_{SS}$):** Spatial-spatial edges are computed based solely on physical distance between $V_S$. For greed-based technologies (such as Visium/Visium HD) the entire grid can be used as a source of $V_S$. Alternatively, a KNN-graph is computed based on $P_S$ using `sklearn NearestNeighbors` method. The distance in space is scaled between 0 and 1, where 1 indicates the shortest distance and then used as an edge feature.

- **Reference-Reference edges ($E_{RR}$):** Reference-reference edges are computed based on transcriptome similarity of nodes $V_R$ in $M_R$. These edges are also computed based on `sklearn NearestNeighbors` method. The distance in transcriptome is scaled between 0 and 1, where 1 indicates the largest similarity and then used as an edge feature.

- **Spatial-Reference edges ($E_{SR}$):** Construction of Spatial-Reference edges is the most challenging step of the initial preprocessing due to possible distribution shifts between $M_R$ and $M_S$. These distribution shifts do not always allow to rely on procedures such as Nearest Neighbours directly. While it is not necessary to have a perfect initial edge definition, good preliminary results can be obtained based on methods such as `scvi`. `scvi` allows to train a Conditional Variational Autoencoder that can address distribution shifts. The optimal choice of the method depends on the user's preferences but in our experiments `scvi`-based embeddings served as a decent preliminary source of $E_{SR}$ even in datasets with compositionally (see more details below). To obtain $E_{SR}$ from the embedding layer, we used the same KNN-based strategy as described above. The distance in transcriptome is then scaled between 0 and 1 and used as an edge feature.

Each of the nodes in $V = V_S \cup V_R$ also has a feature set $X_V$ which consists of the representation obtained during the construction of $E_{RR}$. If scvi was used to obtain a joint representation of $M_S$ and $M_R$ before building the KNN-graph, we recommend using the scvi representation as the feature set. Please see **Figure** 1 **A** for a detailed illustration of the graph construction step.

#### 2.3 Model architecture, training & prediction

##### 2.3.1 Labels and loss functions

For each reference node $v_r \in V_R$, we associate a set of labels $L = \{l_1, l_2, \ldots, l_k\}$, where $k$ is the number of labels for $v_r$. In These labels can be of different types: discrete, continuous, or hierarchical, each requiring a distinct approach for evaluation and error quantification. In principal, any number of labels can be used, but the current implementation have been tested with $L = \{l_1, l_2\}$, where $l_1$ corresponds to cell state labels and $l_2$ corresponds to discrete clonal labels.

For discrete labels, which represent categorical variables, we utilize the negative log-likelihood (NLL) as the evaluation criterion. The total loss function is a geometric mean of NLL los computed for cell state labels and for clone labels, i.e: $Loss = \sqrt{NLL(l_1, p_1) * NLL(l_2, p_2)}$, where $p_1$ and $p_2$ correspond to softmax-transformed logits for clone and cell type predictions.

##### 2.3.2 Model architecture, training & prediction

The GNN model demonstrated on **Figure** 1 B consists of several GAT layers that aim to create a consistent embedding space where nodes that are close in the graph structure tend to have similar embeddings. The model architecture consists of several GAT layers with skip-connections in between them, one embedding layer, and then $k$ ($k = 2$ with clone and cell state mapping) fully connected (FC) layers that are further used for predictions of the corresponding label $l_i$.

The supervision is performed only for reference nodes $V_R$ while the final prediction is done on $V_S$. This implementation represents a transfer learning approach based on the following assumptions:

1. **Shared Structure and Feature Space** Spatial nodes often share some structural similarities of feature space with reference nodes. By learning from the reference

nodes, the GNN can infer a feature representation that is relevant for the spatial nodes as well, despite the lack of direct training on them.

2. **Neighbourhood Aggregation** The GNN learns by aggregating information from a node's neighborhood. Even if spatial nodes do not have labels, their features contribute to the aggregated features of the labeled reference nodes during training, enhancing the feature representations of reference nodes.

To summarize, the model performs two explicit tasks: GAT layers create similar feature representations for both spatial and reference nodes (1), while the last FC layers learn the correct label based on the well-mixed embedding space (2). Once the model is trained, those representations are used to predict labels on spatial nodes (**Figure** 1 **C**).

##### 2.3.3 Model calibration

Model calibration is essential when interpreting the output of a softmax layer as the confidence of a predicted class. Traditional calibration techniques, such as Platt scaling[22] or isotonic regression[23], were not suitable for our case because the training data (reference data) and test data (spatial data) distributions differ significantly. To address these challenges, we leveraged the work by Chuan Guo et al.[24], which demonstrated that temperature scaling—a single-parameter variant of Platt scaling—is an effective method for model calibration.

Temperature scaling works by adjusting the confidence of the model's predictions through a temperature parameter ($T$), which scales the logits before applying the softmax function. This technique allows for recalibration of the predicted probabilities in a manner that is sensitive to the characteristics of the validation data.

Formally, given a vector of logits $\mathbf{z}$ from a neural network, temperature scaling produces calibrated probabilities $\mathbf{p}$ using the following transformation:

$$\mathbf{p} = \text{softmax}\left(\frac{\mathbf{z}}{T}\right)$$

where the softmax function is defined as:

$$\text{softmax}(\mathbf{z})_i = \frac{\exp(z_i/T)}{\sum_j \exp(z_j/T)}$$

Typically, the optimal temperature parameter is determined using a validation set. However, due to the out-of-distribution (OOD) nature of our problem, this approach was not feasible. Intuitively, the temperature parameter $T$ controls the entropy between classes: when $T = 1$, the calibration is equivalent to a normal softmax transformation, whereas $T > 1$ reduces the model's confidence in the dominant class. In our interpretation, $T$ represents the degree of class mixing for each spot. This concept is akin to methods like Cell2location[19] and Tangram[21], which also incorporate a similar parameter to account for the resolution of Visium spots.

Through experimentation, we found that a temperature setting of $T = 1.5$ provided the best correspondence with the results from Cell2location and validation using Xenium data (see section 2.5.5).

#### 2.4 Guidelines on training and hyperparameter settings

As demonstrated in our tutorials, the training procedure consists of the following steps:

- Reference nodes are separated into train and test sets with 70/30 proportions
- ‘NeigborLoader‘ is used on training nodes to perform neighbor sampling as described by Hamilton et al.[25]. This loader allows for mini-batch training of GNNs on large-scale graphs where full-batch training leads to out-of-memory problems. In our experience, the loader also drastically improved prediction robustness.
- The model is trained on train nodes and after each epoch train and validation losses are reported.
- The training is stopped when validation loss does not improve anymore for 10 epochs.

##### 2.4.1 Model adaptation as a function of technology

Although spaceTree was developed as a technology-agnostic approach, in this study focusing on GB, we applied spaceTree exclusively to Visium data. Consequently, all illustrations, including **Figure** 1, showcase Visium as a representative example.

In principle, spaceTree can be applied to any spatial dataset, as demonstrated in the package tutorials.

As described in section 2.5.3, the optimal hyperparameter settings for Visium were: $E_{SR} = E_{RR} = 10$, learning rate of 0.1, 50 hidden dimensions, 1 attention head, and a weight decay of 0.001. Additionally, we show that the validation loss reported by spaceTree is positively correlated with ground truth performance (**Figure** 2 **B**). Therefore, we recommend that users select hyperparameters that minimize the validation loss. **Figure** 2 **A** highlights that learning rate and hidden dimension size are the most critical parameters for achieving good performance. Thus, we suggest focusing primarily on tuning these parameters.

Our tutorials provide recommended starting points for other technologies, such as Xenium and Visium HD:

- **Xenium:** $E_{SR} = E_{RR} = 10$, learning rate of 0.01, 100 hidden dimensions, 2 attention heads, and a weight decay of 0.001.
- **Visium HD:** $E_{SR} = E_{RR} = 10$, learning rate of 0.001, 300 hidden dimensions, 1 attention head, and a weight decay of 0.01.

#### 2.5 Application to GB data

##### 2.5.1 Clone definition

Initially, scRNA-seq clones were defined using infercnvpy[6], and scATAC-seq clones were determined with epiAneufinder[26]. Joint clone assignment between scRNA-seq and scATAC-seq was performed by identifying cells that passed QC thresholds in both datasets.

We observed that scATAC-seq-based CNVs exhibited greater variability than RNA-based CNVs, likely due to the higher sensitivity of ATAC-seq in detecting open chromatin regions, resulting in a larger number of CNV events. To be more conservative and reduce false positives, we focused on prioritizing the RNA-seq data for clone assignment. Specifically, we matched clones from scATAC-seq to scRNA-seq clones, rather than the reverse.

To reduce clone redundancy, we assessed the similarity between scATAC-seq clones using correlation distance. Clones with pairwise distances below a predefined threshold were merged to eliminate highly similar clones.

To mitigate differences in resolution between modalities (250-gene windows for scRNA-seq and 1MB regions for scATAC-seq CNVs), we applied interpolation to the RNA-based CNV profiles, aligning them with the larger genomic windows of scATAC-seq data.

For matching clones across modalities, we computed two metrics: cell overlap and cosine similarity between cluster centroids. Cell overlap was defined as the proportion of cells shared between scRNA-seq and scATAC-seq clones, while cosine similarity was calculated based on the mean CNV profile for each clone. These metrics were averaged to create a composite similarity score.

Next, we retained the top 20% of clone matches based on this composite similarity score. For each RNA clone, we greedily selected the most similar scATAC-seq clones, assigning cells shared between matched RNA and ATAC clones to a new joint clone identifier.

Finally, we applied label spreading to propagate clone labels to unclassified RNA cells based on their CNV profiles. Each cell was assigned a probability of belonging to a clone, allowing flexibility in downstream analyses by using different confidence thresholds. For most analyses, we applied a confidence threshold of 0.9 to ensure high-confidence clone assignments.

##### 2.5.2 Preprocessing

To compute the initial graph, we used scvi-tools to obtain the initial embedding of spatial and reference data. scvi-tools are often used to get rid of batch effects but were not developed to address resolution differences between spatial and single-cell data. Nevertheless, for us, scvi-tools performed rather well for an initial embedding that was further improved with spaceTree. Before running scvi, both count matrices were normalized to a target sum of 10 000 and then log-transformed ($ln(x+1)$). Then scvi model was trained with 2 layers and n_latent=30. The obtained embedding space was used to compute a KNN graph for $E_{RR}$ and $E_{SR}$ and also used as node features.

For $E_{SS}$, we used the original Visium grid structure, connecting node $v_S^i$ to $v_S^j$ if it falls within a 5-by-5 neighborhood, with connection weights based on proximity. Specifically, nodes within a 1-unit distance (first-degree neighborhood) are assigned a weight of 1, while nodes within a 2-unit distance (second-degree neighborhood) are assigned a weight of 0.5.

##### 2.5.3 Hyperparameter Optimization and Benchmark

Hyperparameter optimization can be particularly challenging in cases where a validation set cannot be created due to the out-of-distribution nature of the problem. In these scenarios, traditional accuracy or loss metrics on the validation set may not directly reflect the model's performance on unseen data. To address this, we performed hyperparameter selection using hold-out validation data derived from part of the reference data. In **Section** 2.5.4, we further explore whether model accuracy is a reliable metric for hyperparameter selection.

The hyperparameters of spaceTree can be broadly categorized into two groups: those related to graph construction (e.g., neighborhood sizes in the KNN-graph for $E_{SR}$ and $E_{RR}$) and the standard hyperparameters of a GNN, such as learning rate, hidden dimension size, attention heads, and weight decay.

We evaluated spaceTree's performance on hold-out validation data for 1920 combinations of the following hyperparameters:

- Neighbours for $E_{SR}$ and $E_{RR}$ construction: [5, 10, 25, 50]
- Learning rate: $[0.1, 0.05, 0.01, 0.001, 0.0001]$
- Hidden dimensions: $[50, 100, 200, 300]$
- Attention heads: $[1, 2, 3]$
- Weight decay: $[0.01, 0.001, 0.0001]$

This evaluation was carried out across all samples from AT10 and AT15 patients, as these samples exhibited the most diversity in cell state representation (see **Figure 1** in the main text).

To assess the impact of each hyperparameter, we constructed a linear model where the features were different hyperparameter sets and the response was the model's accuracy. **Figure** 2 **B** highlights the importance of each hyperparameter, with the learning rate being the most significant factor.

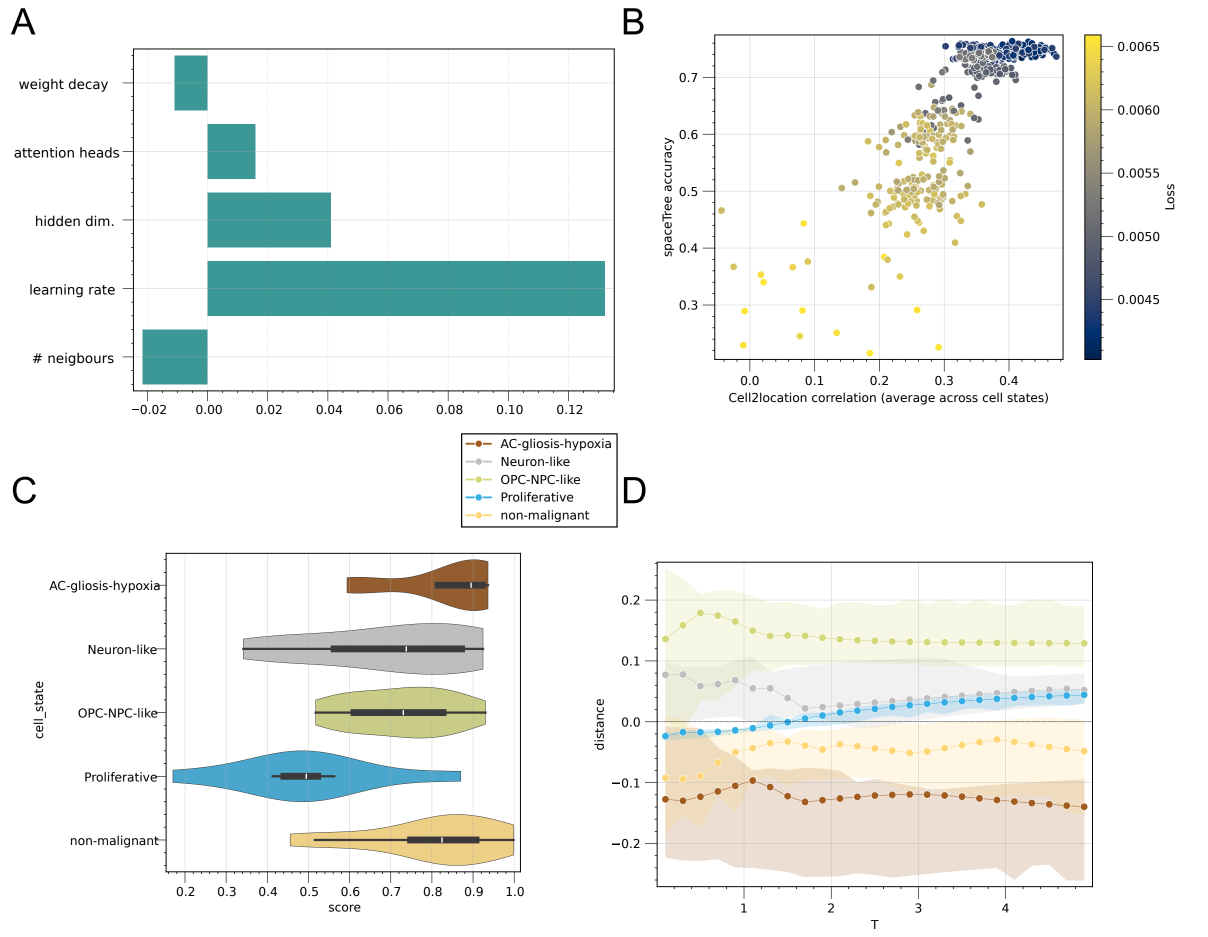


Figure 2: Hyperparameter selection and validation. A: Hyperparameter importance values for spaceTree accuracy prediction. B: Scatter plot illustrating the relationship between spaceTree's accuracy, model loss, and correlation to Cell2location (averaged across four cell states). C: Comparison of the optimal hyperparameter set to Cell2location results. D: Selection of the mixing parameter $T$ based on Euclidean distance to Cell2location results, with confidence intervals indicating variance across samples.

##### 2.5.4 Benchmark

To validate our hyperparameter choices and assess the overall performance of spaceTree, we compared its results to those obtained using the Cell2location model[19] for cell type mapping. Although Cell2location does not serve as a definitive ground truth, we considered it a reliable reference due to several factors. First, Cell2location leverages the statistical strength of multiple samples and patients, and given the extensive size of the dataset analyzed, we expect its performance to be near-optimal. While spaceTree can be applied across multiple samples, clonal mapping poses a unique challenge due to the non-overlapping clonal labels across patients.

The Cell2location results were further validated using the Xenium assay, showing high correspondence (see **Figure 3** in the main text). As there are no existing methods for direct comparison of clonal mapping, we hypothesize that robust cell type propagation will provide reasonable inference on clonal labels. This hypothesis was further validated using Laser Capture Microdissection (LCM) (see **Figure 5** main text).

To measure the correspondence between Cell2location and spaceTree mappings, we focused on four major malignant cell states: AC-gliosis-hypoxia, OPC-NPC-like, proliferative, and neuron-like cells. Cosine similarity between predicted cell state proportions was used to assess the degree of correspondence between spaceTree and Cell2location annotations. We found that spaceTree's validation loss and accuracy correlated well with the correspondence to Cell2location (**Figure** 2 **A**), demonstrating the reliability of these metrics in the absence of ground truth.

**Figure** 2 **C** presents the correspondence between spaceTree and Cell2location for the selected set of hyperparameters across all AT10 and AT15 samples.

###### 2.5.5 Temperature Selection

Cosine similarity was used as a metric in **Figure** 2 **C** because it allows us to disregard the temperature (mixing) parameter $T$ during the initial hyperparameter search, significantly reducing the search space. Since $T$ only affects the scale of class prevalence, it does not greatly impact cosine similarity between spaceTree and Cell2location results. Once the primary hyperparameters were selected, we fine-tuned $T$ to reflect appropriate mixing by minimizing the Euclidean distance between spaceTree and Cell2location results.

**Figure** 2 **D** shows that the mixing parameter affects different classes depending on their prevalence. Negative distances indicate that spaceTree underpredicts a certain cell state compared to Cell2location, while positive distances indicate overprediction. By minimizing the sum of absolute distances, we selected the optimal value of $T$ to be 1.5. The proximity of this value to 1 suggests that even without extensive calibration, spaceTree is not overly confident and produces results comparable to Cell2location.

#### 3. Conclusion

In this work, we present spaceTree, a novel framework for joint modeling of cell type and clonal heterogeneity in spatial omics data. Through the use of a multi-task graph neural network and label propagation, spaceTree integrates transcriptomic, spatial, and clonal information, providing a comprehensive view of tumor heterogeneity. Importantly, the utility of spaceTree was demonstrated in the main manuscript, where its effectiveness was shown in GB samples, capturing both spatially resolved cell types and clonal structures. The results were validated using Laser Capture Microdissection (LCM), further confirming the robustness of the method. This framework is adaptable to a wide range of technologies, offering a powerful tool for understanding the intricate relationships between cellular composition and spatial organization within tumors. The ability to simultaneously map both cell types and clones while accounting for spatial context represents a significant advancement in spatial omics analysis, especially in complex cancer landscapes like GB.
